# Opportunity Drives Spillover: Serological Surveillance across Carnivores, Omnivores and Herbivores in an HPAIV H5 Hotspot in North-East Germany, 2023–2025

**DOI:** 10.1101/2025.09.30.678011

**Authors:** Anne Günther, Josefine Wassermann, Jonas Heck, Marin Bussi, Andrea Aebischer, Christoph Staubach, Hannes Bergmann, Fabian H. Leendertz, Martin Beer, Gereon Schares, Kerstin Wernike

**Affiliations:** Friedrich-Loeffler-Institut, Greifswald - Insel Riems, Germany; Utrecht University, The Netherlands; Helmholtz Institute for One Health, Greifswald, & University of Greifswald, Germany

## Abstract

In North-East Germany’s offshore islands and mainland coast, wild ruminants, boar, and carnivores were tested for H5-HPAI antibodies. Wild ruminants were seronegative; 3.5% of boar and 12.5–21.9% of carnivores were seropositive, evidencing frequent spillover. Because such events may accelerate mammalian—and ultimately human—adaptation, sustained One-Health monitoring is essential.

**Summary:** Across offshore islands and the mainland coast of North-East Germany, H5 HPAI serosurveillance in wildlife found no seroconversion in ruminants, but positives in 3.5% of boar and 12.5–21.9% of carnivores, indicating ongoing spillover and risk of avian-to-mammalian adaptation.

## Background

The shift of seasonal epizootics towards a perpetual enzootic in Europe became a milestone in the epidemiology of avian influenza viruses (AIV) of high pathogenicity (HPAIV) (1). Rooted in a common ancestral virus of the goose-Guangdong lineage in Southeast Asia, the evolving diversity of subtype H5 genotypes in Europe paved the way for the ongoing panzootic, globally threatening domestic and wild birds alike, and resulting in numerous mammalian spillover infections world-wide. A broad and increasing spectrum of terrestrial, semiaquatic and marine/aquatic mammalian species is affected (2). Despite the recent developments of mastitis in domestic cattle via ascending udder infections with HPAIV H5 (3), the more constant interface over the years was direct, often alimentary, exposure to HPAIV H5-positive prey or food (4).

Infections in predatory or scavenging species frequently caused neurological signs, including severe encephalitis as cause of death (5). The concern about increasing chances for spillover events from birds to mammals proved to be justified already in the early stage of enzootic HPAIV H5 in Europe. However, those studies also suggested a certain level of asymptomatic infections and the possibility of surviving exposure (6).

Here, we compared HPAIV H5 antibody prevalences between groups with frequent contact opportunities (carnivores) and conceivable or unlikely contact opportunities (omnivores, herbivores). We identify frequently affected host groups and factors ultimately favor the risk for carnivores considered susceptible.

### The Study

Between December 2023 and February 2025, we collected samples from 644 hunted predator game (group *I*) and 343 hoofed game animals (groups *II* and *III*) in the context of an ongoing disease surveillance project in the federal state of Mecklenburg-Western Pomerania (MWP), North-East Germany. In particular, the offshore islands and the mainland coast form important avian migratory areas, but also breeding habitats with a high overall species diversity and abundance of wild birds. Many of these species have been affected during various HPAI H5-epizootics and the recent enzootic (7).

Predator game comprised five species of the taxonomic families *Canidae, Procyonidae* and *Mustelidae* (Table). *Cervidae, Bovidae* and *Suidae* form groups *II* and *III* with five different species in total (one sampled individual remains unspecified). Nasal swabs and, for group *I*, lung and brain samples were screened by RT-qPCR for influenza A virus (IAV) RNA (n=980) (Technical Annex, Supplementary Table 1) that all tested negative.

By nucleoprotein (NP)-based ELISA, no seroconversion against IAV could be detected in wild ruminants, but 5.2% (95% confidence interval (95%CI): 1.9-11.0%) of the wild boar (*Sus scrofa*) tested positive and all carnivorous species, except the single tested European pine marten (*Martes martes*) (Table). The subsequent assay for detection of H5-specific antibodies revealed 3.5% (0.9-8.7%) seropositive wild boar and between 12.5% (0.3-52.7%) and 21.9% (12.5-34.0%) seropositive carnivorous species (Table; Technical Annex, Supplementary Table 1), thereby confirming the vast majority of seroconversion caused by an IAV of subtype H5. The range of non-H5 IAV antibody-positive individuals varied between 4.7% (0.9- 13.1%) in raccoons (*Procyon lotor*) and 4% (2.4-6.9%) in (red) foxes (*Vulpes sp*.*/vulpes*) to 1.7% (0.2-6.1%) in wild boar and 1.1% (0.1-3.9%) in raccoon dogs (*Nyctereutes procyonoides*). A subset of NP-antibody positive or negative samples has been additionally tested in a serum neutralization test (SNT) to confirm the presence or absence of antibodies against subtype H5 (Technical Annex). Exemplary, we confirmed ELISA-findings for eight IAV-positive, but H5-negative, carnivorous samples, compared to four H5-positives, while the level of H5-seroconversion in the wild boar samples remained below detection limit in the SNT.

Wild ruminants represent mammalian species with rather unlikely contact possibilities to potentially infected wild birds and, as herbivores, no apparent interface in their diet. However, direct exposure cannot be ruled out, and low susceptibility may be another explanation for our consistent seronegative findings. At the other end of the food chain, our serological results suggest previous exposure of carnivores to subtype H5 viruses. Raccoons, foxes and raccoon dogs represent the species with the highest proportion of seroconversion (Table). Despite similar host species tested, our results only partially confirm observations of previous studies, where mainly stone martens and foxes were found to be seropositive in about one third of the animals tested (6). This discrepancy indicates the relevance of other factors than host susceptibility and host occurrence.

Univariable regression analysis (Figure) revealed that foxes from the Island Rügen collected close to the bay coast, close to the Baltic sea or close to watercourses had an increased risk for H5-specific antibodies (P < 0.05; Technical Annex, Supplementary Table 2 and 3). Seropositivity increased with age; 23.5% (43/183) of the adult and 11.6% (10/86) of the juvenile were positive for H5-specific antibodies. A final multivariable model (Technical Annex, Supplementary Table 4) selected by stepwise forward-backward selection which included also age revealed that with increasing distance to “watercourses” (including flowing waters, water source areas, streams, ditches, rivers and canals) the risk of foxes being positive for H5-specific antibodies decreased (P < 0.0001) while with increasing distance to “shrubland” (including hedges, bushes, field crops, group of trees, rows of trees, avenues, dominant single trees and group of shrubs) the risk increased (P < 0.005). In contrast, high proportions of “shrubland” in a 2.5 km buffer zone around the sampling site of foxes represented a protective factor (P < 0.005), allowing for the assumption of less exposure to HPAIV via water-associated hosts of which e.g. *Anseriformes* are known to graze on open lands.

**Figure 1.**
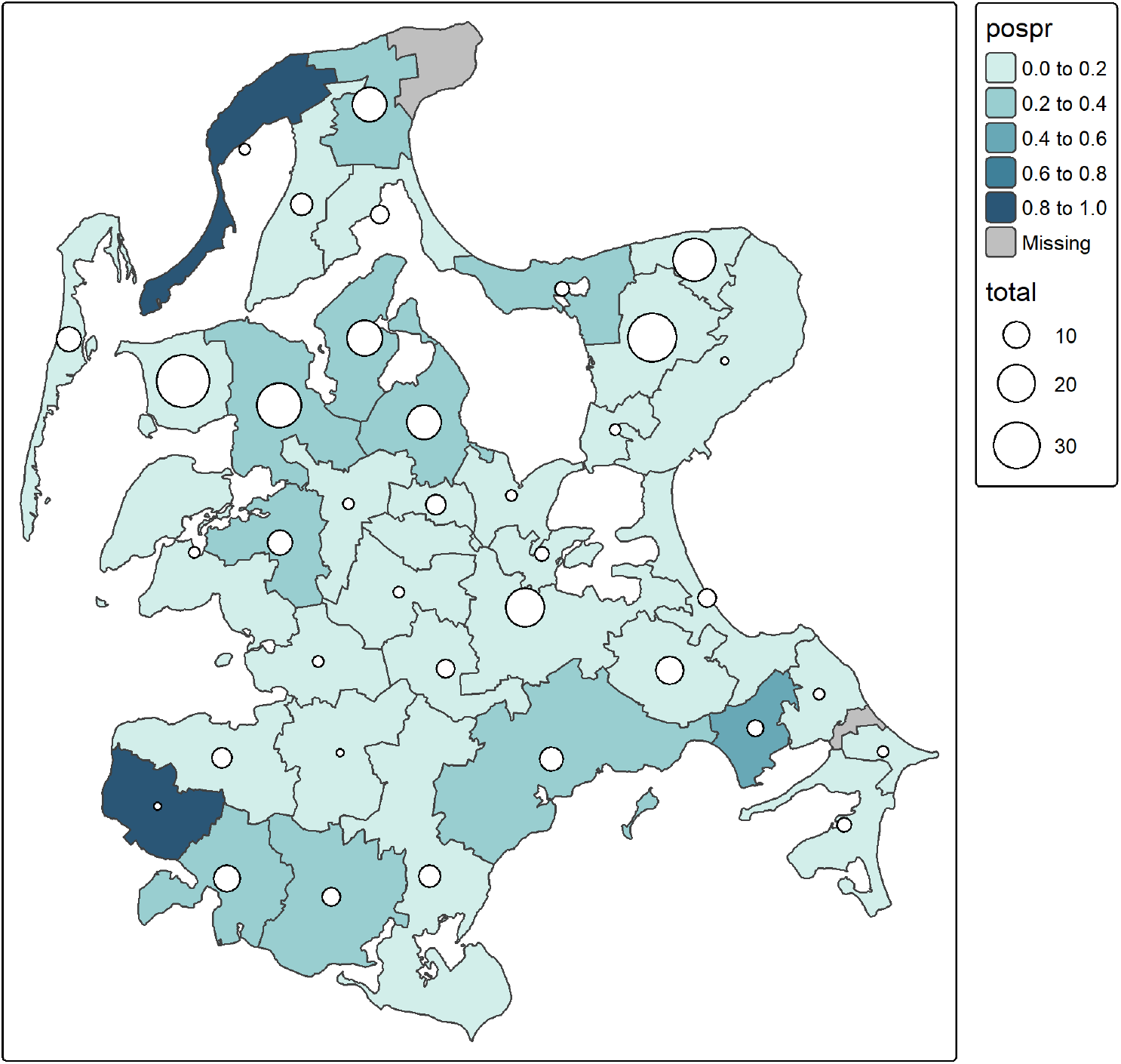
Proportions (pospr) of juvenile and adult foxes with H5-specific antibodies in various municipalities of the Island Rügen. White circles represent (total) numbers of sampled and analyzed foxes per municipality.

The possible existence of a similar correlation for omnivorous species might serve as explanation for the 3.5% (0.9-8.7%) seropositivity in wild boar. In addition to their scavenging or predatory behavior (8), wild boars have been described as nest robbers for waterfowl species in other European wetland areas (9). All individuals with H5-specific antibodies belonged to sounders roaming in water-associated zones (the Island Rügen and Fischland-Darß-Zingst; Technical Annex, Supplementary Table 1), where a higher likelihood of interaction with e.g. waterfowl species can be expected.

After the previous HPAIV H5N8 epizootic in 2016/2017, a broad serological investigation in wild boars has been conducted in Southern Germany (10). Low-level prevalence of H5N8 neutralizing antibodies (1.13%) have been detected, with a single positive case occurring close to a prior HPAI outbreak in waterfowl (10, 11). While in domestic pigs the overall susceptibility in experimental settings remained low despite a high infectious dose of a single 2.3.4.4b strain (12), it cannot be excluded, that the general susceptibility may be varying between different HPAIV H5 genotypes in *Suidae*.

Although little is known about the persistence of AIV antibodies in wild mammals, serological surveillance may provide a more complete picture of past exposure to potentially infected avian (reservoir) hosts than strictly molecular testing. The discrepancy in NP- and H5-antibody detection leaves other AIV subtypes contemplable for initial seroconversion, so a combination of testing approaches seems advisable.

## Conclusions

Our findings of H5-specific antibodies in carnivores and wild *Suidae* indicate an ongoing level of spillover events, especially in wetland-habitats potentially shared with main (reservoir) hosts (*Anseriformes, Charadriiformes)*. The role of wild boars requires further monitoring, as *Suidae* are classified potential mixing vessels for swine, human and avian derived influenza A viruses (13). We recommend continuous surveillance for a broad scale of wildlife species, to estimate the proportion of H5-infections in *Mammalia*, as every such infection provides a chance for mammalian adaptations, eventually also to humans.

## Supporting information

Technical Annex

## Acknowledgments

We thank all hunters contributing to this study especially the hunters organized by the “Jagdverband Rügen-Hiddensee”, Bianka Hillmann, Julien Schäfer, Alrik- Markis Kunisch, Martina Abs, Kristin Trippler and Diana Parlow for excellent technical assistance, and Ralf Redmer, Thomas Pieper, Marco Beerbohm, Sophia Ziegler, Timm Harder and Ronja Piesche for the help during dissection and laboratory diagnostics. Many thanks to the federal state of Mecklenburg-Western Pomerania and to the “Bundesamt für Kartografie und Geodäsie” for providing data (“Biotoptypen- und Nutzungstypenkartierung” (BNTK) and “Digitale Basis-Landschaftsmodell” (DLM), respectively). The study is part of the One Health project Wildlife disease monitoring in Mecklenburg-Western Pomerania through an integrated One Health surveillance-response system (WiMoPOH) funded by the “Initialisierungs- und Vernetzungsfonds für Infektionsforschung” managed by the Helmholtz Institute for One Health. The study received additional funding by the European Union under grant agreement (101084171) - (Kappa-Flu). Views and opinions expressed are however those of the author(s) only and do not necessarily reflect those of the European Union or REA. Neither the European Union nor the granting authority can be held responsible for them.

## Conflict of Interest

The authors declare that the research was conducted in the absence of any commercial or financial relationships that could be construed as a potential conflict of interest.

**Table 1.**
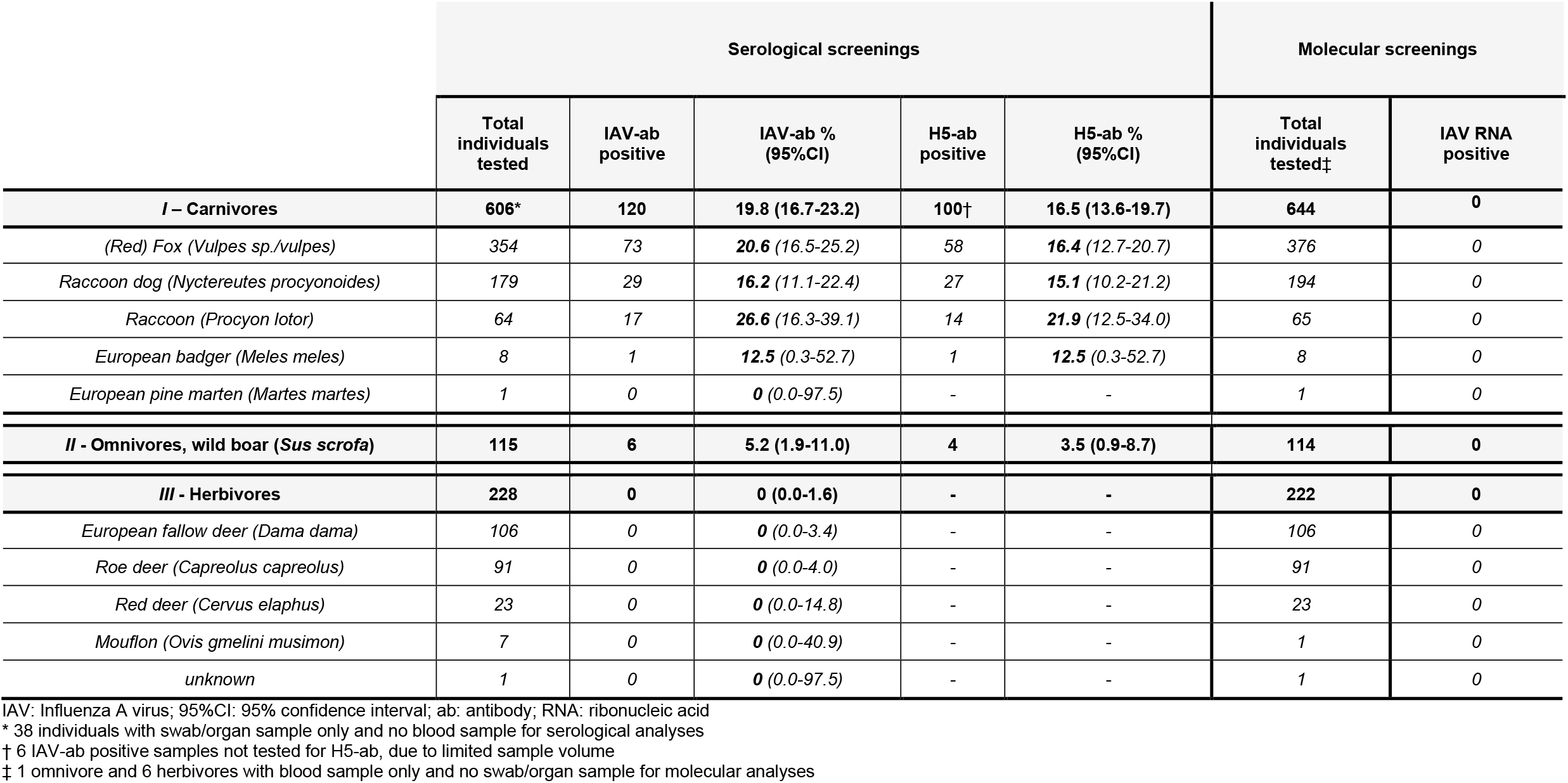
Overview on all species sampled for serological and molecular screenings grouped according to their feeding behavior: (I) carnivorous, (II) omnivorous and (III) herbivorous mammals. The results refer to screenings on generic antibodies against influenza A virus or specific for subtype H5 based on two subsequent multispecies enzyme-linked immunosorbent assays, including the 95% confidence interval. Reverse transcription quantitative real-time polymerase chain reaction results are presented for swab/organ samples tested for influenza A virus ribonucleic acid.

|  | Serological screenings |  |  |  |  | Molecular screenings |  |
| --- | --- | --- | --- | --- | --- | --- | --- |
|  | Total individuals tested | IAV-ab positive | IAV-ab % (95%CI) | H5-ab positive | H5-ab % (95%CI) | Total individuals tested‡ | IAV RNA positive |
| **I – Carnivores** | **606\*** | **120** | **19.8 (16.7-23.2)** | **100†** | **16.5 (13.6-19.7)** | **644** | **0** |
| (Red) Fox (*Vulpes sp./vulpes*) | 354 | 73 | 20.6 (16.5-25.2) | 58 | 16.4 (12.7-20.7) | 376 | 0 |
| Raccoon dog (*Nyctereutes procyonoides*) | 179 | 29 | 16.2 (11.1-22.4) | 27 | 15.1 (10.2-21.2) | 194 | 0 |
| Raccoon (*Procyon lotor*) | 64 | 17 | 26.6 (16.3-39.1) | 14 | 21.9 (12.5-34.0) | 65 | 0 |
| European badger (*Meles meles*) | 8 | 1 | 12.5 (0.3-52.7) | 1 | 12.5 (0.3-52.7) | 8 | 0 |
| European pine marten (*Martes martes*) | 1 | 0 | 0 (0.0-97.5) | - | - | 1 | 0 |
| **II - Omnivores, wild boar (*Sus scrofa*)** | **115** | **6** | **5.2 (1.9-11.0)** | **4** | **3.5 (0.9-8.7)** | **114** | **0** |
| **III - Herbivores** | **228** | **0** | **0 (0.0-1.6)** | **-** | **-** | **222** | **0** |
| European fallow deer (*Dama dama*) | 106 | 0 | 0 (0.0-3.4) | - | - | 106 | 0 |
| Roe deer (*Capreolus capreolus*) | 91 | 0 | 0 (0.0-4.0) | - | - | 91 | 0 |
| Red deer (*Cervus elaphus*) | 23 | 0 | 0 (0.0-14.8) | - | - | 23 | 0 |
| Mouflon (*Ovis gmelini musimon*) | 7 | 0 | 0 (0.0-40.9) | - | - | 1 | 0 |
| unknown | 1 | 0 | 0 (0.0-97.5) | - | - | 1 | 0 |

IAV: Influenza A virus; 95%CI: 95% confidence interval; ab: antibody; RNA: ribonucleic acid
\* 38 individuals with swab/organ sample only and no blood sample for serological analyses
† 6 IAV-ab positive samples not tested for H5-ab, due to limited sample volume
‡ 1 omnivore and 6 herbivores with blood sample only and no swab/organ sample for molecular analyses

## References

1. Pohlmann A, King J, Fusaro A, Zecchin B, Banyard AC, Brown IH, et al. Has Epizootic Become Enzootic? Evidence for a Fundamental Change in the Infection Dynamics of Highly Pathogenic Avian Influenza in Europe, 2021. mBio. 2022;13(4). Epub 2022/06/22. doi: 10.1128/mbio.00609-22. PubMed PMID: 35726917.

2. Plaza PI, Gamarra-Toledo V, Eugui JR, Lambertucci SA. Recent Changes in Patterns of Mammal Infection with Highly Pathogenic Avian Influenza A(H5N1) Virus Worldwide. Emerg Infect Dis. 2024;30(3):444–52. doi: 10.3201/eid3003.231098. PubMed PMID: 38407173; PubMed Central PMCID: PMC10902543.

3. Butt SL, Nooruzzaman M, Covaleda LM, Diel DG. Hot topic: Influenza A H5N1 virus exhibits a broad host range, including dairy cows. JDS Commun. 2024;5(Suppl 1):S13–S9. Epub 20240930. doi: 10.3168/jdsc.2024-0638. PubMed PMID: 39429893; PubMed Central PMCID: PMC11489455.

4. Alkie TN, Cox S, Embury-Hyatt C, Stevens B, Pople N, Pybus MJ, et al. Characterization of neurotropic HPAI H5N1 viruses with novel genome constellations and mammalian adaptive mutations in free-living mesocarnivores in Canada. Emerging microbes & infections. 2023;12(1). Epub 2023/03/08. doi: 10.1080/22221751.2023.2186608. PubMed PMID: 36880345; PubMed Central PMCID: PMC10026807.

5. Bordes L, Vreman S, Heutink R, Roose M, Venema S, Pritz-Verschuren SBE, et al. Highly Pathogenic Avian Influenza H5N1 Virus Infections in Wild Red Foxes (Vulpes vulpes) Show Neurotropism and Adaptive Virus Mutations. Microbiology spectrum. 2023;11(1). Epub 2023/01/24. doi: 10.1128/spectrum.02867-22. PubMed PMID: 36688676; PubMed Central PMCID: PMC9927208.

6. Chestakova IV, van der Linden A, Martin BB, Caliendo V, Vuong O, Thewessen S, et al. High number of HPAI H5 Virus Infections and Antibodies in Wild Carnivores in the Netherlands, 2020-2022. bioRxiv. 2023. doi: 10.1101/2023.05.12.540493.

7. European Food Safety A, Abrahantes JC, Aznar I, Catalin I, Kohnle L, Mulligan KF, et al. Avian influenza annual report 2023. EFSA J. 2025;23(1):e9197. Epub 20250122. doi: 10.2903/j.efsa.2025.9197. PubMed PMID: 39844828; PubMed Central PMCID: PMC11751681.

8. Barrios-Garcia MN, Ballari SA. Impact of wild boar (Sus scrofa) in its introduced and native range: a review. Biol Invasions. 2012;14(11):2283–300. doi: 10.1007/s10530-012-0229-6. PubMed PMID: WOS:000309856500008.

9. Barasona JA, Carpio A, Boadella M, Gortazar C, Piñeiro X, Zumalacárregui C, et al. Expansion of native wild boar populations is a new threat for semi-arid wetland areas. Ecol Indic. 2021;125. doi: 10.1016/j.ecolind.2021.107563. PubMed PMID: WOS:000637200300002.

10. Schülein A, Ritzmann M, Christian J, Schneider K, Neubauer-Juric A. Exposure of wild boar to Influenza A viruses in Bavaria: Analysis of seroprevalences and antibody subtype specificity before and after the panzootic of highly pathogenic avian influenza viruses A (H5N8). Zoonoses Public Hlth. 2021;68(5):503–15. doi: 10.1111/zph.12841. PubMed PMID: WOS:000649976100001.

11. Globig A, Staubach C, Sauter-Louis C, Dietze K, Homeier-Bachmann T, Probst C, et al. Highly Pathogenic Avian Influenza H5N8 Clade 2.3.4.4b in Germany in 2016/2017. Frontiers in veterinary science. 2018;4. doi: 10.3389/fvets.2017.00240. PubMed PMID: WOS:000451814500002.

12. Graaf A, Piesche R, Sehl-Ewert J, Grund C, Pohlmann A, Beer M, et al. Low Susceptibility of Pigs against Experimental Infection with HPAI Virus H5N1 Clade 2.3.4.4b. Emerg Infect Dis. 2023;29(7):1492–5. doi: 10.3201/eid2907.230296. PubMed PMID: 37347930; PubMed Central PMCID: PMC10310384.

13. Abdelwhab EM, Mettenleiter TC. Zoonotic Animal Influenza Virus and Potential Mixing Vessel Hosts. Viruses. 2023;15(4). Epub 2023/04/28. doi: 10.3390/v15040980. PubMed PMID: 37112960; PubMed Central PMCID: PMC10145017.

