## Supplementary material for "Opportunity Drives Spillover: Serological Surveillance across Carnivores, Omnivores and Herbivores in an HPAIV H5 Hotspot in North-East Germany, 2023–2025": Technical Annex

### 1. Sample collection

Samples have been collected during hunts in the field (swab samples from ruminants and wild boar) or during necropsies performed at the Friedrich-Loeffler-Institut (FLI) after a freeze-thaw cycle as described in Kunisch, et al. (1) (swabs and organ samples from carnivores). The number of sampled wildlife animals as well as their origin are available in the Supplementary Table 1. One dry swab per individual was taken from both nostrils of predator and hoofed game species and immediately has been stored in 2 mL Minimum Essential Medium (MEM). Brain and lung samples from carnivorous species were collected during necropsy and transferred into tubes, prepared with 1.5 mL MEM and a 5 mm steal bead for homogenization in a TissueLyser (Qiagen, Hilden, Germany).

Samples for serological analyses were collected in form of whole blood or serum samples taken directly after the hunt out of the exit wounds or from the body cavity of thawed carcasses during necropsy.

The collected samples were stored at -20 °C until analyses.

### 2. Ethics Approval

All samples have been retrieved from animals that were hunted by local hunters according to the appropriate German legislation or that were found dead. No ethical approval is required for this kind of sample collection in the federal state of Mecklenburg-Western Pomerania, Germany.

### 3. Laboratory Analyses

Swab sample fluid and organ homogenate supernatant (100 µL) served as input for bead-based extraction of nucleic acids on a King Fisher 96 Flex (Thermo Fisher Scientific, Braunschweig, Germany) with the NucleoMag® VET kit (Macherey-Nagel, Düren, Germany) following the manufacturer’s instructions.

The nucleic acid was subsequently tested by an RT-qPCR targeting influenza A virus (IAV) matrix (M) gene as described by Hoffmann, et al. (2) in combination with an internal control system based on the beta-actin gene (3).

The serum or plasma-like supernatant was applied to two consecutive commercial enzyme-linked immunosorbent assays (ELISA) according to the manufacturer's instructions. However, for the carnivores, the 1:10 dilution of the sample was chosen to account for possible initial dilution due to the nature of the sample collection. The ID Screen® Influenza A Antibody Competition Multi-species (Innovative Diagnostics, Grabels, France) allows for the detection of antibodies against the nucleoprotein (NP) of IAV. In case of positive or questionable results, those samples have been tested in the ID Screen® Influenza H5 Antibody Competition ELISA (Innovative Diagnostics, Grabels, France) to confirm the presence or absence of antibodies specific against hemagglutinin of subtype H5.

To confirm the ELISA results, samples have been applied to an in-house serum neutralization assay (SNT). In preparation, blood samples have been heat inactivated at 56 °C for 30 minutes. Due to the highly hemolytic nature of the blood samples, the sample suitability for heat inactivation was severely compromised and only allowed for a very limited subset finally being tested via SNT (carnivore-panel n=12; wild boar samples n=6). We followed the procedure described by Hennig, et al. (4) with the following modifications: A trypsin-independent and highly pathogenic avian influenza virus was applied (HPAIV H5 strain A/chicken/Ger-N1/4286/2022) for the incubation, before a suspension of MDCK II cells (no. 0606; Collection of Cell Lines in Veterinary Medicine (CCLV), FLI, Greifswald - Insel Riems, Germany) in cell culture medium ($2x10^5$/mL) was added to each well.

The applied negative and positive control sera and the additional virus back-titration were tested in quadruplicates, so were the samples with few exceptions that lacked sufficient volume. The dilution cell culture medium for all steps was stocked with Baytril® (Enrofloxacine; 2.5 µg/mL; Elanco GmbH, Cuxhaven, Germany) and 5% fetal calf serum.

### 4. <u>Spatial analysis</u>

<u>Methods and data used for the determination of landcover proportions and distance measurements</u>

Statistical spatial analysis largely followed a procedure previously described (5). Coordinates of each animal capture site, spatial data on land cover in the entire area of interest and data on individual animals (e.g. age) were statistically assessed. A circular buffer zone of 2.5 km radius around the location where each animal carcass was found was used to approximates the home range of individual animals. The proportion of each landcover type in this buffer zone was calculated. Further, the distance of each animal location to the nearest polygon of a certain landcover type was calculated. Proportions, distances and individual animal data were used in logistic regression to model H5-specific seropositivity.

Data on land cover was retrieved from “Biotoptypen- und Nutzungstypenkartierung” (BNTK) provided by the federal state of Mecklenburg-Western Pomerania (last updated in 2012). Because the urban landscape of the Island Rügen (study area for foxes, racoon dogs and racoons) has expanded, replacing natural areas during the last two decades, BNTK data was combined with data from the "Digitale Basis-Landschaftsmodell" (DLM) provided by the "Bundesamt für Kartografie und Geodäsie", increasing accuracy and better reflecting the actual situation.

Methods used for spatial analyses and other statistical tests

To identify environmental risk factors for H5-specific seropositvity, the proportion of each land cover type falling in the respective buffer zone for each animal was calculated using the sf package in R. In R, the distance of each animal to its nearest landcover of the specified landcover types was calculated using the st_nn function. Each of the distances was scaled to 100 m steps and buffer proportions to 10 % steps.

Differences in H5-specific seropositivity by sex and age were tested using Chi-square tests. The number of foxes, racoon dogs and racoons submitted and the proportion of animals testing positive were tabulated by the municipality and visualized in a choropleth map. Municipality boundaries were chosen for visualization purposes. As hunters are submitting animals voluntarily, the distribution of animals submitted per hunter was skewed, with a median of three animals per hunter, but as many as 80 or as few as one animal.

Univariable logistic regression was conducted for all distance and buffer variables of each landscape type, and if the P value was $< 0.2$, the variable was included in a multivariable model. This threshold was established given the large number of variables in this dataset, aiming to only include those with potential relevance for H5 influenza infection in the multivariable model. The multivariable model was reduced by using forward-backwards selection in R, and the model with the lowest Akaike's Information Criterion (AIC; Akaike, 1974) is reported as final model. Additionally, the principle of Occam's razor was used when selecting the final variables, which states that when the difference in AIC between including a variable in a model is $\leq 2$ compared to a model without it, the simpler model (meaning without the variable in question) is the better fit. A weighted Moran's I and Geary's C were run on the standardized deviance residuals of each univariable logistic regression model to check for underlying spatial autocorrelation. If following these tests, autocorrelation would be apparent, a spatial logistic regression model would be used that allows for accounting for the spatial clustering present (Rogoll et al., 2024). The goodness of fit of the reduced multivariable model was evaluated using the omnibus test and pseudo $R^2$.

Results of analysis

Results in foxes: Foxes represent the most frequently sampled carnivorous species with background information on their location of origin available. Regression analysis using environmental variables (i.e. either proportions in a buffer zone of 2.5 km radius or nearest distances to a particular environmental characteristic; Supplementary Table 2) revealed a number of statistically significant associations, i.e. one at the level of $P < 0.01$

(Balticsea_dist100) and five at the level of $P \geq 0.01$ and $P < 0.05$ (Shrubland_prop, Urban_prop, Watercourses_prop, Baycoast_dist100, Watercourses_dist100). For the remaining variables displayed in Supplementary Table 3 values of $P \geq 0.05$ and $P <0.2$ had been calculated. All of these variables were included into a forward-backward selection process to identify the optimal final multivariable model. The initial analysis of putative associations of H5-specific antibodies with sex or age revealed that in adult foxes there was a statistically higher proportion of H5-specific seropositivity as compared to juvenile foxes Therefore, this variable was also included into the modelling to not disregard potential effects of age.

Results in raccoon dogs and racoons: Less data points were available for these animal species and same analyses as for foxes were performed. However, in this dataset, spatial autocorrelation was present, mandating the use of different analysis methods moving forward. Thus, results of these analyses are not displayed.

### 5. References

1. Kunisch AM, Wassermann J, Peters M, Feldmann K, Prinzenberg E, Klink JC, et al. In situ collected small intestinal mucosal swabs as alternative analytes to diagnose Echinococcus multilocularis infection in its definitive hosts by real-time polymerase chain reaction. Vet Parasitol. 2025;335:110418. Epub 2025/02/20. doi: 10.1016/j.vetpar.2025.110418. PubMed PMID: 39970833.

2. Hoffmann B, Harder T, Lange E, Kalthoff D, Reimann I, Grund C, et al. New real-time reverse transcriptase polymerase chain reactions facilitate detection and differentiation of novel A/H1N1 influenza virus in porcine and human samples. Berl Munch Tierarztl Wochenschr. 2010;123(7-8):286-92. Epub 2010/08/10. doi: 10.2376/0005-9366-123-286. PubMed PMID: 20690540.

3. Toussaint JF, Sailleau C, Breard E, Zientara S, De Clercq K. Bluetongue virus detection by two real-time RT-qPCRs targeting two different genomic segments. J Virol Methods. 2007;140(1-2):115-23. Epub 2007/01/02. doi: 10.1016/j.jviromet.2006.11.007. PubMed PMID: 17196266.

4. Hennig C, Graaf-Rau A, Schmies K, Elling R, Henneke P, Dürrwald R, et al. *Preprint*: High Serological Barriers May Contribute to Restricted Influenza-a-Virus Transmission between Pigs and Humans. 2025. doi: 10.2139/ssrn.5144941.

5. Rogoll L, Schulz K, Staubach C, Oļševskis E, Seržants M, Lamberga K, et al. Identification of predilection sites for wild boar carcass search based on spatial analysis of Latvian ASF surveillance data. Sci Rep. 2024;14(1):382. Epub 20240103. doi: 10.1038/s41598-023-50477-7. PubMed PMID: 38172492; PubMed Central PMCID: PMC10764341.

***Supplementary Table 1*** *Overview on metadata for the analyzed samples, including sampling ID, group designation, (scientific) species name and sample origin. The results of enzyme linked immunosorbent assay (ELISA) approaches targeting influenza A virus' (IAV) nucleoprotein (NP) or hemagglutinin of subtype H5 and the reverse transcription quantitative real-time polymerase chain reaction (RT-qPCR) screening results for IAV ribonucleic acid (RNA) are presented, if available. For samples, not tested in a specific test, the respective columns are filled with "-".*

| **Sample ID** | **Group** | **Species** | **Scientific species name** | **Sample origin rural district** | **Result IAV-NP antibody ELISA in percent** | **Assessment IAV-NP antibody ELISA** | **Result IAV-H5 antibody ELISA in percent** | **Assessment IAV-H5 antibody ELISA** | **Titer serum neutralization assay** | **IAV RT-qPCR nasal swab** | **IAV RT-qPCR lung** | **IAV RT-qPCR brain** |
|---|---|---|---|---|---|---|---|---|---|---|---|---|
| 2024BVD16887 | Herbivore | European fallow deer | *Dama dama* | Vorpommern-Rügen | 87 | negative | - | - | - | negative | - | - |
| 2024BVD16889 | Herbivore | European fallow deer | *Dama dama* | Vorpommern-Rügen | 107 | negative | - | - | - | negative | - | - |
| 2024BVD16890 | Herbivore | European fallow deer | *Dama dama* | Vorpommern-Rügen | 90 | negative | - | - | - | negative | - | - |
| 2024BVD16891 | Herbivore | European fallow deer | *Dama dama* | Vorpommern-Rügen | 123 | negative | - | - | - | negative | - | - |
| 2024BVD16892 | Herbivore | European fallow deer | *Dama dama* | Vorpommern-Rügen | 116 | negative | - | - | - | negative | - | - |
| 2024BVD16893 | Herbivore | European fallow deer | *Dama dama* | Vorpommern-Rügen | 119 | negative | - | - | - | negative | - | - |
| 2024BVD16894 | Omnivore | Wild boar | *Sus scrofa* | Vorpommern-Rügen | 115 | negative | - | - | - | negative | - | - |
| 2024BVD16895 | Herbivore | European fallow deer | *Dama dama* | Vorpommern-Rügen | 113 | negative | - | - | - | negative | - | - |
| 2024BVD16896 | Herbivore | European fallow deer | *Dama dama* | Vorpommern-Rügen | 117 | negative | - | - | - | negative | - | - |

| | | | | | | | | | | | | |
|---|---|---|---|---|---|---|---|---|---|---|---|---|
| 2024BVD16897 | Herbivore | European fallow deer | *Dama dama* | Vorpommern-Rügen | 121 | negative | - | - | - | negative | - | - |
| 2024BVD16898 | Herbivore | European fallow deer | *Dama dama* | Vorpommern-Rügen | 125 | negative | - | - | - | negative | - | - |
| 2024BVD16899 | Herbivore | Roe deer | *Capreolus capreolus* | Vorpommern-Rügen | 110 | negative | - | - | - | negative | - | - |
| 2024BVD16901 | Herbivore | Roe deer | *Capreolus capreolus* | Vorpommern-Rügen | 117 | negative | - | - | - | negative | - | - |
| 2024BVD16902 | Herbivore | European fallow deer | *Dama dama* | Vorpommern-Rügen | 111 | negative | - | - | - | negative | - | - |
| 2024BVD16903 | Omnivore | Wild boar | *Sus scrofa* | Vorpommern-Rügen | 106 | negative | - | - | - | negative | - | - |
| 2024BVD16904 | Herbivore | European fallow deer | *Dama dama* | Vorpommern-Rügen | 112 | negative | - | - | - | negative | - | - |
| 2024BVD16905 | Herbivore | European fallow deer | *Dama dama* | Vorpommern-Rügen | 78 | negative | - | - | - | negative | - | - |
| 2024BVD16906 | Herbivore | European fallow deer | *Dama dama* | Vorpommern-Rügen | 100 | negative | - | - | - | negative | - | - |
| 2024BVD16907 | Herbivore | European fallow deer | *Dama dama* | Vorpommern-Rügen | 111 | negative | - | - | - | negative | - | - |
| 2024BVD16908 | Herbivore | European fallow deer | *Dama dama* | Vorpommern-Rügen | 122 | negative | - | - | - | negative | - | - |
| 2024BVD16909 | Herbivore | European fallow deer | *Dama dama* | Vorpommern-Rügen | 106 | negative | - | - | - | negative | - | - |
| 2024BVD16910 | Herbivore | European fallow deer | *Dama dama* | Vorpommern-Rügen | 106 | negative | - | - | - | negative | - | - |

| | | | | | | | | | | | | |
|---|---|---|---|---|---|---|---|---|---|---|---|---|
| 2024BVD16911 | Herbivore | European fallow deer | *Dama dama* | Vorpommern-Rügen | 100 | negative | - | - | - | negative | - | - |
| 2024BVD16913 | Herbivore | European fallow deer | *Dama dama* | Vorpommern-Rügen | 115 | negative | - | - | - | negative | - | - |
| 2024BVD16914 | Herbivore | European fallow deer | *Dama dama* | Vorpommern-Rügen | 102 | negative | - | - | - | negative | - | - |
| 2024BVD16915 | Herbivore | European fallow deer | *Dama dama* | Vorpommern-Rügen | 112 | negative | - | - | - | negative | - | - |
| 2024BVD16916 | Herbivore | European fallow deer | *Dama dama* | Vorpommern-Rügen | 108 | negative | - | - | - | negative | - | - |
| 2024BVD16917 | Herbivore | European fallow deer | *Dama dama* | Vorpommern-Rügen | 110 | negative | - | - | - | negative | - | - |
| 2024BVD16918 | Herbivore | European fallow deer | *Dama dama* | Vorpommern-Rügen | 105 | negative | - | - | - | negative | - | - |
| 2024BVD16919 | Herbivore | European fallow deer | *Dama dama* | Vorpommern-Rügen | 103 | negative | - | - | - | negative | - | - |
| 2024BVD16920 | Herbivore | European fallow deer | *Dama dama* | Vorpommern-Rügen | 112 | negative | - | - | - | negative | - | - |
| 2024BVD16921 | Herbivore | European fallow deer | *Dama dama* | Vorpommern-Rügen | 114 | negative | - | - | - | negative | - | - |
| 2024BVD16922 | Omnivore | Wild boar | *Sus scrofa* | Vorpommern-Rügen | 100 | negative | - | - | - | negative | - | - |
| 2024BVD16923 | Herbivore | Roe deer | *Capreolus capreolus* | Vorpommern-Rügen | 97 | negative | - | - | - | negative | - | - |
| 2024BVD16924 | Herbivore | Roe deer | *Capreolus capreolus* | Vorpommern-Rügen | 98 | negative | - | - | - | negative | - | - |
| 2024BVD16925 | Herbivore | Mouflon | *Ovis gmelini musimon* | Vorpommern-Rügen | 112 | negative | - | - | - | negative | - | - |

| | | | | | | | | | | | | |
|---|---|---|---|---|---|---|---|---|---|---|---|---|
| 2024BVD16926 | Herbivore | Red deer | *Cervus elaphus* | Vorpommern-Rügen | 107 | negative | - | - | - | negative | - | - |
| 2024BVD16927 | Herbivore | Red deer | *Cervus elaphus* | Vorpommern-Rügen | 106 | negative | - | - | - | negative | - | - |
| 2024BVD16928 | Omnivore | Wild boar | *Sus scrofa* | Vorpommern-Rügen | 107 | negative | - | - | - | negative | - | - |
| 2024BVD16929 | Herbivore | Red deer | *Cervus elaphus* | Vorpommern-Rügen | 112 | negative | - | - | - | negative | - | - |
| 2024BVD16930 | Omnivore | Wild boar | *Sus scrofa* | Vorpommern-Rügen | 125 | negative | - | - | - | negative | - | - |
| 2024BVD16931 | Herbivore | Red deer | *Cervus elaphus* | Vorpommern-Rügen | 107 | negative | - | - | - | negative | - | - |
| 2024BVD16932 | Herbivore | Roe deer | *Capreolus capreolus* | Vorpommern-Rügen | 105 | negative | - | - | - | negative | - | - |
| 2024BVD16934 | Omnivore | Wild boar | *Sus scrofa* | Vorpommern-Rügen | 23 | positive | 33 | positive | <1:20 | negative | - | - |
| 2024BVD16935 | Omnivore | Wild boar | *Sus scrofa* | Vorpommern-Rügen | 102 | negative | - | - | - | negative | - | - |
| 2024BVD16936 | Omnivore | Wild boar | *Sus scrofa* | Vorpommern-Rügen | 111 | negative | - | - | - | negative | - | - |
| 2024BVD16937 | Herbivore | Red deer | *Cervus elaphus* | Vorpommern-Rügen | 109 | negative | - | - | - | negative | - | - |
| 2024BVD16938 | Herbivore | Red deer | *Cervus elaphus* | Vorpommern-Rügen | 102 | negative | - | - | - | negative | - | - |
| 2024BVD16939 | Omnivore | Wild boar | *Sus scrofa* | Vorpommern-Rügen | 69 | negative | - | - | - | negative | - | - |
| 2024BVD16940 | Omnivore | Wild boar | *Sus scrofa* | Vorpommern-Rügen | 109 | negative | - | - | - | negative | - | - |
| 2024BVD16941 | Omnivore | Wild boar | *Sus scrofa* | Vorpommern-Rügen | 116 | negative | - | - | - | negative | - | - |
| 2024BVD16942 | Omnivore | Wild boar | *Sus scrofa* | Vorpommern-Rügen | 99 | negative | - | - | - | negative | - | - |
| 2024BVD16943 | Herbivore | Red deer | *Cervus elaphus* | Vorpommern-Rügen | 110 | negative | - | - | - | negative | - | - |
| 2024BVD16944 | Herbivore | Red deer | *Cervus elaphus* | Vorpommern-Rügen | 105 | negative | - | - | - | negative | - | - |
| 2024BVD16945 | Herbivore | Red deer | *Cervus elaphus* | Vorpommern-Rügen | 106 | negative | - | - | - | negative | - | - |

| | | | | | | | | | | | | |
|---|---|---|---|---|---|---|---|---|---|---|---|---|
| 2024BVD16946 | Herbivore | Roe deer | *Capreolus capreolus* | Vorpommern-Rügen | 109 | negative | - | - | - | negative | - | - |
| 2024BVD16947 | Omnivore | Wild boar | *Sus scrofa* | Vorpommern-Rügen | 112 | negative | - | - | - | negative | - | - |
| 2024BVD16948 | Herbivore | Red deer | *Cervus elaphus* | Vorpommern-Rügen | 105 | negative | - | - | - | negative | - | - |
| 2024BVD16949 | Herbivore | Roe deer | *Capreolus capreolus* | Vorpommern-Rügen | 99 | negative | - | - | - | negative | - | - |
| 2024BVD16950 | Herbivore | Red deer | *Cervus elaphus* | Vorpommern-Rügen | 105 | negative | - | - | - | negative | - | - |
| 2024BVD16951 | Herbivore | Red deer | *Cervus elaphus* | Vorpommern-Rügen | 107 | negative | - | - | - | negative | - | - |
| 2024BVD16952 | Herbivore | Red deer | *Cervus elaphus* | Vorpommern-Rügen | 119 | negative | - | - | - | negative | - | - |
| 2024BVD16953 | Herbivore | Red deer | *Cervus elaphus* | Vorpommern-Rügen | 108 | negative | - | - | - | negative | - | - |
| 2024BVD16954 | Omnivore | Wild boar | *Sus scrofa* | Vorpommern-Rügen | 107 | negative | - | - | - | negative | - | - |
| 2024BVD16955 | Omnivore | Wild boar | *Sus scrofa* | Vorpommern-Rügen | 100 | negative | - | - | - | negative | - | - |
| 2024BVD16956 | Omnivore | Wild boar | *Sus scrofa* | Vorpommern-Rügen | 114 | negative | - | - | - | negative | - | - |
| 2024BVD16957 | Omnivore | Wild boar | *Sus scrofa* | Vorpommern-Rügen | 95 | negative | - | - | - | negative | - | - |
| 2024BVD16958 | Herbivore | Red deer | *Cervus elaphus* | Vorpommern-Rügen | 110 | negative | - | - | - | negative | - | - |
| 2024BVD16959 | Omnivore | Wild boar | *Sus scrofa* | Vorpommern-Rügen | 99 | negative | - | - | - | negative | - | - |
| 2024BVD16960 | Omnivore | Wild boar | *Sus scrofa* | Vorpommern-Rügen | 111 | negative | - | - | - | negative | - | - |
| 2024BVD16961 | Omnivore | Wild boar | *Sus scrofa* | Vorpommern-Rügen | 109 | negative | - | - | - | negative | - | - |
| 2024BVD16962 | Omnivore | Wild boar | *Sus scrofa* | Vorpommern-Rügen | 111 | negative | - | - | - | negative | - | - |
| 2024BVD16963 | Omnivore | Wild boar | *Sus scrofa* | Vorpommern-Rügen | 27 | positive | 17 | positive | <1:20 | negative | - | - |
| 2024BVD16964 | Herbivore | Roe deer | *Capreolus capreolus* | Mecklenburgische Seenplatte | 97 | negative | - | - | - | negative | - | - |

| | | | | | | | | | | | | |
|---|---|---|---|---|---|---|---|---|---|---|---|---|
| 2024BVD16965 | Omnivore | Wild boar | *Sus scrofa* | Mecklenburgische Seenplatte | 114 | negative | - | - | - | negative | - | - |
| 2024BVD16966 | Omnivore | Wild boar | *Sus scrofa* | Mecklenburgische Seenplatte | 107 | negative | - | - | - | negative | - | - |
| 2024BVD17016 | Herbivore | Roe deer | *Capreolus capreolus* | Vorpommern-Rügen | 107 | negative | - | - | - | negative | - | - |
| 2024BVD17017 | Herbivore | European fallow deer | *Dama dama* | Vorpommern-Rügen | 113 | negative | - | - | - | negative | - | - |
| 2024BVD17018 | Herbivore | Roe deer | *Capreolus capreolus* | Vorpommern-Rügen | 115 | negative | - | - | - | negative | - | - |
| 2024BVD17019 | Herbivore | European fallow deer | *Dama dama* | Vorpommern-Rügen | 109 | negative | - | - | - | negative | - | - |
| 2024BVD17021 | Herbivore | European fallow deer | *Dama dama* | Vorpommern-Rügen | 106 | negative | - | - | - | negative | - | - |
| 2024BVD17022 | Herbivore | European fallow deer | *Dama dama* | Vorpommern-Rügen | 97 | negative | - | - | - | negative | - | - |
| 2024BVD17023 | Herbivore | European fallow deer | *Dama dama* | Vorpommern-Rügen | 116 | negative | - | - | - | negative | - | - |
| 2024BVD17024 | Herbivore | European fallow deer | *Dama dama* | Vorpommern-Rügen | 116 | negative | - | - | - | negative | - | - |
| 2024BVD17025 | Herbivore | Roe deer | *Capreolus capreolus* | Vorpommern-Rügen | 117 | negative | - | - | - | negative | - | - |
| 2024BVD17026 | Herbivore | Roe deer | *Capreolus capreolus* | Vorpommern-Rügen | 68 | negative | - | - | - | negative | - | - |
| 2024BVD17027 | Herbivore | European fallow deer | *Dama dama* | Vorpommern-Rügen | 76 | negative | - | - | - | negative | - | - |
| 2024BVD17028 | Herbivore | European fallow deer | *Dama dama* | Vorpommern-Rügen | 108 | negative | - | - | - | negative | - | - |
| 2024BVD17029 | Omnivore | Wild boar | *Sus scrofa* | Vorpommern-Rügen | 96 | negative | - | - | - | negative | - | - |

| | | | | | | | | | | | | |
|---|---|---|---|---|---|---|---|---|---|---|---|---|
| 2024BVD17030 | Omnivore | Wild boar | *Sus scrofa* | Vorpommern-Rügen | 41 | positive | 100 | negative | <1:20 | negative | - | - |
| 2024BVD17031 | Omnivore | Wild boar | *Sus scrofa* | Vorpommern-Rügen | 94 | negative | - | - | - | negative | - | - |
| 2024BVD17032 | Omnivore | Wild boar | *Sus scrofa* | Vorpommern-Rügen | 134 | negative | - | - | - | negative | - | - |
| 2024BVD17033 | Omnivore | Wild boar | *Sus scrofa* | Vorpommern-Rügen | 94 | negative | - | - | - | negative | - | - |
| 2024BVD17034 | Herbivore | European fallow deer | *Dama dama* | Vorpommern-Rügen | 96 | negative | - | - | - | negative | - | - |
| 2024BVD17035 | Herbivore | Red deer | *Cervus elaphus* | Vorpommern-Rügen | 90 | negative | - | - | - | negative | - | - |
| 2024BVD17036 | Herbivore | European fallow deer | *Dama dama* | Vorpommern-Rügen | 78 | negative | - | - | - | negative | - | - |
| 2024BVD17037 | Herbivore | European fallow deer | *Dama dama* | Vorpommern-Rügen | 97 | negative | - | - | - | negative | - | - |
| 2024BVD17038 | Herbivore | European fallow deer | *Dama dama* | Vorpommern-Rügen | 94 | negative | - | - | - | negative | - | - |
| 2024BVD17039 | Herbivore | European fallow deer | *Dama dama* | Vorpommern-Rügen | 97 | negative | - | - | - | negative | - | - |
| 2024BVD17040 | Herbivore | Roe deer | *Capreolus capreolus* | Vorpommern-Rügen | 94 | negative | - | - | - | negative | - | - |
| 2024BVD17041 | Herbivore | European fallow deer | *Dama dama* | Vorpommern-Rügen | 89 | negative | - | - | - | negative | - | - |
| 2024BVD17042 | Herbivore | Roe deer | *Capreolus capreolus* | Vorpommern-Rügen | 92 | negative | - | - | - | negative | - | - |
| 2024BVD17043 | Herbivore | European fallow deer | *Dama dama* | Vorpommern-Rügen | 87 | negative | - | - | - | negative | - | - |
| 2024BVD17044 | Herbivore | European fallow deer | *Dama dama* | Vorpommern-Rügen | 91 | negative | - | - | - | negative | - | - |

| | | | | | | | | | | | | |
|---|---|---|---|---|---|---|---|---|---|---|---|---|
| 2024BVD17045 | Omnivore | Wild boar | *Sus scrofa* | Vorpommern-Rügen | 97 | negative | - | - | - | negative | - | - |
| 2024BVD17046 | Omnivore | Wild boar | *Sus scrofa* | Vorpommern-Rügen | 94 | negative | - | - | - | negative | - | - |
| 2024BVD17047 | Omnivore | Wild boar | *Sus scrofa* | Vorpommern-Rügen | 99 | negative | - | - | - | negative | - | - |
| 2024BVD17048 | Omnivore | Wild boar | *Sus scrofa* | Vorpommern-Rügen | 98 | negative | - | - | - | negative | - | - |
| 2024BVD17049 | Omnivore | Wild boar | *Sus scrofa* | Vorpommern-Rügen | 93 | negative | - | - | - | negative | - | - |
| 2024BVD17050 | Omnivore | Wild boar | *Sus scrofa* | Vorpommern-Rügen | 92 | negative | - | - | - | negative | - | - |
| 2024BVD17051 | Omnivore | Wild boar | *Sus scrofa* | Vorpommern-Rügen | 94 | negative | - | - | - | negative | - | - |
| 2024BVD17052 | Herbivore | European fallow deer | *Dama dama* | Vorpommern-Rügen | 96 | negative | - | - | - | negative | - | - |
| 2024BVD17053 | Omnivore | Wild boar | *Sus scrofa* | Vorpommern-Rügen | 93 | negative | - | - | - | negative | - | - |
| 2024BVD17055 | Omnivore | Wild boar | *Sus scrofa* | Vorpommern-Rügen | 103 | negative | - | - | - | negative | - | - |
| 2024BVD17056 | Herbivore | European fallow deer | *Dama dama* | Vorpommern-Rügen | 79 | negative | - | - | - | negative | - | - |
| 2024BVD17058 | Herbivore | European fallow deer | *Dama dama* | Vorpommern-Rügen | 94 | negative | - | - | - | negative | - | - |
| 2024BVD17059 | Omnivore | Wild boar | *Sus scrofa* | Vorpommern-Rügen | 92 | negative | - | - | - | negative | - | - |
| 2024BVD17060 | Omnivore | Wild boar | *Sus scrofa* | Vorpommern-Rügen | 94 | negative | - | - | - | negative | - | - |
| 2024BVD17061 | Omnivore | Wild boar | *Sus scrofa* | Vorpommern-Rügen | 96 | negative | - | - | - | negative | - | - |
| 2024BVD17062 | Omnivore | Wild boar | *Sus scrofa* | Vorpommern-Rügen | 70 | negative | - | - | - | negative | - | - |
| 2024BVD17063 | Omnivore | Wild boar | *Sus scrofa* | Vorpommern-Rügen | 103 | negative | - | - | - | negative | - | - |

| | | | | | | | | | | | | |
|---|---|---|---|---|---|---|---|---|---|---|---|---|
| 2024BVD17064 | Omnivore | Wild boar | *Sus scrofa* | Vorpommern-Rügen | 103 | negative | - | - | - | negative | - | - |
| 2024BVD17065 | Omnivore | Wild boar | *Sus scrofa* | Vorpommern-Rügen | 121 | negative | - | - | - | negative | - | - |
| 2024BVD17066 | Omnivore | Wild boar | *Sus scrofa* | Vorpommern-Rügen | 95 | negative | - | - | - | negative | - | - |
| 2024BVD17067 | Omnivore | Wild boar | *Sus scrofa* | Vorpommern-Rügen | 87 | negative | - | - | - | negative | - | - |
| 2024BVD17160 | Omnivore | Wild boar | *Sus scrofa* | Vorpommern-Greifswald | 93 | negative | - | - | - | negative | - | - |
| 2024BVD17161 | Herbivore | Roe deer | *Capreolus capreolus* | Vorpommern-Greifswald | 97 | negative | - | - | - | negative | - | - |
| 2024BVD17162 | Omnivore | Wild boar | *Sus scrofa* | Vorpommern-Greifswald | 88 | negative | - | - | - | negative | - | - |
| 2024BVD17164 | Omnivore | Wild boar | *Sus scrofa* | Vorpommern-Greifswald | 95 | negative | - | - | - | negative | - | - |
| 2024BVD17165 | Herbivore | Roe deer | *Capreolus capreolus* | Vorpommern-Greifswald | 64 | negative | - | - | - | negative | - | - |
| 2024BVD17166 | Herbivore | Roe deer | *Capreolus capreolus* | Vorpommern-Greifswald | 98 | negative | - | - | - | negative | - | - |
| 2024BVD17167 | Omnivore | Wild boar | *Sus scrofa* | Vorpommern-Greifswald | 93 | negative | - | - | - | negative | - | - |
| 2024BVD17168 | Herbivore | Roe deer | *Capreolus capreolus* | Vorpommern-Greifswald | 86 | negative | - | - | - | negative | - | - |
| 2024BVD17169 | Herbivore | Roe deer | *Capreolus capreolus* | Vorpommern-Greifswald | 100 | negative | - | - | - | negative | - | - |
| 2024BVD17170 | Omnivore | Wild boar | *Sus scrofa* | Vorpommern-Greifswald | 97 | negative | - | - | - | negative | - | - |
| 2024BVD17171 | Omnivore | Wild boar | *Sus scrofa* | Vorpommern-Greifswald | 86 | negative | - | - | - | negative | - | - |
| 2024BVD17172 | Omnivore | Wild boar | *Sus scrofa* | Vorpommern-Greifswald | 97 | negative | - | - | - | negative | - | - |
| 2024BVD17173 | Omnivore | Wild boar | *Sus scrofa* | Vorpommern-Greifswald | 95 | negative | - | - | - | negative | - | - |
| 2024BVD17174 | Herbivore | Roe deer | *Capreolus capreolus* | Vorpommern-Greifswald | 92 | negative | - | - | - | negative | - | - |
| 2024BVD17175 | Omnivore | Wild boar | *Sus scrofa* | Vorpommern-Greifswald | 93 | negative | - | - | - | negative | - | - |

| | | | | | | | | | | | | |
|---|---|---|---|---|---|---|---|---|---|---|---|---|
| 2024BVD17176 | Omnivore | Wild boar | *Sus scrofa* | Vorpommern-Greifswald | 69 | negative | - | - | - | negative | - | - |
| 2024BVD17177 | Omnivore | Wild boar | *Sus scrofa* | Vorpommern-Greifswald | 95 | negative | - | - | - | negative | - | - |
| 2024BVD17178 | Herbivore | Roe deer | *Capreolus capreolus* | Vorpommern-Greifswald | 95 | negative | - | - | - | negative | - | - |
| 2024BVD17179 | Herbivore | Roe deer | *Capreolus capreolus* | Vorpommern-Greifswald | 89 | negative | - | - | - | negative | - | - |
| 2024BVD17180 | Omnivore | Wild boar | *Sus scrofa* | Vorpommern-Greifswald | 92 | negative | - | - | - | negative | - | - |
| 2024BVD17181 | Omnivore | Wild boar | *Sus scrofa* | Vorpommern-Greifswald | 90 | negative | - | - | - | negative | - | - |
| 2024BVD17183 | Omnivore | Wild boar | *Sus scrofa* | Vorpommern-Greifswald | 95 | negative | - | - | - | negative | - | - |
| 2024BVD17184 | Herbivore | Roe deer | *Capreolus capreolus* | Vorpommern-Greifswald | 70 | negative | - | - | - | negative | - | - |
| 2024BVD17185 | Herbivore | Roe deer | *Capreolus capreolus* | Vorpommern-Greifswald | 98 | negative | - | - | - | negative | - | - |
| 2024BVD17186 | Herbivore | Roe deer | *Capreolus capreolus* | Vorpommern-Greifswald | 92 | negative | - | - | - | negative | - | - |
| 2024BVD17187 | Omnivore | Wild boar | *Sus scrofa* | Vorpommern-Greifswald | 96 | negative | - | - | - | negative | - | - |
| 2024BVD17188 | Omnivore | Wild boar | *Sus scrofa* | Vorpommern-Greifswald | 87 | negative | - | - | - | negative | - | - |
| 2024BVD17189 | Omnivore | Wild boar | *Sus scrofa* | Vorpommern-Greifswald | 86 | negative | - | - | - | negative | - | - |
| 2024BVD17190 | Omnivore | Wild boar | *Sus scrofa* | Vorpommern-Greifswald | 87 | negative | - | - | - | negative | - | - |
| 2024BVD17191 | Herbivore | European fallow deer | *Dama dama* | Mecklenburgische Seenplatte | 95 | negative | - | - | - | negative | - | - |
| 2024BVD17192 | Omnivore | Wild boar | *Sus scrofa* | Mecklenburgische Seenplatte | 95 | negative | - | - | - | negative | - | - |
| 2024BVD17193 | Omnivore | Wild boar | *Sus scrofa* | Mecklenburgische Seenplatte | 96 | negative | - | - | - | negative | - | - |
| 2024BVD17194 | Herbivore | Roe deer | *Capreolus capreolus* | Mecklenburgische Seenplatte | 89 | negative | - | - | - | negative | - | - |

| | | | | | | | | | | | | |
|---|---|---|---|---|---|---|---|---|---|---|---|---|
| 2024BVD17195 | Herbivore | Roe deer | *Capreolus capreolus* | Mecklenburgische Seenplatte | 102 | negative | - | - | - | negative | - | - |
| 2024BVD17196 | Herbivore | European fallow deer | *Dama dama* | Mecklenburgische Seenplatte | 92 | negative | - | - | - | negative | - | - |
| 2024BVD17197 | Herbivore | Roe deer | *Capreolus capreolus* | Mecklenburgische Seenplatte | 97 | negative | - | - | - | negative | - | - |
| 2024BVD17198 | Herbivore | Roe deer | *Capreolus capreolus* | Mecklenburgische Seenplatte | 95 | negative | - | - | - | negative | - | - |
| 2024BVD17199 | Herbivore | Roe deer | *Capreolus capreolus* | Mecklenburgische Seenplatte | 100 | negative | - | - | - | negative | - | - |
| 2024BVD17200 | Omnivore | Wild boar | *Sus scrofa* | Mecklenburgische Seenplatte | 93 | negative | - | - | - | negative | - | - |
| 2024BVD17201 | Herbivore | Roe deer | *Capreolus capreolus* | Mecklenburgische Seenplatte | 78 | negative | - | - | - | negative | - | - |
| 2024BVD17202 | Herbivore | Roe deer | *Capreolus capreolus* | Mecklenburgische Seenplatte | 101 | negative | - | - | - | negative | - | - |
| 2024BVD17203 | Herbivore | Roe deer | *Capreolus capreolus* | Mecklenburgische Seenplatte | 97 | negative | - | - | - | negative | - | - |
| 2024BVD17204 | Herbivore | Roe deer | *Capreolus capreolus* | Mecklenburgische Seenplatte | 106 | negative | - | - | - | negative | - | - |
| 2024BVD17205 | Herbivore | European fallow deer | *Dama dama* | Mecklenburgische Seenplatte | 104 | negative | - | - | - | negative | - | - |
| 2024BVD17206 | Omnivore | Wild boar | *Sus scrofa* | Mecklenburgische Seenplatte | 104 | negative | - | - | - | negative | - | - |
| 2024BVD17208 | Omnivore | Wild boar | *Sus scrofa* | Mecklenburgische Seenplatte | 91 | negative | - | - | - | negative | - | - |
| 2024BVD17209 | Herbivore | Roe deer | *Capreolus capreolus* | Mecklenburgische Seenplatte | 96 | negative | - | - | - | negative | - | - |
| 2024BVD17210 | Herbivore | European fallow deer | *Dama dama* | Mecklenburgische Seenplatte | 93 | negative | - | - | - | negative | - | - |
| 2024BVD17211 | Omnivore | Wild boar | *Sus scrofa* | Mecklenburgische Seenplatte | 106 | negative | - | - | - | negative | - | - |
| 2024BVD17212 | Omnivore | Wild boar | *Sus scrofa* | Mecklenburgische Seenplatte | 99 | negative | - | - | - | negative | - | - |

| | | | | | | | | | | | | |
|---|---|---|---|---|---|---|---|---|---|---|---|---|
| 2024BVD17213 | Omnivore | Wild boar | *Sus scrofa* | Mecklenburgische Seenplatte | 89 | negative | - | - | - | negative | - | - |
| 2024BVD17214 | Omnivore | Wild boar | *Sus scrofa* | Mecklenburgische Seenplatte | 96 | negative | - | - | - | negative | - | - |
| 2024BVD17215 | Omnivore | Wild boar | *Sus scrofa* | Mecklenburgische Seenplatte | 87 | negative | - | - | - | negative | - | - |
| 2024BVD17216 | Omnivore | Wild boar | *Sus scrofa* | Mecklenburgische Seenplatte | 100 | negative | - | - | - | negative | - | - |
| 2024BVD17217 | Herbivore | European fallow deer | *Dama dama* | Mecklenburgische Seenplatte | 78 | negative | - | - | - | negative | - | - |
| 2024BVD17218 | Omnivore | Wild boar | *Sus scrofa* | Mecklenburgische Seenplatte | 99 | negative | - | - | - | negative | - | - |
| 2024BVD17219 | Herbivore | Roe deer | *Capreolus capreolus* | Mecklenburgische Seenplatte | 96 | negative | - | - | - | negative | - | - |
| 2024BVD17220 | Omnivore | Wild boar | *Sus scrofa* | Mecklenburgische Seenplatte | 98 | negative | - | - | - | negative | - | - |
| 2024BVD17221 | Omnivore | Wild boar | *Sus scrofa* | Mecklenburgische Seenplatte | 87 | negative | - | - | - | negative | - | - |
| 2024BVD17222 | Omnivore | Wild boar | *Sus scrofa* | Mecklenburgische Seenplatte | 105 | negative | - | - | - | negative | - | - |
| 2024BVD17223 | Herbivore | unknown | unknown | Mecklenburgische Seenplatte | 87 | negative | - | - | - | negative | - | - |
| 2024BVD17224 | Herbivore | Roe deer | *Capreolus capreolus* | Mecklenburgische Seenplatte | 77 | negative | - | - | - | negative | - | - |
| 2024BVD17225 | Herbivore | Roe deer | *Capreolus capreolus* | Mecklenburgische Seenplatte | 106 | negative | - | - | - | negative | - | - |
| 2024BVD17226 | Herbivore | Roe deer | *Capreolus capreolus* | Mecklenburgische Seenplatte | 103 | negative | - | - | - | negative | - | - |
| 2024BVD17227 | Herbivore | Roe deer | *Capreolus capreolus* | Mecklenburgische Seenplatte | 100 | negative | - | - | - | negative | - | - |
| 2024BVD17228 | Herbivore | Roe deer | *Capreolus capreolus* | Mecklenburgische Seenplatte | 95 | negative | - | - | - | negative | - | - |
| 2024BVD17229 | Herbivore | Roe deer | *Capreolus capreolus* | Mecklenburgische Seenplatte | 79 | negative | - | - | - | negative | - | - |
| 2024BVD17230 | Herbivore | Roe deer | *Capreolus capreolus* | Mecklenburgische Seenplatte | 73 | negative | - | - | - | negative | - | - |

| | | | | | | | | | | | | |
|---|---|---|---|---|---|---|---|---|---|---|---|---|
| 2024BVD17231 | Herbivore | Roe deer | *Capreolus capreolus* | Mecklenburgische Seenplatte | 86 | negative | - | - | - | negative | - | - |
| 2024BVD17232 | Herbivore | European fallow deer | *Dama dama* | Mecklenburgische Seenplatte | 111 | negative | - | - | - | negative | - | - |
| 2024BVD17233 | Herbivore | Roe deer | *Capreolus capreolus* | Mecklenburgische Seenplatte | 86 | negative | - | - | - | negative | - | - |
| 2024BVD17234 | Herbivore | Roe deer | *Capreolus capreolus* | Mecklenburgische Seenplatte | 75 | negative | - | - | - | negative | - | - |
| 2024BVD17235 | Herbivore | Roe deer | *Capreolus capreolus* | Mecklenburgische Seenplatte | 96 | negative | - | - | - | negative | - | - |
| 2024BVD17236 | Herbivore | Roe deer | *Capreolus capreolus* | Mecklenburgische Seenplatte | 91 | negative | - | - | - | negative | - | - |
| 2024BVD17237 | Herbivore | Roe deer | *Capreolus capreolus* | Mecklenburgische Seenplatte | 77 | negative | - | - | - | negative | - | - |
| 2024BVD17238 | Herbivore | European fallow deer | *Dama dama* | Mecklenburgische Seenplatte | 92 | negative | - | - | - | negative | - | - |
| 2024BVD17239 | Herbivore | European fallow deer | *Dama dama* | Mecklenburgische Seenplatte | 91 | negative | - | - | - | negative | - | - |
| 2024BVD17240 | Herbivore | Roe deer | *Capreolus capreolus* | Mecklenburgische Seenplatte | 95 | negative | - | - | - | negative | - | - |
| 2024BVD17241 | Herbivore | Roe deer | *Capreolus capreolus* | Mecklenburgische Seenplatte | 104 | negative | - | - | - | negative | - | - |
| 2024BVD17242 | Herbivore | Roe deer | *Capreolus capreolus* | Mecklenburgische Seenplatte | 103 | negative | - | - | - | negative | - | - |
| 2024BVD17243 | Herbivore | Roe deer | *Capreolus capreolus* | Mecklenburgische Seenplatte | 99 | negative | - | - | - | negative | - | - |
| 2024BVD17244 | Herbivore | Roe deer | *Capreolus capreolus* | Mecklenburgische Seenplatte | 105 | negative | - | - | - | negative | - | - |
| 2024BVD17245 | Herbivore | Roe deer | *Capreolus capreolus* | Mecklenburgische Seenplatte | 103 | negative | - | - | - | negative | - | - |
| 2024BVD17246 | Herbivore | Roe deer | *Capreolus capreolus* | Mecklenburgische Seenplatte | 97 | negative | - | - | - | negative | - | - |
| 2024BVD17247 | Herbivore | Roe deer | *Capreolus capreolus* | Mecklenburgische Seenplatte | 103 | negative | - | - | - | negative | - | - |

| | | | | | | | | | | | | |
|---|---|---|---|---|---|---|---|---|---|---|---|---|
| 2024BVD17248 | Herbivore | Roe deer | *Capreolus capreolus* | Mecklenburgische Seenplatte | 93 | negative | - | - | - | negative | - | - |
| 2024BVD17249 | Herbivore | Roe deer | *Capreolus capreolus* | Mecklenburgische Seenplatte | 86 | negative | - | - | - | negative | - | - |
| 2024BVD17257 | Herbivore | European fallow deer | *Dama dama* | Mecklenburgische Seenplatte | 86 | negative | - | - | - | negative | - | - |
| 2024BVD17258 | Herbivore | Roe deer | *Capreolus capreolus* | Mecklenburgische Seenplatte | 75 | negative | - | - | - | negative | - | - |
| 2024BVD17259 | Herbivore | Roe deer | *Capreolus capreolus* | Mecklenburgische Seenplatte | 96 | negative | - | - | - | negative | - | - |
| 2024BVD17260 | Herbivore | Roe deer | *Capreolus capreolus* | Mecklenburgische Seenplatte | 91 | negative | - | - | - | negative | - | - |
| 2024BVD17261 | Herbivore | Roe deer | *Capreolus capreolus* | Mecklenburgische Seenplatte | 77 | negative | - | - | - | negative | - | - |
| 2024BVD17262 | Herbivore | Roe deer | *Capreolus capreolus* | Mecklenburgische Seenplatte | 92 | negative | - | - | - | negative | - | - |
| 2024BVD17263 | Herbivore | Roe deer | *Capreolus capreolus* | Mecklenburgische Seenplatte | 91 | negative | - | - | - | negative | - | - |
| 2024BVD17264 | Herbivore | Roe deer | *Capreolus capreolus* | Mecklenburgische Seenplatte | 95 | negative | - | - | - | negative | - | - |
| 2024BVD17265 | Herbivore | Roe deer | *Capreolus capreolus* | Mecklenburgische Seenplatte | 104 | negative | - | - | - | negative | - | - |
| 2024BVD17266 | Herbivore | Roe deer | *Capreolus capreolus* | Mecklenburgische Seenplatte | 103 | negative | - | - | - | negative | - | - |
| 2024BVD17267 | Herbivore | Roe deer | *Capreolus capreolus* | Mecklenburgische Seenplatte | 99 | negative | - | - | - | negative | - | - |
| 2024BVD17268 | Herbivore | Roe deer | *Capreolus capreolus* | Mecklenburgische Seenplatte | 105 | negative | - | - | - | negative | - | - |
| 2024BVD17269 | Herbivore | Roe deer | *Capreolus capreolus* | Mecklenburgische Seenplatte | 103 | negative | - | - | - | negative | - | - |
| 2024BVD17270 | Herbivore | Roe deer | *Capreolus capreolus* | Mecklenburgische Seenplatte | 97 | negative | - | - | - | negative | - | - |
| 2024BVD17271 | Herbivore | Roe deer | *Capreolus capreolus* | Mecklenburgische Seenplatte | 103 | negative | - | - | - | negative | - | - |
| 2024BVD17272 | Herbivore | Roe deer | *Capreolus capreolus* | Mecklenburgische Seenplatte | 93 | negative | - | - | - | negative | - | - |

| | | | | | | | | | | | | |
|---|---|---|---|---|---|---|---|---|---|---|---|---|
| 2024BVD17273 | Herbivore | Roe deer | *Capreolus capreolus* | Mecklenburgische Seenplatte | 86 | negative | - | - | - | negative | - | - |
| 2024BVD17274 | Herbivore | Roe deer | *Capreolus capreolus* | Mecklenburgische Seenplatte | 106 | negative | - | - | - | negative | - | - |
| 2024BVD17275 | Herbivore | Roe deer | *Capreolus capreolus* | Mecklenburgische Seenplatte | 107 | negative | - | - | - | negative | - | - |
| 2024BVD17276 | Herbivore | European fallow deer | *Dama dama* | Mecklenburgische Seenplatte | 97 | negative | - | - | - | negative | - | - |
| 2024BVD17277 | Herbivore | Roe deer | *Capreolus capreolus* | Mecklenburgische Seenplatte | 105 | negative | - | - | - | negative | - | - |
| 2024BVD17278 | Omnivore | Wild boar | *Sus scrofa* | Mecklenburgische Seenplatte | 110 | negative | - | - | - | negative | - | - |
| 2024BVD17279 | Herbivore | Roe deer | *Capreolus capreolus* | Mecklenburgische Seenplatte | 93 | negative | - | - | - | negative | - | - |
| 2024BVD17280 | Omnivore | Wild boar | *Sus scrofa* | Mecklenburgische Seenplatte | 94 | negative | - | - | - | negative | - | - |
| 2024BVD17281 | Omnivore | Wild boar | *Sus scrofa* | Mecklenburgische Seenplatte | 97 | negative | - | - | - | negative | - | - |
| 2024BVD17282 | Herbivore | Roe deer | *Capreolus capreolus* | Mecklenburgische Seenplatte | 101 | negative | - | - | - | negative | - | - |
| 2024BVD17283 | Omnivore | Wild boar | *Sus scrofa* | Mecklenburgische Seenplatte | 106 | negative | - | - | - | negative | - | - |
| 2024BVD17284 | Omnivore | Wild boar | *Sus scrofa* | Mecklenburgische Seenplatte | 84 | negative | - | - | - | negative | - | - |
| 2024BVD17285 | Omnivore | Wild boar | *Sus scrofa* | Mecklenburgische Seenplatte | 110 | negative | - | - | - | negative | - | - |
| 2024BVD17286 | Omnivore | Wild boar | *Sus scrofa* | Mecklenburgische Seenplatte | 104 | negative | - | - | - | negative | - | - |
| 2024BVD17287 | Omnivore | Wild boar | *Sus scrofa* | Mecklenburgische Seenplatte | 106 | negative | - | - | - | negative | - | - |
| 2024BVD17288 | Omnivore | Wild boar | *Sus scrofa* | Mecklenburgische Seenplatte | 115 | negative | - | - | - | negative | - | - |
| 2024BVD17289 | Omnivore | Wild boar | *Sus scrofa* | Mecklenburgische Seenplatte | 107 | negative | - | - | - | negative | - | - |
| 2024BVD17369 | Herbivore | European fallow deer | *Dama dama* | Vorpommern-Rügen | 102 | negative | - | - | - | negative | - | - |

| | | | | | | | | | | | | |
|---|---|---|---|---|---|---|---|---|---|---|---|---|
| 2024BVD17370 | Herbivore | European fallow deer | *Dama dama* | Vorpommern-Rügen | 114 | negative | - | - | - | negative | - | - |
| 2024BVD17371 | Herbivore | European fallow deer | *Dama dama* | Vorpommern-Rügen | 103 | negative | - | - | - | negative | - | - |
| 2024BVD17372 | Herbivore | European fallow deer | *Dama dama* | Vorpommern-Rügen | 81 | negative | - | - | - | negative | - | - |
| 2024BVD17373 | Herbivore | European fallow deer | *Dama dama* | Vorpommern-Rügen | 104 | negative | - | - | - | negative | - | - |
| 2024BVD17374 | Herbivore | European fallow deer | *Dama dama* | Vorpommern-Rügen | 102 | negative | - | - | - | negative | - | - |
| 2024BVD17375 | Herbivore | Roe deer | *Capreolus capreolus* | Vorpommern-Rügen | 109 | negative | - | - | - | negative | - | - |
| 2024BVD17376 | Herbivore | Roe deer | *Capreolus capreolus* | Vorpommern-Rügen | 107 | negative | - | - | - | negative | - | - |
| 2024BVD17377 | Herbivore | European fallow deer | *Dama dama* | Vorpommern-Rügen | 97 | negative | - | - | - | negative | - | - |
| 2024BVD17378 | Herbivore | Roe deer | *Capreolus capreolus* | Vorpommern-Rügen | 105 | negative | - | - | - | negative | - | - |
| 2024BVD17379 | Herbivore | European fallow deer | *Dama dama* | Vorpommern-Rügen | 102 | negative | - | - | - | negative | - | - |
| 2024BVD17380 | Omnivore | Wild boar | *Sus scrofa* | Vorpommern-Rügen | 104 | negative | - | - | - | negative | - | - |
| 2024BVD17381 | Omnivore | Wild boar | *Sus scrofa* | Vorpommern-Rügen | 107 | negative | - | - | - | negative | - | - |
| 2024BVD17382 | Herbivore | European fallow deer | *Dama dama* | Vorpommern-Rügen | 106 | negative | - | - | - | negative | - | - |
| 2024BVD17383 | Herbivore | European fallow deer | *Dama dama* | Vorpommern-Rügen | 110 | negative | - | - | - | negative | - | - |

| | | | | | | | | | | | | |
|---|---|---|---|---|---|---|---|---|---|---|---|---|
| 2024BVD17467 | Herbivore | European fallow deer | *Dama dama* | Vorpommern-Rügen | 108 | negative | - | - | - | negative | - | - |
| 2024BVD17468 | Herbivore | European fallow deer | *Dama dama* | Vorpommern-Rügen | 97 | negative | - | - | - | negative | - | - |
| 2024BVD17469 | Omnivore | Wild boar | *Sus scrofa* | Vorpommern-Rügen | 100 | negative | - | - | - | negative | - | - |
| 2024BVD17470 | Omnivore | Wild boar | *Sus scrofa* | Vorpommern-Rügen | 103 | negative | - | - | - | negative | - | - |
| 2024BVD17471 | Herbivore | European fallow deer | *Dama dama* | Vorpommern-Rügen | 89 | negative | - | - | - | negative | - | - |
| 2024BVD17472 | Herbivore | European fallow deer | *Dama dama* | Vorpommern-Rügen | 88 | negative | - | - | - | negative | - | - |
| 2024BVD17473 | Herbivore | European fallow deer | *Dama dama* | Vorpommern-Rügen | 94 | negative | - | - | - | negative | - | - |
| 2024BVD17474 | Herbivore | European fallow deer | *Dama dama* | Vorpommern-Rügen | 94 | negative | - | - | - | negative | - | - |
| 2024BVD17475 | Herbivore | European fallow deer | *Dama dama* | Vorpommern-Rügen | 92 | negative | - | - | - | negative | - | - |
| 2024BVD17476 | Herbivore | European fallow deer | *Dama dama* | Vorpommern-Rügen | 89 | negative | - | - | - | negative | - | - |
| 2024BVD17477 | Herbivore | European fallow deer | *Dama dama* | Vorpommern-Rügen | 106 | negative | - | - | - | negative | - | - |
| 2024BVD17478 | Omnivore | Wild boar | *Sus scrofa* | Vorpommern-Rügen | 99 | negative | - | - | - | negative | - | - |
| 2024BVD17479 | Herbivore | European fallow deer | *Dama dama* | Vorpommern-Rügen | 95 | negative | - | - | - | negative | - | - |
| 2024BVD17480 | Omnivore | Wild boar | *Sus scrofa* | Vorpommern-Rügen | 85 | negative | - | - | - | negative | - | - |

| | | | | | | | | | | | | |
|---|---|---|---|---|---|---|---|---|---|---|---|---|
| 2024BVD17481 | Omnivore | Wild boar | *Sus scrofa* | Vorpommern-Rügen | 42 | positive | 4 | positive | 1:20 | negative | - | - |
| 2024BVD17482 | Omnivore | Wild boar | *Sus scrofa* | Vorpommern-Rügen | 20 | positive | 5 | positive | <1:20 | negative | - | - |
| 2024BVD17483 | Herbivore | Roe deer | *Capreolus capreolus* | Vorpommern-Rügen | 98 | negative | - | - | - | negative | - | - |
| 2024BVD17484 | Herbivore | European fallow deer | *Dama dama* | Vorpommern-Rügen | 89 | negative | - | - | - | negative | - | - |
| 2024BVD17485 | Herbivore | European fallow deer | *Dama dama* | Vorpommern-Rügen | 114 | negative | - | - | - | negative | - | - |
| 2024BVD17487 | Omnivore | Wild boar | *Sus scrofa* | Vorpommern-Rügen | 31 | positive | 53 | undetermined | <1:20 | negative | - | - |
| 2024BVD17488 | Herbivore | Roe deer | *Capreolus capreolus* | Vorpommern-Rügen | 103 | negative | - | - | - | negative | - | - |
| 2024BVD17489 | Herbivore | European fallow deer | *Dama dama* | Vorpommern-Rügen | 101 | negative | - | - | - | negative | - | - |
| 2024BVD17490 | Herbivore | European fallow deer | *Dama dama* | Vorpommern-Rügen | 102 | negative | - | - | - | negative | - | - |
| 2024BVD17491 | Herbivore | European fallow deer | *Dama dama* | Vorpommern-Rügen | 99 | negative | - | - | - | negative | - | - |
| 2024BVD17492 | Herbivore | European fallow deer | *Dama dama* | Vorpommern-Rügen | 90 | negative | - | - | - | negative | - | - |
| 2024BVD17493 | Herbivore | European fallow deer | *Dama dama* | Vorpommern-Rügen | 103 | negative | - | - | - | negative | - | - |
| 2024BVD17494 | Herbivore | European fallow deer | *Dama dama* | Vorpommern-Rügen | 98 | negative | - | - | - | negative | - | - |
| 2024BVD17495 | Herbivore | European fallow deer | *Dama dama* | Vorpommern-Rügen | 102 | negative | - | - | - | negative | - | - |

| | | | | | | | | | | | | |
|---|---|---|---|---|---|---|---|---|---|---|---|---|
| 2024BVD17496 | Omnivore | Wild boar | *Sus scrofa* | Vorpommern-Rügen | 100 | negative | - | - | - | negative | - | - |
| 2024BVD17497 | Herbivore | European fallow deer | *Dama dama* | Vorpommern-Rügen | 97 | negative | - | - | - | negative | - | - |
| 2024BVD17498 | Herbivore | European fallow deer | *Dama dama* | Vorpommern-Rügen | 104 | negative | - | - | - | negative | - | - |
| 2024BVD17499 | Herbivore | European fallow deer | *Dama dama* | Vorpommern-Rügen | 101 | negative | - | - | - | negative | - | - |
| 2024BVD17500 | Herbivore | European fallow deer | *Dama dama* | Vorpommern-Rügen | 98 | negative | - | - | - | negative | - | - |
| 2024BVD17501 | Herbivore | European fallow deer | *Dama dama* | Vorpommern-Rügen | 97 | negative | - | - | - | negative | - | - |
| 2024BVD17502 | Omnivore | Wild boar | *Sus scrofa* | Vorpommern-Rügen | 96 | negative | - | - | - | negative | - | - |
| 2024BVD17503 | Omnivore | Wild boar | *Sus scrofa* | Vorpommern-Rügen | 100 | negative | - | - | - | negative | - | - |
| 2024BVD17504 | Herbivore | Red deer | *Cervus elaphus* | Vorpommern-Rügen | 88 | negative | - | - | - | negative | - | - |
| 2024BVD17505 | Herbivore | Red deer | *Cervus elaphus* | Vorpommern-Rügen | 92 | negative | - | - | - | negative | - | - |
| 2024BVD17506 | Herbivore | European fallow deer | *Dama dama* | Vorpommern-Rügen | 100 | negative | - | - | - | negative | - | - |
| 2024BVD17507 | Herbivore | European fallow deer | *Dama dama* | Vorpommern-Rügen | 93 | negative | - | - | - | negative | - | - |
| 2024BVD17508 | Herbivore | European fallow deer | *Dama dama* | Vorpommern-Rügen | 92 | negative | - | - | - | negative | - | - |
| 2024BVD17509 | Herbivore | European fallow deer | *Dama dama* | Vorpommern-Rügen | 106 | negative | - | - | - | negative | - | - |

| | | | | | | | | | | | | |
|---|---|---|---|---|---|---|---|---|---|---|---|---|
| 2024BVD17510 | Herbivore | European fallow deer | *Dama dama* | Vorpommern-Rügen | 104 | negative | - | - | - | negative | - | - |
| 2024BVD17511 | Herbivore | European fallow deer | *Dama dama* | Vorpommern-Rügen | 103 | negative | - | - | - | negative | - | - |
| 2024BVD17512 | Herbivore | European fallow deer | *Dama dama* | Vorpommern-Rügen | 101 | negative | - | - | - | negative | - | - |
| 2024BVD17513 | Herbivore | European fallow deer | *Dama dama* | Vorpommern-Rügen | 98 | negative | - | - | - | negative | - | - |
| 2024BVD17515 | Herbivore | European fallow deer | *Dama dama* | Vorpommern-Rügen | 102 | negative | - | - | - | negative | - | - |
| 2024BVD17516 | Herbivore | European fallow deer | *Dama dama* | Vorpommern-Rügen | 100 | negative | - | - | - | negative | - | - |
| 2024BVD17517 | Herbivore | Roe deer | *Capreolus capreolus* | Vorpommern-Rügen | 102 | negative | - | - | - | negative | - | - |
| 2024BVD17519 | Herbivore | European fallow deer | *Dama dama* | Vorpommern-Rügen | 102 | negative | - | - | - | negative | - | - |
| 2024BVD17520 | Omnivore | Wild boar | *Sus scrofa* | Vorpommern-Rügen | 95 | negative | - | - | - | negative | - | - |
| 2024BVD17521 | Omnivore | Wild boar | *Sus scrofa* | Vorpommern-Rügen | 94 | negative | - | - | - | negative | - | - |
| 2024BVD17522 | Herbivore | Red deer | *Cervus elaphus* | Vorpommern-Rügen | 98 | negative | - | - | - | negative | - | - |
| 2024BVD17523 | Omnivore | Wild boar | *Sus scrofa* | Vorpommern-Rügen | 100 | negative | - | - | - | negative | - | - |
| 2024BVD17524 | Herbivore | European fallow deer | *Dama dama* | Vorpommern-Rügen | 95 | negative | - | - | - | negative | - | - |
| 2024BVD17525 | Herbivore | European fallow deer | *Dama dama* | Vorpommern-Rügen | 100 | negative | - | - | - | negative | - | - |

| | | | | | | | | | | | | |
|---|---|---|---|---|---|---|---|---|---|---|---|---|
| 2024BVD17526 | Omnivore | Wild boar | *Sus scrofa* | Vorpommern-Rügen | 97 | negative | - | - | - | negative | - | - |
| 2024BVD17527 | Omnivore | Wild boar | *Sus scrofa* | Vorpommern-Rügen | 97 | negative | - | - | - | negative | - | - |
| 2024BVD17528 | Omnivore | Wild boar | *Sus scrofa* | Vorpommern-Rügen | 97 | negative | - | - | - | negative | - | - |
| 2024BVD17529 | Herbivore | Red deer | *Cervus elaphus* | Vorpommern-Rügen | 97 | negative | - | - | - | negative | - | - |
| 2024BVD17530 | Omnivore | Wild boar | *Sus scrofa* | Vorpommern-Rügen | 102 | negative | - | - | - | negative | - | - |
| 2024BVD17531 | Omnivore | Wild boar | *Sus scrofa* | Vorpommern-Rügen | 98 | negative | - | - | - | negative | - | - |
| 2024BVD17532 | Omnivore | Wild boar | *Sus scrofa* | Vorpommern-Rügen | 92 | negative | - | - | - | negative | - | - |
| 2024BVD17533 | Omnivore | Wild boar | *Sus scrofa* | Vorpommern-Rügen | 102 | negative | - | - | - | negative | - | - |
| 2024BVD17534 | Omnivore | Wild boar | *Sus scrofa* | Vorpommern-Rügen | 93 | negative | - | - | - | negative | - | - |
| 2024BVD17535 | Herbivore | Red deer | *Cervus elaphus* | Vorpommern-Rügen | 98 | negative | - | - | - | negative | - | - |
| 2024BVD17536 | Omnivore | Wild boar | *Sus scrofa* | Vorpommern-Rügen | 94 | negative | - | - | - | negative | - | - |
| 2024BVD17537 | Herbivore | Red deer | *Cervus elaphus* | Vorpommern-Rügen | 89 | negative | - | - | - | negative | - | - |
| 2024BVD17538 | Herbivore | Red deer | *Cervus elaphus* | Vorpommern-Rügen | 95 | negative | - | - | - | negative | - | - |
| 2024BVD17539 | Omnivore | Wild boar | *Sus scrofa* | Vorpommern-Rügen | 94 | negative | - | - | - | negative | - | - |
| 2024BVD17542 | Herbivore | Roe deer | *Capreolus capreolus* | Vorpommern-Greifswald | 90 | negative | - | - | - | negative | - | - |
| 2024BVD17543 | Herbivore | Roe deer | *Capreolus capreolus* | Vorpommern-Greifswald | 86 | negative | - | - | - | negative | - | - |
| 2024BVD17544 | Herbivore | European fallow deer | *Dama dama* | Vorpommern-Greifswald | 96 | negative | - | - | - | negative | - | - |
| 2024BVD17545 | Herbivore | Roe deer | *Capreolus capreolus* | Vorpommern-Greifswald | 91 | negative | - | - | - | negative | - | - |

| | | | | | | | | | | | | |
|---|---|---|---|---|---|---|---|---|---|---|---|---|
| 2024BVD17546 | Herbivore | Roe deer | *Capreolus capreolus* | Vorpommern-Greifswald | 92 | negative | - | - | - | negative | - | - |
| 2024BVD17547 | Herbivore | Roe deer | *Capreolus capreolus* | Vorpommern-Greifswald | 72 | negative | - | - | - | negative | - | - |
| 2024BVD17548 | Herbivore | European fallow deer | *Dama dama* | Vorpommern-Greifswald | 77 | negative | - | - | - | negative | - | - |
| 2025BVD00072 | Herbivore | Mouflon | *Ovis gmelini musimon* | Vorpommern-Rügen | 95 | negative | - | - | - | not taken | - | - |
| 2025BVD00073 | Herbivore | Mouflon | *Ovis gmelini musimon* | Vorpommern-Rügen | 96 | negative | - | - | - | not taken | - | - |
| 2025BVD00074 | Herbivore | Mouflon | *Ovis gmelini musimon* | Vorpommern-Rügen | 96 | negative | - | - | - | not taken | - | - |
| 2025BVD00075 | Herbivore | Mouflon | *Ovis gmelini musimon* | Vorpommern-Rügen | 94 | negative | - | - | - | not taken | - | - |
| 2025BVD00076 | Herbivore | Mouflon | *Ovis gmelini musimon* | Vorpommern-Rügen | 93 | negative | - | - | - | not taken | - | - |
| 2025BVD00846 | Herbivore | Mouflon | *Ovis gmelini musimon* | Vorpommern-Rügen | 96 | negative | - | - | - | not taken | - | - |
| 2023BVD08586 | Carnivore | Red fox | *Vulpes vulpes* | Vorpommern-Rügen | 26 | positive | 44 | positive | - | negative | negative | negative |
| 2023BVD08587 | Carnivore | Red fox | *Vulpes vulpes* | Vorpommern-Rügen | 97 | negative | - | - | - | negative | negative | negative |
| 2023BVD08588 | Carnivore | Raccoon | *Procyon lotor* | Vorpommern-Rügen | 104 | negative | - | - | - | negative | negative | negative |
| 2023BVD08589 | Carnivore | Red fox | *Vulpes vulpes* | Vorpommern-Rügen | 82 | negative | - | - | - | negative | negative | negative |
| 2023BVD08590 | Carnivore | Red fox | *Vulpes vulpes* | Vorpommern-Rügen | 48 | undetermined | 41 | positive | - | negative | negative | negative |
| 2023BVD08591 | Carnivore | Red fox | *Vulpes vulpes* | Vorpommern-Rügen | 29 | positive | 48 | positive | - | negative | negative | negative |
| 2023BVD08592 | Carnivore | Red fox | *Vulpes vulpes* | Vorpommern-Rügen | 67 | negative | - | - | - | negative | negative | negative |
| 2023BVD08593 | Carnivore | Raccoon dog | *Nyctereutes procyonoides* | Vorpommern-Rügen | 90 | negative | - | - | - | negative | negative | negative |
| 2023BVD08594 | Carnivore | Red fox | *Vulpes vulpes* | Vorpommern-Rügen | 98 | negative | - | - | - | negative | negative | negative |

| | | | | | | | | | | | | |
|---|---|---|---|---|---|---|---|---|---|---|---|---|
| 2023BVD08595 | Carnivore | Raccoon dog | *Nyctereutes procyonoides* | Vorpommern-Rügen | 73 | negative | - | - | - | negative | negative | negative |
| 2023BVD08596 | Carnivore | Raccoon dog | *Nyctereutes procyonoides* | Vorpommern-Rügen | 86 | negative | - | - | - | negative | negative | negative |
| 2023BVD08597 | Carnivore | Red fox | *Vulpes vulpes* | Vorpommern-Rügen | 79 | negative | - | - | - | negative | negative | negative |
| 2023BVD08598 | Carnivore | Red fox | *Vulpes vulpes* | Vorpommern-Rügen | 80 | negative | - | - | - | negative | negative | negative |
| 2023BVD08599 | Carnivore | Red fox | *Vulpes vulpes* | Vorpommern-Rügen | not taken | not taken | not taken | not taken | - | negative | negative | negative |
| 2023BVD08600 | Carnivore | Red fox | *Vulpes vulpes* | Vorpommern-Rügen | 86 | negative | - | - | - | negative | negative | negative |
| 2023BVD08601 | Carnivore | Raccoon | *Procyon lotor* | Vorpommern-Rügen | 61 | negative | - | - | - | negative | negative | negative |
| 2023BVD08602 | Carnivore | Raccoon dog | *Nyctereutes procyonoides* | Vorpommern-Rügen | 41 | positive | 37 | positive | - | negative | negative | negative |
| 2023BVD08603 | Carnivore | Red fox | *Vulpes vulpes* | Vorpommern-Rügen | 74 | negative | - | - | - | negative | negative | not taken |
| 2023BVD08604 | Carnivore | Red fox | *Vulpes vulpes* | Vorpommern-Rügen | 90 | negative | - | - | - | negative | negative | negative |
| 2023BVD08605 | Carnivore | Red fox | *Vulpes vulpes* | Vorpommern-Rügen | 78 | negative | - | - | - | negative | negative | negative |
| 2023BVD08606 | Carnivore | Red fox | *Vulpes vulpes* | Vorpommern-Rügen | 86 | negative | - | - | - | negative | negative | negative |
| 2023BVD08607 | Carnivore | Red fox | *Vulpes vulpes* | Vorpommern-Rügen | 86 | negative | - | - | - | negative | negative | negative |
| 2023BVD08608 | Carnivore | Raccoon | *Procyon lotor* | Vorpommern-Rügen | 80 | negative | - | - | - | negative | negative | negative |
| 2023BVD08609 | Carnivore | Raccoon dog | *Nyctereutes procyonoides* | Vorpommern-Rügen | 113 | negative | - | - | - | negative | negative | negative |
| 2023BVD08610 | Carnivore | Raccoon | *Procyon lotor* | Vorpommern-Rügen | 92 | negative | - | - | - | negative | negative | negative |
| 2023BVD08611 | Carnivore | Red fox | *Vulpes vulpes* | Vorpommern-Rügen | 96 | negative | - | - | - | negative | negative | negative |
| 2023BVD08694 | Carnivore | Red fox | *Vulpes vulpes* | Vorpommern-Rügen | 97 | negative | - | - | - | negative | negative | negative |
| 2023BVD08695 | Carnivore | Red fox | *Vulpes vulpes* | Vorpommern-Rügen | 74 | negative | - | - | - | negative | negative | negative |

| | | | | | | | | | | | | |
|---|---|---|---|---|---|---|---|---|---|---|---|---|
| 2023BVD08696 | Carnivore | Red fox | *Vulpes vulpes* | Vorpommern-Rügen | 89 | negative | - | - | - | negative | negative | negative |
| 2023BVD08697 | Carnivore | Red fox | *Vulpes vulpes* | Vorpommern-Rügen | 36 | positive | not available | not available | - | negative | negative | negative |
| 2023BVD08698 | Carnivore | Raccoon | *Procyon lotor* | Vorpommern-Rügen | 91 | negative | - | - | - | negative | negative | negative |
| 2023BVD08699 | Carnivore | Red fox | *Vulpes vulpes* | Vorpommern-Rügen | 99 | negative | - | - | - | negative | negative | negative |
| 2023BVD08700 | Carnivore | Red fox | *Vulpes vulpes* | Vorpommern-Rügen | 93 | negative | - | - | - | negative | negative | negative |
| 2023BVD08701 | Carnivore | Red fox | *Vulpes vulpes* | Vorpommern-Rügen | 72 | negative | - | - | - | negative | negative | negative |
| 2023BVD08702 | Carnivore | Red fox | *Vulpes vulpes* | Vorpommern-Rügen | 19 | positive | 7 | positive | - | negative | negative | negative |
| 2023BVD08703 | Carnivore | Red fox | *Vulpes vulpes* | Vorpommern-Rügen | 84 | negative | - | - | - | negative | negative | negative |
| 2023BVD08704 | Carnivore | Red fox | *Vulpes vulpes* | Vorpommern-Rügen | 91 | negative | - | - | - | negative | negative | negative |
| 2023BVD08705 | Carnivore | Red fox | *Vulpes vulpes* | Vorpommern-Rügen | 82 | negative | - | - | - | negative | negative | negative |
| 2023BVD08706 | Carnivore | Red fox | *Vulpes vulpes* | Vorpommern-Rügen | 85 | negative | - | - | - | negative | negative | negative |
| 2023BVD08707 | Carnivore | Red fox | *Vulpes vulpes* | Vorpommern-Rügen | 89 | negative | - | - | - | negative | negative | negative |
| 2023BVD08708 | Carnivore | Red fox | *Vulpes vulpes* | Vorpommern-Rügen | 38 | positive | 70 | negative | - | negative | negative | negative |
| 2023BVD09100 | Carnivore | Red fox | *Vulpes vulpes* | Vorpommern-Rügen | 23 | positive | 48 | positive | - | negative | negative | negative |
| 2024BVD00105 | Carnivore | Raccoon dog | *Nyctereutes procyonoides* | Vorpommern-Rügen | 90 | negative | - | - | - | negative | negative | negative |
| 2024BVD00106 | Carnivore | Red fox | *Vulpes vulpes* | Vorpommern-Rügen | 92 | negative | - | - | - | negative | negative | negative |
| 2024BVD00107 | Carnivore | Raccoon dog | *Nyctereutes procyonoides* | Vorpommern-Rügen | 84 | negative | - | - | - | negative | negative | negative |
| 2024BVD00108 | Carnivore | Red fox | *Vulpes vulpes* | Vorpommern-Rügen | 11 | positive | 11 | positive | - | negative | negative | negative |
| 2024BVD00109 | Carnivore | Red fox | *Vulpes vulpes* | Vorpommern-Rügen | 88 | negative | - | - | - | negative | negative | negative |

| | | | | | | | | | | | | |
|---|---|---|---|---|---|---|---|---|---|---|---|---|
| 2024BVD00110 | Carnivore | Red fox | *Vulpes vulpes* | Vorpommern-Rügen | 64 | negative | - | - | - | negative | negative | negative |
| 2024BVD00111 | Carnivore | Raccoon dog | *Nyctereutes procyonoides* | Vorpommern-Rügen | 92 | negative | - | - | - | negative | negative | negative |
| 2024BVD00112 | Carnivore | Red fox | *Vulpes vulpes* | Vorpommern-Rügen | 90 | negative | - | - | - | negative | negative | negative |
| 2024BVD00113 | Carnivore | Raccoon | *Procyon lotor* | Vorpommern-Rügen | 96 | negative | - | - | - | negative | negative | negative |
| 2024BVD00114 | Carnivore | Red fox | *Vulpes vulpes* | Vorpommern-Rügen | 90 | negative | - | - | - | negative | negative | negative |
| 2024BVD00115 | Carnivore | Raccoon | *Procyon lotor* | Vorpommern-Rügen | 106 | negative | - | - | - | negative | negative | negative |
| 2024BVD00116 | Carnivore | Red fox | *Vulpes vulpes* | Vorpommern-Rügen | 89 | negative | - | - | - | negative | negative | negative |
| 2024BVD00117 | Carnivore | Red fox | *Vulpes vulpes* | Vorpommern-Rügen | not available | not available | not available | not available | - | negative | negative | negative |
| 2024BVD00118 | Carnivore | Red fox | *Vulpes vulpes* | Vorpommern-Rügen | 20 | positive | 41 | positive | - | negative | negative | not taken |
| 2024BVD00119 | Carnivore | Red fox | *Vulpes vulpes* | Vorpommern-Rügen | 89 | negative | - | - | - | negative | negative | negative |
| 2024BVD00120 | Carnivore | Red fox | *Vulpes vulpes* | Vorpommern-Rügen | 81 | negative | - | - | - | negative | negative | negative |
| 2024BVD00121 | Carnivore | Red fox | *Vulpes vulpes* | Vorpommern-Rügen | 81 | negative | - | - | - | negative | negative | negative |
| 2024BVD00122 | Carnivore | Red fox | *Vulpes vulpes* | Vorpommern-Rügen | 90 | negative | - | - | - | negative | negative | negative |
| 2024BVD00123 | Carnivore | Raccoon dog | *Nyctereutes procyonoides* | Vorpommern-Rügen | 109 | negative | - | - | - | negative | negative | negative |
| 2024BVD00124 | Carnivore | Red fox | *Vulpes vulpes* | Vorpommern-Rügen | 92 | negative | - | - | - | negative | negative | negative |
| 2024BVD00125 | Carnivore | Red fox | *Vulpes vulpes* | Vorpommern-Rügen | 59 | negative | - | - | - | negative | negative | negative |
| 2024BVD00127 | Carnivore | Red fox | *Vulpes vulpes* | Vorpommern-Rügen | 16 | positive | 8 | positive | - | negative | negative | negative |
| 2024BVD00128 | Carnivore | Raccoon dog | *Nyctereutes procyonoides* | Vorpommern-Rügen | 83 | negative | - | - | - | negative | negative | negative |
| 2024BVD00129 | Carnivore | Red fox | *Vulpes vulpes* | Vorpommern-Rügen | 95 | negative | - | - | - | negative | negative | negative |

| | | | | | | | | | | | | |
|---|---|---|---|---|---|---|---|---|---|---|---|---|
| 2024BVD00130 | Carnivore | Red fox | *Vulpes vulpes* | Vorpommern-Rügen | 33 | positive | 5 | positive | - | negative | negative | negative |
| 2024BVD00131 | Carnivore | Red fox | *Vulpes vulpes* | Vorpommern-Rügen | 33 | positive | 67 | negative | - | negative | negative | negative |
| 2024BVD00132 | Carnivore | Red fox | *Vulpes vulpes* | Vorpommern-Rügen | 23 | positive | 5 | positive | - | negative | negative | negative |
| 2024BVD00133 | Carnivore | Raccoon dog | *Nyctereutes procyonoides* | Vorpommern-Rügen | 91 | negative | - | - | - | negative | negative | negative |
| 2024BVD00134 | Carnivore | Raccoon dog | *Nyctereutes procyonoides* | Vorpommern-Rügen | 74 | negative | - | - | - | negative | negative | negative |
| 2024BVD00135 | Carnivore | Red fox | *Vulpes vulpes* | Vorpommern-Rügen | 79 | negative | - | - | - | negative | negative | negative |
| 2024BVD00136 | Carnivore | Raccoon dog | *Nyctereutes procyonoides* | Vorpommern-Rügen | 35 | positive | 37 | positive | - | negative | negative | not taken |
| 2024BVD00137 | Carnivore | Red fox | *Vulpes vulpes* | Vorpommern-Rügen | 82 | negative | - | - | - | negative | negative | negative |
| 2024BVD00138 | Carnivore | Raccoon dog | *Nyctereutes procyonoides* | Vorpommern-Rügen | 87 | negative | - | - | - | negative | negative | negative |
| 2024BVD00139 | Carnivore | Red fox | *Vulpes vulpes* | Vorpommern-Rügen | 34 | positive | 26 | positive | - | negative | negative | negative |
| 2024BVD00140 | Carnivore | Raccoon dog | *Nyctereutes procyonoides* | Vorpommern-Rügen | 89 | negative | - | - | - | negative | negative | negative |
| 2024BVD00141 | Carnivore | Red fox | *Vulpes vulpes* | Vorpommern-Rügen | 48 | undetermined | 55 | undetermined | - | negative | negative | negative |
| 2024BVD00142 | Carnivore | Red fox | *Vulpes vulpes* | Vorpommern-Rügen | 17 | positive | 68 | negative | - | negative | negative | negative |
| 2024BVD00143 | Carnivore | Red fox | *Vulpes vulpes* | Vorpommern-Rügen | 36 | positive | 12 | positive | - | negative | negative | negative |
| 2024BVD00144 | Carnivore | Red fox | *Vulpes vulpes* | Vorpommern-Rügen | 98 | negative | - | - | - | negative | negative | negative |
| 2024BVD00145 | Carnivore | Red fox | *Vulpes vulpes* | Vorpommern-Rügen | 86 | negative | - | - | - | negative | negative | negative |
| 2024BVD00146 | Carnivore | Red fox | *Vulpes vulpes* | Vorpommern-Rügen | 94 | negative | - | - | - | negative | negative | negative |
| 2024BVD00147 | Carnivore | Red fox | *Vulpes vulpes* | Vorpommern-Rügen | 100 | negative | - | - | - | negative | negative | negative |
| 2024BVD00148 | Carnivore | Red fox | *Vulpes vulpes* | Vorpommern-Rügen | 11 | positive | 21 | positive | - | not taken | negative | negative |

| | | | | | | | | | | | | |
|---|---|---|---|---|---|---|---|---|---|---|---|---|
| 2024BVD00149 | Carnivore | Red fox | *Vulpes vulpes* | Vorpommern-Rügen | 12 | positive | 14 | positive | - | not taken | negative | negative |
| 2024BVD00150 | Carnivore | Red fox | *Vulpes vulpes* | Vorpommern-Rügen | 105 | negative | - | - | - | not taken | negative | negative |
| 2024BVD00151 | Carnivore | Red fox | *Vulpes vulpes* | Vorpommern-Rügen | 106 | negative | - | - | - | not taken | negative | negative |
| 2024BVD02899 | Carnivore | Red fox | *Vulpes vulpes* | Vorpommern-Rügen | 10 | positive | 27 | positive | - | negative | negative | negative |
| 2024BVD02900 | Carnivore | Raccoon | *Procyon lotor* | Vorpommern-Rügen | 83 | negative | - | - | - | negative | negative | negative |
| 2024BVD02901 | Carnivore | Red fox | *Vulpes vulpes* | Vorpommern-Rügen | 81 | negative | - | - | - | negative | negative | negative |
| 2024BVD02902 | Carnivore | Raccoon dog | *Nyctereutes procyonoides* | Vorpommern-Rügen | 67 | negative | - | - | - | negative | negative | negative |
| 2024BVD02903 | Carnivore | Red fox | *Vulpes vulpes* | Vorpommern-Rügen | 40 | positive | 69 | negative | - | negative | negative | negative |
| 2024BVD02904 | Carnivore | Red fox | *Vulpes vulpes* | Vorpommern-Rügen | 5 | positive | 40 | positive | - | negative | negative | negative |
| 2024BVD02905 | Carnivore | Raccoon dog | *Nyctereutes procyonoides* | Vorpommern-Rügen | not taken | not taken | not taken | not taken | - | negative | negative | negative |
| 2024BVD02906 | Carnivore | Red fox | *Vulpes vulpes* | Vorpommern-Rügen | 90 | negative | - | - | - | negative | negative | negative |
| 2024BVD02907 | Carnivore | Red fox | *Vulpes vulpes* | Vorpommern-Rügen | 99 | negative | - | - | - | negative | negative | negative |
| 2024BVD02908 | Carnivore | Red fox | *Vulpes vulpes* | Vorpommern-Rügen | 107 | negative | - | - | - | negative | negative | negative |
| 2024BVD02909 | Carnivore | Raccoon dog | *Nyctereutes procyonoides* | Vorpommern-Rügen | 96 | negative | - | - | - | negative | negative | negative |
| 2024BVD02910 | Carnivore | Red fox | *Vulpes vulpes* | Vorpommern-Rügen | 10 | positive | 46 | positive | 1:40 | negative | negative | negative |
| 2024BVD02911 | Carnivore | Raccoon dog | *Nyctereutes procyonoides* | Vorpommern-Rügen | 87 | negative | - | - | - | negative | negative | negative |
| 2024BVD02912 | Carnivore | Red fox | *Vulpes vulpes* | Vorpommern-Rügen | 9 | positive | 6 | positive | - | negative | negative | negative |
| 2024BVD02913 | Carnivore | Raccoon dog | *Nyctereutes procyonoides* | Vorpommern-Rügen | 94 | negative | - | - | - | negative | negative | negative |
| 2024BVD02914 | Carnivore | Red fox | *Vulpes vulpes* | Vorpommern-Rügen | not taken | not taken | not taken | not taken | - | negative | negative | negative |

| | | | | | | | | | | | | |
|---|---|---|---|---|---|---|---|---|---|---|---|---|
| 2024BVD02915 | Carnivore | Red fox | *Vulpes vulpes* | Vorpommern-Rügen | 73 | negative | - | - | - | negative | negative | negative |
| 2024BVD02916 | Carnivore | Red fox | *Vulpes vulpes* | Vorpommern-Rügen | 76 | negative | - | - | - | negative | negative | negative |
| 2024BVD02917 | Carnivore | Red fox | *Vulpes vulpes* | Vorpommern-Rügen | 68 | negative | - | - | - | negative | negative | negative |
| 2024BVD02918 | Carnivore | Red fox | *Vulpes vulpes* | Vorpommern-Rügen | 88 | negative | - | - | - | negative | negative | negative |
| 2024BVD02919 | Carnivore | Red fox | *Vulpes vulpes* | Vorpommern-Rügen | 10 | positive | 5 | positive | - | negative | negative | negative |
| 2024BVD02920 | Carnivore | Red fox | *Vulpes vulpes* | Vorpommern-Rügen | 94 | negative | - | - | - | negative | negative | negative |
| 2024BVD02921 | Carnivore | Raccoon dog | *Nyctereutes procyonoides* | Vorpommern-Rügen | 90 | negative | - | - | - | negative | negative | negative |
| 2024BVD02922 | Carnivore | Red fox | *Vulpes vulpes* | Vorpommern-Rügen | 88 | negative | - | - | - | negative | negative | negative |
| 2024BVD02923 | Carnivore | Red fox | *Vulpes vulpes* | Vorpommern-Rügen | 15 | positive | 31 | positive | - | negative | negative | not taken |
| 2024BVD03122 | Carnivore | Raccoon dog | *Nyctereutes procyonoides* | Vorpommern-Rügen | 92 | negative | - | - | - | negative | negative | negative |
| 2024BVD03123 | Carnivore | Red fox | *Vulpes vulpes* | Vorpommern-Rügen | 86 | negative | - | - | - | negative | negative | negative |
| 2024BVD03124 | Carnivore | Red fox | *Vulpes vulpes* | Vorpommern-Rügen | 7 | positive | 20 | positive | - | negative | negative | negative |
| 2024BVD03125 | Carnivore | Red fox | *Vulpes vulpes* | Vorpommern-Rügen | 86 | negative | - | - | - | negative | negative | negative |
| 2024BVD03126 | Carnivore | Red fox | *Vulpes vulpes* | Vorpommern-Rügen | 54 | negative | - | - | - | negative | negative | negative |
| 2024BVD03127 | Carnivore | Raccoon | *Procyon lotor* | Vorpommern-Rügen | 6 | positive | 41 | positive | - | negative | negative | negative |
| 2024BVD03128 | Carnivore | Raccoon dog | *Nyctereutes procyonoides* | Vorpommern-Rügen | 91 | negative | - | - | - | negative | negative | negative |
| 2024BVD03129 | Carnivore | Red fox | *Vulpes vulpes* | Vorpommern-Rügen | 91 | negative | - | - | - | negative | negative | negative |
| 2024BVD03130 | Carnivore | Red fox | *Vulpes vulpes* | Vorpommern-Rügen | 91 | negative | - | - | - | negative | negative | negative |
| 2024BVD03131 | Carnivore | Red fox | *Vulpes vulpes* | Vorpommern-Rügen | 83 | negative | - | - | - | negative | negative | negative |

| | | | | | | | | | | | | |
|---|---|---|---|---|---|---|---|---|---|---|---|---|
| 2024BVD03132 | Carnivore | Red fox | *Vulpes vulpes* | Vorpommern-Rügen | 13 | positive | 7 | positive | - | negative | negative | negative |
| 2024BVD03133 | Carnivore | Raccoon dog | *Nyctereutes procyonoides* | Vorpommern-Rügen | 50 | negative | - | - | - | negative | negative | negative |
| 2024BVD03134 | Carnivore | Red fox | *Vulpes vulpes* | Vorpommern-Rügen | 60 | negative | - | - | - | negative | negative | negative |
| 2024BVD03135 | Carnivore | Red fox | *Vulpes vulpes* | Vorpommern-Rügen | 84 | negative | - | - | - | negative | negative | negative |
| 2024BVD03136 | Carnivore | Red fox | *Vulpes vulpes* | Vorpommern-Rügen | 91 | negative | - | - | - | negative | negative | negative |
| 2024BVD03137 | Carnivore | Red fox | *Vulpes vulpes* | Vorpommern-Rügen | 95 | negative | - | - | - | negative | negative | negative |
| 2024BVD03138 | Carnivore | Red fox | *Vulpes vulpes* | Vorpommern-Rügen | 92 | negative | - | - | - | negative | negative | negative |
| 2024BVD03139 | Carnivore | Red fox | *Vulpes vulpes* | Vorpommern-Rügen | 101 | negative | - | - | - | negative | negative | negative |
| 2024BVD03140 | Carnivore | Red fox | *Vulpes vulpes* | Vorpommern-Rügen | 94 | negative | - | - | - | negative | negative | negative |
| 2024BVD03141 | Carnivore | European badger | *Meles meles* | Vorpommern-Rügen | 80 | negative | - | - | - | negative | negative | negative |
| 2024BVD03142 | Carnivore | Red fox | *Vulpes vulpes* | Vorpommern-Rügen | 105 | negative | - | - | - | negative | negative | negative |
| 2024BVD03143 | Carnivore | Red fox | *Vulpes vulpes* | Vorpommern-Rügen | 80 | negative | - | - | - | negative | negative | negative |
| 2024BVD03144 | Carnivore | Red fox | *Vulpes vulpes* | Vorpommern-Rügen | 84 | negative | - | - | - | negative | negative | negative |
| 2024BVD03145 | Carnivore | Red fox | *Vulpes vulpes* | Vorpommern-Rügen | 6 | positive | 6 | positive | - | negative | negative | negative |
| 2024BVD03146 | Carnivore | Raccoon dog | *Nyctereutes procyonoides* | Vorpommern-Rügen | 100 | negative | - | - | - | negative | negative | negative |
| 2024BVD03207 | Carnivore | Raccoon dog | *Nyctereutes procyonoides* | Vorpommern-Rügen | 24 | positive | 29 | positive | - | negative | negative | negative |
| 2024BVD03208 | Carnivore | Red fox | *Vulpes vulpes* | Vorpommern-Rügen | 15 | positive | 7 | positive | - | negative | negative | negative |
| 2024BVD03209 | Carnivore | Red fox | *Vulpes vulpes* | Vorpommern-Rügen | 70 | negative | - | - | - | negative | negative | negative |
| 2024BVD03210 | Carnivore | Red fox | *Vulpes vulpes* | Vorpommern-Rügen | 99 | negative | - | - | - | negative | negative | negative |

| | | | | | | | | | | | | |
|---|---|---|---|---|---|---|---|---|---|---|---|---|
| 2024BVD03211 | Carnivore | Red fox | *Vulpes vulpes* | Vorpommern-Rügen | 106 | negative | - | - | - | negative | negative | negative |
| 2024BVD03212 | Carnivore | Red fox | *Vulpes vulpes* | Vorpommern-Rügen | 88 | negative | - | - | - | negative | negative | negative |
| 2024BVD03213 | Carnivore | Raccoon dog | *Nyctereutes procyonoides* | Vorpommern-Rügen | 95 | negative | - | - | - | negative | negative | negative |
| 2024BVD03214 | Carnivore | Red fox | *Vulpes vulpes* | Vorpommern-Rügen | 91 | negative | - | - | - | negative | negative | negative |
| 2024BVD03215 | Carnivore | Red fox | *Vulpes vulpes* | Vorpommern-Rügen | 83 | negative | - | - | - | negative | negative | negative |
| 2024BVD03216 | Carnivore | Red fox | *Vulpes vulpes* | Vorpommern-Rügen | 85 | negative | - | - | - | negative | negative | negative |
| 2024BVD03217 | Carnivore | Red fox | *Vulpes vulpes* | Vorpommern-Rügen | 102 | negative | - | - | - | negative | negative | negative |
| 2024BVD03218 | Carnivore | Red fox | *Vulpes vulpes* | Vorpommern-Rügen | 86 | negative | - | - | - | negative | negative | negative |
| 2024BVD03219 | Carnivore | Red fox | *Vulpes vulpes* | Vorpommern-Rügen | 90 | negative | - | - | - | negative | negative | negative |
| 2024BVD03220 | Carnivore | Red fox | *Vulpes vulpes* | Vorpommern-Rügen | 107 | negative | - | - | - | negative | negative | negative |
| 2024BVD03221 | Carnivore | Red fox | *Vulpes vulpes* | Vorpommern-Rügen | 91 | negative | - | - | - | negative | negative | negative |
| 2024BVD03222 | Carnivore | Red fox | *Vulpes vulpes* | Vorpommern-Rügen | 49 | undetermined | 18 | positive | - | negative | negative | negative |
| 2024BVD03223 | Carnivore | Red fox | *Vulpes vulpes* | Vorpommern-Rügen | 94 | negative | - | - | - | negative | negative | negative |
| 2024BVD03224 | Carnivore | Red fox | *Vulpes vulpes* | Vorpommern-Rügen | 98 | negative | - | - | - | negative | negative | negative |
| 2024BVD03225 | Carnivore | Red fox | *Vulpes vulpes* | Vorpommern-Rügen | 101 | negative | - | - | - | negative | negative | negative |
| 2024BVD03226 | Carnivore | Red fox | *Vulpes vulpes* | Vorpommern-Rügen | 9 | positive | 48 | positive | 1:60 | negative | negative | negative |
| 2024BVD03227 | Carnivore | Red fox | *Vulpes vulpes* | Vorpommern-Rügen | 91 | negative | - | - | - | negative | negative | negative |
| 2024BVD03228 | Carnivore | Red fox | *Vulpes vulpes* | Vorpommern-Rügen | 86 | negative | - | - | - | negative | negative | negative |
| 2024BVD03229 | Carnivore | Red fox | *Vulpes vulpes* | Vorpommern-Rügen | 5 | positive | 27 | positive | - | negative | negative | negative |

| | | | | | | | | | | | | |
|---|---|---|---|---|---|---|---|---|---|---|---|---|
| 2024BVD03230 | Carnivore | Raccoon dog | *Nyctereutes procyonoides* | Vorpommern-Rügen | 66 | negative | - | - | - | negative | negative | negative |
| 2024BVD03231 | Carnivore | Raccoon | *Procyon lotor* | Vorpommern-Rügen | 96 | negative | - | - | - | negative | negative | negative |
| 2024BVD03232 | Carnivore | Red fox | *Vulpes vulpes* | Vorpommern-Rügen | 97 | negative | - | - | - | negative | negative | negative |
| 2024BVD03233 | Carnivore | Raccoon dog | *Nyctereutes procyonoides* | Vorpommern-Rügen | 82 | negative | - | - | - | negative | negative | negative |
| 2024BVD03234 | Carnivore | Red fox | *Vulpes vulpes* | Vorpommern-Rügen | 45 | undetermined | 15 | positive | - | negative | negative | negative |
| 2024BVD03235 | Carnivore | Raccoon | *Procyon lotor* | Vorpommern-Rügen | 8 | positive | 36 | positive | - | negative | negative | negative |
| 2024BVD03236 | Carnivore | Red fox | *Vulpes vulpes* | Vorpommern-Rügen | 83 | negative | - | - | - | negative | negative | negative |
| 2024BVD04004 | Carnivore | Raccoon dog | *Nyctereutes procyonoides* | Vorpommern-Rügen | 32 | positive | 22 | positive | - | negative | negative | negative |
| 2024BVD04005 | Carnivore | Red fox | *Vulpes vulpes* | Vorpommern-Rügen | 115 | negative | - | - | - | negative | negative | negative |
| 2024BVD04006 | Carnivore | Raccoon | *Procyon lotor* | Vorpommern-Rügen | 8 | positive | 35 | positive | - | negative | negative | negative |
| 2024BVD04007 | Carnivore | European badger | *Meles meles* | Vorpommern-Rügen | 96 | negative | - | - | - | negative | negative | negative |
| 2024BVD04008 | Carnivore | Raccoon dog | *Nyctereutes procyonoides* | Vorpommern-Rügen | 13 | positive | 50 | positive | - | negative | negative | negative |
| 2024BVD04009 | Carnivore | Raccoon dog | *Nyctereutes procyonoides* | Vorpommern-Rügen | 81 | negative | - | - | - | negative | negative | negative |
| 2024BVD04010 | Carnivore | Red fox | *Vulpes vulpes* | Vorpommern-Rügen | 9 | positive | 12 | positive | - | negative | negative | negative |
| 2024BVD04011 | Carnivore | Red fox | *Vulpes vulpes* | Vorpommern-Rügen | 51 | negative | - | - | - | negative | negative | negative |
| 2024BVD04012 | Carnivore | Red fox | *Vulpes vulpes* | Vorpommern-Rügen | 97 | negative | - | - | - | negative | negative | negative |
| 2024BVD04013 | Carnivore | Red fox | *Vulpes vulpes* | Vorpommern-Rügen | 15 | positive | 5 | positive | - | negative | negative | negative |
| 2024BVD04014 | Carnivore | Red fox | *Vulpes vulpes* | Vorpommern-Rügen | 91 | negative | - | - | - | negative | negative | negative |
| 2024BVD04015 | Carnivore | Red fox | *Vulpes vulpes* | Vorpommern-Rügen | 85 | negative | - | - | - | negative | negative | negative |

| | | | | | | | | | | | | |
|---|---|---|---|---|---|---|---|---|---|---|---|---|
| 2024BVD04016 | Carnivore | Red fox | *Vulpes vulpes* | Vorpommern-Rügen | 14 | positive | 48 | positive | - | negative | negative | negative |
| 2024BVD04017 | Carnivore | Red fox | *Vulpes vulpes* | Vorpommern-Rügen | 7 | positive | 13 | positive | - | negative | negative | negative |
| 2024BVD04018 | Carnivore | Red fox | *Vulpes vulpes* | Vorpommern-Rügen | 6 | positive | 22 | positive | - | negative | negative | negative |
| 2024BVD04019 | Carnivore | Red fox | *Vulpes vulpes* | Vorpommern-Rügen | 56 | negative | - | - | - | negative | negative | negative |
| 2024BVD04020 | Carnivore | Raccoon | *Procyon lotor* | Vorpommern-Rügen | 4 | positive | 32 | positive | - | negative | negative | negative |
| 2024BVD04021 | Carnivore | Red fox | *Vulpes vulpes* | Vorpommern-Rügen | 86 | negative | - | - | - | negative | negative | negative |
| 2024BVD04022 | Carnivore | Raccoon dog | *Nyctereutes procyonoides* | Vorpommern-Rügen | 84 | negative | - | - | - | negative | negative | negative |
| 2024BVD04023 | Carnivore | Raccoon dog | *Nyctereutes procyonoides* | Vorpommern-Rügen | 75 | negative | - | - | - | negative | negative | negative |
| 2024BVD04024 | Carnivore | Red fox | *Vulpes vulpes* | Vorpommern-Rügen | not taken | not taken | not taken | not taken | - | negative | negative | negative |
| 2024BVD04025 | Carnivore | Red fox | *Vulpes vulpes* | Vorpommern-Rügen | 94 | negative | - | - | - | negative | negative | negative |
| 2024BVD04026 | Carnivore | Red fox | *Vulpes vulpes* | Vorpommern-Rügen | not taken | not taken | not taken | not taken | - | negative | negative | negative |
| 2024BVD04027 | Carnivore | Red fox | *Vulpes vulpes* | Vorpommern-Rügen | 90 | negative | - | - | - | negative | negative | negative |
| 2024BVD04028 | Carnivore | Red fox | *Vulpes vulpes* | Vorpommern-Rügen | not taken | not taken | not taken | not taken | - | negative | negative | negative |
| 2024BVD04029 | Carnivore | Red fox | *Vulpes vulpes* | Vorpommern-Rügen | 88 | negative | - | - | - | negative | negative | negative |
| 2024BVD04030 | Carnivore | Red fox | *Vulpes vulpes* | Vorpommern-Rügen | 90 | negative | - | - | - | negative | negative | negative |
| 2024BVD04031 | Carnivore | Red fox | *Vulpes vulpes* | Vorpommern-Rügen | 29 | positive | 10 | positive | - | negative | negative | negative |
| 2024BVD04032 | Carnivore | Red fox | *Vulpes vulpes* | Vorpommern-Rügen | not taken | not taken | not taken | not taken | - | negative | negative | negative |
| 2024BVD04033 | Carnivore | Red fox | *Vulpes vulpes* | Vorpommern-Rügen | 89 | negative | - | - | - | negative | negative | negative |
| 2024BVD04034 | Carnivore | Red fox | *Vulpes vulpes* | Vorpommern-Rügen | 86 | negative | - | - | - | negative | negative | negative |

| | | | | | | | | | | | | |
|---|---|---|---|---|---|---|---|---|---|---|---|---|
| 2024BVD04035 | Carnivore | Raccoon dog | *Nyctereutes procyonoides* | Vorpommern-Rügen | 5 | positive | 40 | positive | - | negative | negative | negative |
| 2024BVD04036 | Carnivore | European badger | *Meles meles* | Vorpommern-Rügen | 16 | positive | 5 | positive | - | negative | negative | negative |
| 2024BVD04037 | Carnivore | Raccoon dog | *Nyctereutes procyonoides* | Vorpommern-Rügen | 57 | negative | - | - | - | negative | negative | negative |
| 2024BVD04038 | Carnivore | Raccoon dog | *Nyctereutes procyonoides* | Vorpommern-Rügen | 38 | positive | 31 | positive | - | negative | negative | negative |
| 2024BVD08288 | Carnivore | Raccoon dog | *Nyctereutes procyonoides* | Vorpommern-Rügen | 90 | negative | - | - | - | negative | negative | negative |
| 2024BVD10141 | Carnivore | Raccoon dog | *Nyctereutes procyonoides* | Vorpommern-Rügen | 83 | negative | - | - | - | negative | negative | negative |
| 2024BVD10142 | Carnivore | Red fox | *Vulpes vulpes* | Vorpommern-Rügen | not taken | not taken | not taken | not taken | - | negative | negative | negative |
| 2024BVD10143 | Carnivore | Raccoon dog | *Nyctereutes procyonoides* | Vorpommern-Rügen | 86 | negative | - | - | - | negative | negative | negative |
| 2024BVD10144 | Carnivore | Red fox | *Vulpes vulpes* | Vorpommern-Rügen | 79 | negative | - | - | - | negative | negative | negative |
| 2024BVD10145 | Carnivore | Red fox | *Vulpes vulpes* | Vorpommern-Rügen | 58 | negative | - | - | - | negative | negative | negative |
| 2024BVD10146 | Carnivore | Red fox | *Vulpes vulpes* | Vorpommern-Rügen | 86 | negative | - | - | - | negative | negative | negative |
| 2024BVD10147 | Carnivore | Red fox | *Vulpes vulpes* | Vorpommern-Rügen | 110 | negative | - | - | - | negative | negative | not taken |
| 2024BVD10148 | Carnivore | Red fox | *Vulpes vulpes* | Vorpommern-Rügen | 87 | negative | - | - | - | negative | negative | negative |
| 2024BVD10149 | Carnivore | Red fox | *Vulpes vulpes* | Vorpommern-Rügen | 95 | negative | - | - | - | negative | negative | negative |
| 2024BVD10150 | Carnivore | Red fox | *Vulpes vulpes* | Vorpommern-Rügen | 80 | negative | - | - | - | negative | negative | negative |
| 2024BVD10151 | Carnivore | Red fox | *Vulpes vulpes* | Vorpommern-Rügen | 79 | negative | - | - | - | negative | negative | negative |
| 2024BVD10152 | Carnivore | Red fox | *Vulpes vulpes* | Vorpommern-Rügen | 93 | negative | - | - | - | negative | negative | negative |
| 2024BVD10153 | Carnivore | Red fox | *Vulpes vulpes* | Vorpommern-Rügen | 83 | negative | - | - | - | negative | negative | negative |
| 2024BVD10154 | Carnivore | Red fox | *Vulpes vulpes* | Vorpommern-Rügen | 88 | negative | - | - | - | negative | negative | negative |

| | | | | | | | | | | | | |
|---|---|---|---|---|---|---|---|---|---|---|---|---|
| 2024BVD10155 | Carnivore | Red fox | *Vulpes vulpes* | Vorpommern-Rügen | 44 | positive | 77 | negative | - | negative | negative | negative |
| 2024BVD10156 | Carnivore | Red fox | *Vulpes vulpes* | Vorpommern-Rügen | 77 | negative | - | - | - | negative | negative | negative |
| 2024BVD10157 | Carnivore | Red fox | *Vulpes vulpes* | Vorpommern-Rügen | 84 | negative | - | - | - | negative | negative | negative |
| 2024BVD10158 | Carnivore | Raccoon dog | *Nyctereutes procyonoides* | Vorpommern-Rügen | 60 | negative | - | - | - | negative | negative | negative |
| 2024BVD10159 | Carnivore | Red fox | *Vulpes vulpes* | Vorpommern-Rügen | 25 | positive | 94 | negative | <1:20 | negative | negative | negative |
| 2024BVD10160 | Carnivore | Red fox | *Vulpes vulpes* | Vorpommern-Rügen | 68 | negative | - | - | - | negative | negative | negative |
| 2024BVD10161 | Carnivore | Red fox | *Vulpes vulpes* | Vorpommern-Rügen | 45 | undetermined | 105 | negative | - | negative | negative | negative |
| 2024BVD10162 | Carnivore | Raccoon dog | *Nyctereutes procyonoides* | Vorpommern-Rügen | 87 | negative | - | - | - | negative | negative | negative |
| 2024BVD10163 | Carnivore | Red fox | *Vulpes vulpes* | Vorpommern-Rügen | 17 | positive | 81 | negative | <1:20 | negative | negative | negative |
| 2024BVD10164 | Carnivore | Red fox | *Vulpes vulpes* | Vorpommern-Rügen | 28 | positive | 81 | negative | <1:20 | negative | negative | negative |
| 2024BVD10165 | Carnivore | Red fox | *Vulpes vulpes* | Vorpommern-Rügen | 68 | negative | - | - | - | negative | negative | negative |
| 2024BVD12400 | Carnivore | Red fox | *Vulpes vulpes* | Vorpommern-Rügen | 29 | positive | 88 | negative | - | negative | negative | negative |
| 2024BVD12401 | Carnivore | Red fox | *Vulpes vulpes* | Vorpommern-Rügen | 41 | positive | 87 | negative | <1:20 | negative | negative | negative |
| 2024BVD12402 | Carnivore | Red fox | *Vulpes vulpes* | Vorpommern-Rügen | 60 | negative | - | - | - | negative | negative | negative |
| 2024BVD12502 | Carnivore | Red fox | *Vulpes vulpes* | Vorpommern-Rügen | 47 | undetermined | not available | not available | - | negative | negative | negative |
| 2024BVD12503 | Carnivore | Red fox | *Vulpes vulpes* | Vorpommern-Rügen | not taken | not taken | not taken | not taken | - | negative | negative | negative |
| 2024BVD12504 | Carnivore | Raccoon dog | *Nyctereutes procyonoides* | Vorpommern-Rügen | 13 | positive | 5 | positive | - | negative | negative | negative |
| 2024BVD12505 | Carnivore | Raccoon dog | *Nyctereutes procyonoides* | Vorpommern-Rügen | 92 | negative | - | - | - | negative | negative | negative |
| 2024BVD12506 | Carnivore | Raccoon dog | *Nyctereutes procyonoides* | Vorpommern-Rügen | 78 | negative | - | - | - | negative | not taken | negative |

| | | | | | | | | | | | | |
|---|---|---|---|---|---|---|---|---|---|---|---|---|
| 2024BVD12507 | Carnivore | European badger | *Meles meles* | Vorpommern-Rügen | 103 | negative | - | - | - | negative | negative | not taken |
| 2024BVD12508 | Carnivore | Red fox | *Vulpes vulpes* | Vorpommern-Rügen | 92 | negative | - | - | - | negative | negative | negative |
| 2024BVD12509 | Carnivore | Red fox | *Vulpes vulpes* | Vorpommern-Rügen | 98 | negative | - | - | - | negative | negative | negative |
| 2024BVD12510 | Carnivore | Red fox | *Vulpes vulpes* | Vorpommern-Rügen | 99 | negative | - | - | - | negative | negative | negative |
| 2024BVD12511 | Carnivore | Red fox | *Vulpes vulpes* | Vorpommern-Rügen | 85 | negative | - | - | - | negative | negative | negative |
| 2024BVD12512 | Carnivore | Red fox | *Vulpes vulpes* | Vorpommern-Rügen | 107 | negative | - | - | - | negative | negative | negative |
| 2024BVD12513 | Carnivore | Red fox | *Vulpes vulpes* | Vorpommern-Rügen | 92 | negative | - | - | - | negative | negative | negative |
| 2024BVD12514 | Carnivore | Red fox | *Vulpes vulpes* | Vorpommern-Rügen | 70 | negative | - | - | - | negative | negative | negative |
| 2024BVD12515 | Carnivore | Red fox | *Vulpes vulpes* | Vorpommern-Rügen | 94 | negative | - | - | - | negative | negative | negative |
| 2024BVD12516 | Carnivore | Red fox | *Vulpes vulpes* | Vorpommern-Rügen | not available | not available | not available | not available | - | negative | negative | negative |
| 2024BVD12517 | Carnivore | Red fox | *Vulpes vulpes* | Vorpommern-Rügen | 91 | negative | - | - | - | negative | negative | negative |
| 2024BVD12518 | Carnivore | Red fox | *Vulpes vulpes* | Vorpommern-Rügen | not taken | not taken | not taken | not taken | - | negative | negative | negative |
| 2024BVD12519 | Carnivore | Raccoon dog | *Nyctereutes procyonoides* | Vorpommern-Rügen | 86 | negative | - | - | - | negative | negative | negative |
| 2024BVD12520 | Carnivore | Red fox | *Vulpes vulpes* | Vorpommern-Rügen | 103 | negative | - | - | - | negative | negative | negative |
| 2024BVD12521 | Carnivore | Red fox | *Vulpes vulpes* | Vorpommern-Rügen | 72 | negative | - | - | - | negative | negative | negative |
| 2024BVD12522 | Carnivore | Red fox | *Vulpes vulpes* | Vorpommern-Rügen | not taken | not taken | not taken | not taken | - | negative | not taken | not taken |
| 2024BVD12523 | Carnivore | Red fox | *Vulpes vulpes* | Vorpommern-Rügen | 98 | negative | - | - | - | negative | negative | negative |
| 2024BVD12524 | Carnivore | Red fox | *Vulpes vulpes* | Vorpommern-Rügen | 89 | negative | - | - | - | negative | negative | negative |
| 2024BVD12525 | Carnivore | Red fox | *Vulpes vulpes* | Vorpommern-Rügen | 87 | negative | - | - | - | negative | negative | negative |

| | | | | | | | | | | | | |
|---|---|---|---|---|---|---|---|---|---|---|---|---|
| 2024BVD12526 | Carnivore | Raccoon dog | *Nyctereutes procyonoides* | Vorpommern-Rügen | not available | not available | not available | not available | - | negative | negative | negative |
| 2024BVD12527 | Carnivore | Raccoon dog | *Nyctereutes procyonoides* | Vorpommern-Rügen | 96 | negative | - | - | - | negative | negative | negative |
| 2024BVD12528 | Carnivore | Raccoon dog | *Nyctereutes procyonoides* | Vorpommern-Rügen | 101 | negative | - | - | - | negative | negative | negative |
| 2024BVD12529 | Carnivore | Raccoon dog | *Nyctereutes procyonoides* | Vorpommern-Rügen | 96 | negative | - | - | - | negative | negative | negative |
| 2024BVD12530 | Carnivore | Raccoon dog | *Nyctereutes procyonoides* | Vorpommern-Rügen | 87 | negative | - | - | - | negative | negative | negative |
| 2024BVD12531 | Carnivore | Raccoon dog | *Nyctereutes procyonoides* | Vorpommern-Rügen | 82 | negative | - | - | - | negative | negative | negative |
| 2024BVD12532 | Carnivore | Raccoon dog | *Nyctereutes procyonoides* | Vorpommern-Rügen | 83 | negative | - | - | - | negative | negative | negative |
| 2024BVD12533 | Carnivore | Red fox | *Vulpes vulpes* | Vorpommern-Rügen | 81 | negative | - | - | - | negative | negative | negative |
| 2024BVD12534 | Carnivore | Red fox | *Vulpes vulpes* | Vorpommern-Rügen | 80 | negative | - | - | - | negative | negative | negative |
| 2024BVD12535 | Carnivore | Red fox | *Vulpes vulpes* | Vorpommern-Rügen | 70 | negative | - | - | - | negative | negative | negative |
| 2024BVD12536 | Carnivore | Raccoon dog | *Nyctereutes procyonoides* | Vorpommern-Rügen | 94 | negative | - | - | - | negative | negative | negative |
| 2024BVD12537 | Carnivore | Raccoon dog | *Nyctereutes procyonoides* | Vorpommern-Rügen | 95 | negative | - | - | - | negative | negative | negative |
| 2024BVD12538 | Carnivore | Red fox | *Vulpes vulpes* | Vorpommern-Rügen | 79 | negative | - | - | - | negative | negative | negative |
| 2024BVD12539 | Carnivore | Red fox | *Vulpes vulpes* | Vorpommern-Rügen | 95 | negative | - | - | - | negative | negative | negative |
| 2024BVD12540 | Carnivore | Raccoon dog | *Nyctereutes procyonoides* | Vorpommern-Rügen | 102 | negative | - | - | - | negative | negative | negative |
| 2024BVD12541 | Carnivore | Raccoon dog | *Nyctereutes procyonoides* | Vorpommern-Rügen | 91 | negative | - | - | - | negative | negative | negative |
| 2024BVD12542 | Carnivore | Raccoon dog | *Nyctereutes procyonoides* | Vorpommern-Rügen | 83 | negative | - | - | - | negative | negative | negative |
| 2024BVD12543 | Carnivore | Red fox | *Vulpes vulpes* | Vorpommern-Rügen | 11 | positive | 94 | negative | - | negative | negative | negative |
| 2024BVD12544 | Carnivore | Red fox | *Vulpes vulpes* | Vorpommern-Rügen | 2 | positive | 95 | negative | - | negative | negative | negative |

| | | | | | | | | | | | | |
|---|---|---|---|---|---|---|---|---|---|---|---|---|
| 2024BVD12545 | Carnivore | Red fox | *Vulpes vulpes* | Vorpommern-Rügen | 52 | negative | - | - | - | negative | negative | negative |
| 2024BVD12546 | Carnivore | Red fox | *Vulpes vulpes* | Vorpommern-Rügen | 41 | positive | 114 | negative | <1:20 | negative | negative | negative |
| 2024BVD12547 | Carnivore | Red fox | *Vulpes vulpes* | Vorpommern-Rügen | 66 | negative | - | - | - | negative | negative | not taken |
| 2024BVD12548 | Carnivore | Raccoon dog | *Nyctereutes procyonoides* | Vorpommern-Rügen | 73 | negative | - | - | - | negative | negative | negative |
| 2024BVD12549 | Carnivore | Red fox | *Vulpes vulpes* | Vorpommern-Rügen | 59 | negative | - | - | - | negative | negative | negative |
| 2024BVD12550 | Carnivore | Red fox | *Vulpes vulpes* | Vorpommern-Rügen | 83 | negative | - | - | - | negative | negative | negative |
| 2024BVD12551 | Carnivore | Red fox | *Vulpes vulpes* | Vorpommern-Rügen | 80 | negative | - | - | - | negative | negative | negative |
| 2024BVD12657 | Carnivore | Red fox | *Vulpes vulpes* | Vorpommern-Rügen | 95 | negative | - | - | - | negative | negative | negative |
| 2024BVD12658 | Carnivore | Red fox | *Vulpes vulpes* | Vorpommern-Rügen | 90 | negative | - | - | - | negative | negative | negative |
| 2024BVD12659 | Carnivore | Red fox | *Vulpes vulpes* | Vorpommern-Rügen | 97 | negative | - | - | - | negative | negative | negative |
| 2024BVD12660 | Carnivore | Raccoon dog | *Nyctereutes procyonoides* | Vorpommern-Rügen | 91 | negative | - | - | - | negative | negative | not taken |
| 2024BVD12661 | Carnivore | Red fox | *Vulpes vulpes* | Vorpommern-Rügen | 97 | negative | - | - | - | negative | negative | negative |
| 2024BVD12662 | Carnivore | Red fox | *Vulpes vulpes* | Vorpommern-Rügen | 94 | negative | - | - | - | negative | negative | negative |
| 2024BVD12663 | Carnivore | European pine marten | *Martes martes* | Vorpommern-Rügen | 91 | negative | - | - | - | negative | negative | negative |
| 2024BVD12664 | Carnivore | Red fox | *Vulpes vulpes* | Vorpommern-Rügen | 85 | negative | - | - | - | negative | negative | negative |
| 2024BVD12665 | Carnivore | Red fox | *Vulpes vulpes* | Vorpommern-Rügen | 22 | positive | 6 | positive | - | negative | negative | negative |
| 2024BVD12667 | Carnivore | Raccoon dog | *Nyctereutes procyonoides* | Vorpommern-Rügen | not taken | not taken | not taken | not taken | - | negative | negative | negative |
| 2024BVD12668 | Carnivore | Red fox | *Vulpes vulpes* | Vorpommern-Rügen | 75 | negative | - | - | - | negative | negative | negative |

| | | | | | | | | | | | | |
|---|---|---|---|---|---|---|---|---|---|---|---|---|
| 2024BVD12669 | Carnivore | Red fox | *Vulpes vulpes* | Vorpommern-Rügen | 66 | negative | - | - | - | negative | negative | negative |
| 2024BVD12670 | Carnivore | Raccoon dog | *Nyctereutes procyonoides* | Vorpommern-Rügen | 64 | negative | - | - | - | negative | negative | negative |
| 2024BVD12671 | Carnivore | European badger | *Meles meles* | Vorpommern-Rügen | 75 | negative | - | - | - | negative | negative | negative |
| 2024BVD12672 | Carnivore | Raccoon dog | *Nyctereutes procyonoides* | Vorpommern-Rügen | 63 | negative | - | - | - | negative | negative | negative |
| 2024BVD12673 | Carnivore | Red fox | *Vulpes vulpes* | Vorpommern-Rügen | 91 | negative | - | - | - | negative | negative | negative |
| 2024BVD12674 | Carnivore | Raccoon dog | *Nyctereutes procyonoides* | Vorpommern-Rügen | 87 | negative | - | - | - | negative | negative | negative |
| 2024BVD12675 | Carnivore | Raccoon dog | *Nyctereutes procyonoides* | Vorpommern-Rügen | 97 | negative | - | - | - | negative | negative | negative |
| 2024BVD12676 | Carnivore | Red fox | *Vulpes vulpes* | Vorpommern-Rügen | 94 | negative | - | - | - | negative | negative | negative |
| 2024BVD12677 | Carnivore | Red fox | *Vulpes vulpes* | Vorpommern-Rügen | 83 | negative | - | - | - | negative | negative | negative |
| 2024BVD12678 | Carnivore | Raccoon dog | *Nyctereutes procyonoides* | Vorpommern-Rügen | 90 | negative | - | - | - | negative | negative | negative |
| 2024BVD12679 | Carnivore | Red fox | *Vulpes vulpes* | Vorpommern-Rügen | 88 | negative | - | - | - | negative | negative | negative |
| 2024BVD12680 | Carnivore | Red fox | *Vulpes vulpes* | Vorpommern-Rügen | 85 | negative | - | - | - | negative | negative | negative |
| 2024BVD12681 | Carnivore | Red fox | *Vulpes vulpes* | Vorpommern-Rügen | 47 | undetermined | not available | not available | - | negative | negative | negative |
| 2024BVD12682 | Carnivore | Raccoon | *Procyon lotor* | Vorpommern-Rügen | 95 | negative | - | - | - | negative | negative | negative |
| 2024BVD12683 | Carnivore | Raccoon dog | *Nyctereutes procyonoides* | Vorpommern-Rügen | not taken | not taken | not taken | not taken | - | negative | negative | negative |
| 2024BVD12684 | Carnivore | Red fox | *Vulpes vulpes* | Vorpommern-Rügen | 64 | negative | - | - | - | negative | negative | negative |
| 2024BVD12685 | Carnivore | Raccoon dog | *Nyctereutes procyonoides* | Vorpommern-Rügen | 89 | negative | - | - | - | negative | negative | negative |
| 2024BVD12686 | Carnivore | Raccoon dog | *Nyctereutes procyonoides* | Vorpommern-Rügen | 69 | negative | - | - | - | negative | negative | negative |
| 2024BVD12687 | Carnivore | Raccoon | *Procyon lotor* | Vorpommern-Rügen | 85 | negative | - | - | - | negative | negative | negative |

| | | | | | | | | | | | | |
|---|---|---|---|---|---|---|---|---|---|---|---|---|
| 2024BVD12688 | Carnivore | Raccoon dog | *Nyctereutes procyonoides* | Vorpommern-Rügen | 83 | negative | - | - | - | negative | negative | negative |
| 2024BVD12689 | Carnivore | Red fox | *Vulpes vulpes* | Vorpommern-Rügen | 91 | negative | - | - | - | negative | negative | negative |
| 2024BVD12690 | Carnivore | Red fox | *Vulpes vulpes* | Vorpommern-Rügen | 95 | negative | - | - | - | negative | negative | negative |
| 2024BVD12691 | Carnivore | Red fox | *Vulpes vulpes* | Vorpommern-Rügen | 96 | negative | - | - | - | negative | negative | negative |
| 2024BVD12692 | Carnivore | Red fox | *Vulpes vulpes* | Vorpommern-Rügen | 99 | negative | - | - | - | negative | negative | negative |
| 2024BVD12693 | Carnivore | Raccoon | *Procyon lotor* | Vorpommern-Rügen | 101 | negative | - | - | - | negative | negative | negative |
| 2024BVD12694 | Carnivore | Raccoon | *Procyon lotor* | Vorpommern-Rügen | 92 | negative | - | - | - | negative | negative | negative |
| 2024BVD12695 | Carnivore | Raccoon dog | *Nyctereutes procyonoides* | Vorpommern-Rügen | 87 | negative | - | - | - | negative | negative | negative |
| 2024BVD12696 | Carnivore | Red fox | *Vulpes vulpes* | Vorpommern-Rügen | 91 | negative | - | - | - | negative | negative | negative |
| 2024BVD12697 | Carnivore | Raccoon dog | *Nyctereutes procyonoides* | Vorpommern-Rügen | not taken | not taken | not taken | not taken | - | negative | negative | negative |
| 2024BVD12698 | Carnivore | Raccoon dog | *Nyctereutes procyonoides* | Vorpommern-Rügen | 90 | negative | - | - | - | negative | negative | negative |
| 2024BVD12699 | Carnivore | Red fox | *Vulpes vulpes* | Vorpommern-Rügen | 95 | negative | - | - | - | negative | negative | negative |
| 2024BVD12700 | Carnivore | Raccoon dog | *Nyctereutes procyonoides* | Vorpommern-Rügen | 87 | negative | - | - | - | negative | negative | negative |
| 2024BVD12701 | Carnivore | Raccoon dog | *Nyctereutes procyonoides* | Vorpommern-Rügen | 23 | positive | 17 | positive | - | negative | negative | negative |
| 2024BVD15064 | Carnivore | Raccoon dog | *Nyctereutes procyonoides* | Vorpommern-Rügen | 96 | negative | - | - | - | negative | negative | negative |
| 2024BVD15065 | Carnivore | Red fox | *Vulpes vulpes* | Vorpommern-Rügen | 82 | negative | - | - | - | negative | negative | negative |
| 2024BVD15066 | Carnivore | Raccoon dog | *Nyctereutes procyonoides* | Vorpommern-Rügen | 96 | negative | - | - | - | negative | negative | negative |
| 2024BVD15067 | Carnivore | Raccoon | *Procyon lotor* | Vorpommern-Rügen | 98 | negative | - | - | - | negative | negative | negative |
| 2024BVD15068 | Carnivore | Raccoon dog | *Nyctereutes procyonoides* | Vorpommern-Rügen | 10 | positive | 5 | positive | - | negative | negative | negative |

| | | | | | | | | | | | | |
|---|---|---|---|---|---|---|---|---|---|---|---|---|
| 2024BVD15069 | Carnivore | Raccoon | *Procyon lotor* | Vorpommern-Rügen | 89 | negative | - | - | - | negative | negative | negative |
| 2024BVD15070 | Carnivore | Raccoon dog | *Nyctereutes procyonoides* | Vorpommern-Rügen | 67 | negative | - | - | - | negative | negative | negative |
| 2024BVD15071 | Carnivore | Red fox | *Vulpes vulpes* | Vorpommern-Rügen | 97 | negative | - | - | - | negative | negative | negative |
| 2024BVD15072 | Carnivore | Raccoon dog | *Nyctereutes procyonoides* | Vorpommern-Rügen | 91 | negative | - | - | - | negative | negative | negative |
| 2024BVD15073 | Carnivore | Red fox | *Vulpes vulpes* | Vorpommern-Rügen | 57 | negative | - | - | - | negative | negative | negative |
| 2024BVD15074 | Carnivore | Raccoon dog | *Nyctereutes procyonoides* | Vorpommern-Rügen | 110 | negative | - | - | - | negative | negative | negative |
| 2024BVD15075 | Carnivore | Red fox | *Vulpes vulpes* | Vorpommern-Rügen | 100 | negative | - | - | - | negative | negative | negative |
| 2024BVD15076 | Carnivore | Raccoon dog | *Nyctereutes procyonoides* | Vorpommern-Rügen | 103 | negative | - | - | - | negative | negative | negative |
| 2024BVD15077 | Carnivore | Raccoon dog | *Nyctereutes procyonoides* | Vorpommern-Rügen | 98 | negative | - | - | - | negative | negative | negative |
| 2024BVD15078 | Carnivore | Raccoon | *Procyon lotor* | Vorpommern-Rügen | 92 | negative | - | - | - | negative | negative | not taken |
| 2024BVD15079 | Carnivore | Red fox | *Vulpes vulpes* | Vorpommern-Rügen | 99 | negative | - | - | - | negative | negative | negative |
| 2024BVD15080 | Carnivore | Raccoon | *Procyon lotor* | Vorpommern-Rügen | 11 | positive | 4 | positive | - | negative | negative | negative |
| 2024BVD15081 | Carnivore | Raccoon dog | *Nyctereutes procyonoides* | Vorpommern-Rügen | 105 | negative | - | - | - | negative | negative | negative |
| 2024BVD15082 | Carnivore | Red fox | *Vulpes vulpes* | Vorpommern-Rügen | 103 | negative | - | - | - | negative | negative | negative |
| 2024BVD15083 | Carnivore | Raccoon dog | *Nyctereutes procyonoides* | Vorpommern-Rügen | 103 | negative | - | - | - | negative | negative | negative |
| 2024BVD15084 | Carnivore | Raccoon dog | *Nyctereutes procyonoides* | Vorpommern-Rügen | 106 | negative | - | - | - | negative | negative | negative |
| 2024BVD15085 | Carnivore | Red fox | *Vulpes vulpes* | Vorpommern-Rügen | 75 | negative | - | - | - | negative | negative | negative |
| 2024BVD15086 | Carnivore | Red fox | *Vulpes vulpes* | Vorpommern-Rügen | 84 | negative | - | - | - | negative | negative | negative |
| 2024BVD15087 | Carnivore | Raccoon dog | *Nyctereutes procyonoides* | Vorpommern-Rügen | 82 | negative | - | - | - | negative | negative | negative |

| | | | | | | | | | | | | |
|---|---|---|---|---|---|---|---|---|---|---|---|---|
| 2024BVD15088 | Carnivore | Raccoon dog | *Nyctereutes procyonoides* | Vorpommern-Rügen | 95 | negative | - | - | - | negative | negative | negative |
| 2024BVD15089 | Carnivore | Raccoon dog | *Nyctereutes procyonoides* | Vorpommern-Rügen | 4 | positive | 5 | positive | - | negative | negative | negative |
| 2024BVD15090 | Carnivore | Raccoon dog | *Nyctereutes procyonoides* | Vorpommern-Rügen | 101 | negative | - | - | - | negative | negative | negative |
| 2024BVD15091 | Carnivore | Raccoon dog | *Nyctereutes procyonoides* | Vorpommern-Rügen | 86 | negative | - | - | - | negative | negative | negative |
| 2024BVD15092 | Carnivore | Raccoon dog | *Nyctereutes procyonoides* | Vorpommern-Rügen | 100 | negative | - | - | - | negative | negative | negative |
| 2024BVD15093 | Carnivore | Raccoon dog | *Nyctereutes procyonoides* | Vorpommern-Rügen | 95 | negative | - | - | - | negative | negative | negative |
| 2024BVD15094 | Carnivore | Red fox | *Vulpes vulpes* | Vorpommern-Rügen | 90 | negative | - | - | - | negative | negative | negative |
| 2024BVD15095 | Carnivore | Raccoon dog | *Nyctereutes procyonoides* | Vorpommern-Rügen | 83 | negative | - | - | - | negative | negative | negative |
| 2024BVD15096 | Carnivore | Raccoon dog | *Nyctereutes procyonoides* | Vorpommern-Rügen | 10 | positive | 9 | positive | - | negative | negative | negative |
| 2024BVD15097 | Carnivore | Raccoon dog | *Nyctereutes procyonoides* | Vorpommern-Rügen | 91 | negative | - | - | - | negative | negative | negative |
| 2024BVD15098 | Carnivore | Red fox | *Vulpes vulpes* | Vorpommern-Rügen | 97 | negative | - | - | - | negative | negative | negative |
| 2024BVD15099 | Carnivore | Raccoon dog | *Nyctereutes procyonoides* | Vorpommern-Rügen | 89 | negative | - | - | - | negative | negative | negative |
| 2024BVD15100 | Carnivore | Red fox | *Vulpes vulpes* | Vorpommern-Rügen | 102 | negative | - | - | - | negative | negative | negative |
| 2024BVD15101 | Carnivore | Red fox | *Vulpes vulpes* | Vorpommern-Rügen | 5 | positive | 6 | positive | - | negative | negative | negative |
| 2024BVD15102 | Carnivore | Red fox | *Vulpes vulpes* | Vorpommern-Rügen | 108 | negative | - | - | - | negative | negative | negative |
| 2024BVD15103 | Carnivore | Red fox | *Vulpes vulpes* | Vorpommern-Rügen | 103 | negative | - | - | - | negative | negative | negative |
| 2024BVD15104 | Carnivore | Raccoon dog | *Nyctereutes procyonoides* | Vorpommern-Rügen | 96 | negative | - | - | - | negative | negative | negative |
| 2024BVD15105 | Carnivore | Raccoon | *Procyon lotor* | Vorpommern-Rügen | 106 | negative | - | - | - | negative | negative | negative |
| 2024BVD15106 | Carnivore | Raccoon | *Procyon lotor* | Vorpommern-Rügen | 36 | positive | 93 | negative | <1:20 | negative | negative | negative |

| | | | | | | | | | | | | |
|---|---|---|---|---|---|---|---|---|---|---|---|---|
| 2024BVD15107 | Carnivore | Raccoon | *Procyon lotor* | Vorpommern-Rügen | 80 | negative | - | - | - | negative | negative | negative |
| 2024BVD15108 | Carnivore | Red fox | *Vulpes vulpes* | Vorpommern-Rügen | 21 | positive | 61 | negative | <1:20 | negative | negative | negative |
| 2024BVD16173 | Carnivore | Raccoon | *Procyon lotor* | Vorpommern-Rügen | 30 | positive | 99 | negative | - | negative | negative | negative |
| 2024BVD16174 | Carnivore | Red fox | *Vulpes vulpes* | Vorpommern-Rügen | 4 | positive | 80 | negative | - | negative | negative | negative |
| 2024BVD16175 | Carnivore | Raccoon dog | *Nyctereutes procyonoides* | Vorpommern-Rügen | 87 | negative | - | - | - | negative | negative | negative |
| 2024BVD16176 | Carnivore | Red fox | *Vulpes vulpes* | Vorpommern-Rügen | 52 | negative | - | - | - | negative | negative | negative |
| 2024BVD16177 | Carnivore | Red fox | *Vulpes vulpes* | Vorpommern-Rügen | 16 | positive | 32 | positive | - | negative | negative | negative |
| 2024BVD16178 | Carnivore | Raccoon dog | *Nyctereutes procyonoides* | Vorpommern-Rügen | 94 | negative | - | - | - | negative | not taken | negative |
| 2024BVD16179 | Carnivore | Raccoon dog | *Nyctereutes procyonoides* | Vorpommern-Rügen | 96 | negative | - | - | - | negative | negative | negative |
| 2024BVD16180 | Carnivore | Raccoon dog | *Nyctereutes procyonoides* | Vorpommern-Rügen | 93 | negative | - | - | - | negative | negative | negative |
| 2024BVD16181 | Carnivore | Raccoon | *Procyon lotor* | Vorpommern-Rügen | 12 | positive | 5 | positive | - | negative | negative | negative |
| 2024BVD16182 | Carnivore | Raccoon dog | *Nyctereutes procyonoides* | Vorpommern-Rügen | 86 | negative | - | - | - | negative | negative | negative |
| 2024BVD16183 | Carnivore | Raccoon dog | *Nyctereutes procyonoides* | Vorpommern-Rügen | 85 | negative | - | - | - | negative | negative | negative |
| 2024BVD16184 | Carnivore | Raccoon dog | *Nyctereutes procyonoides* | Vorpommern-Rügen | 92 | negative | - | - | - | negative | negative | negative |
| 2024BVD16185 | Carnivore | Raccoon dog | *Nyctereutes procyonoides* | Vorpommern-Rügen | 90 | negative | - | - | - | negative | negative | negative |
| 2024BVD16186 | Carnivore | Red fox | *Vulpes vulpes* | Vorpommern-Rügen | 79 | negative | - | - | - | negative | negative | negative |
| 2024BVD16187 | Carnivore | European badger | *Meles meles* | Vorpommern-Rügen | 83 | negative | - | - | - | negative | negative | negative |
| 2024BVD16188 | Carnivore | Raccoon dog | *Nyctereutes procyonoides* | Vorpommern-Rügen | 88 | negative | - | - | - | negative | negative | negative |
| 2024BVD16189 | Carnivore | Raccoon | *Procyon lotor* | Vorpommern-Rügen | 4 | positive | 4 | positive | ≥1:160 | negative | negative | negative |

| | | | | | | | | | | | | |
|---|---|---|---|---|---|---|---|---|---|---|---|---|
| 2024BVD16190 | Carnivore | Red fox | *Vulpes vulpes* | Vorpommern-Rügen | 31 | positive | 5 | positive | - | negative | negative | negative |
| 2024BVD16191 | Carnivore | Raccoon dog | *Nyctereutes procyonoides* | Vorpommern-Rügen | 102 | negative | - | - | - | negative | negative | negative |
| 2024BVD16192 | Carnivore | Raccoon | *Procyon lotor* | Vorpommern-Rügen | 89 | negative | - | - | - | negative | negative | negative |
| 2024BVD16193 | Carnivore | Raccoon dog | *Nyctereutes procyonoides* | Vorpommern-Rügen | 83 | negative | - | - | - | negative | negative | negative |
| 2024BVD16194 | Carnivore | Raccoon dog | *Nyctereutes procyonoides* | Vorpommern-Rügen | 83 | negative | - | - | - | negative | negative | negative |
| 2024BVD16195 | Carnivore | Red fox | *Vulpes vulpes* | Vorpommern-Rügen | 44 | positive | 5 | positive | - | negative | negative | negative |
| 2024BVD16196 | Carnivore | European badger | *Meles meles* | Vorpommern-Rügen | 85 | negative | - | - | - | negative | negative | negative |
| 2024BVD16197 | Carnivore | Red fox | *Vulpes vulpes* | Vorpommern-Rügen | 77 | negative | - | - | - | negative | negative | negative |
| 2024BVD16198 | Carnivore | Red fox | *Vulpes vulpes* | Vorpommern-Rügen | 79 | negative | - | - | - | negative | negative | negative |
| 2024BVD16199 | Carnivore | Raccoon dog | *Nyctereutes procyonoides* | Vorpommern-Rügen | 5 | positive | 5 | positive | - | negative | negative | negative |
| 2024BVD16200 | Carnivore | Raccoon dog | *Nyctereutes procyonoides* | Vorpommern-Rügen | 89 | negative | - | - | - | negative | negative | negative |
| 2024BVD16201 | Carnivore | Raccoon dog | *Nyctereutes procyonoides* | Vorpommern-Rügen | 103 | negative | - | - | - | negative | negative | negative |
| 2024BVD16202 | Carnivore | Raccoon | *Procyon lotor* | Vorpommern-Rügen | 13 | positive | 50 | undetermined | - | negative | negative | negative |
| 2024BVD16203 | Carnivore | Red fox | *Vulpes vulpes* | Vorpommern-Rügen | 104 | negative | - | - | - | negative | negative | negative |
| 2024BVD16204 | Carnivore | Red fox | *Vulpes vulpes* | Vorpommern-Rügen | 99 | negative | - | - | - | negative | negative | negative |
| 2024BVD16205 | Carnivore | Raccoon dog | *Nyctereutes procyonoides* | Vorpommern-Rügen | 97 | negative | - | - | - | negative | negative | negative |
| 2024BVD16206 | Carnivore | Raccoon dog | *Nyctereutes procyonoides* | Vorpommern-Rügen | 91 | negative | - | - | - | negative | negative | negative |
| 2024BVD16207 | Carnivore | Red fox | *Vulpes vulpes* | Vorpommern-Rügen | 39 | positive | 17 | positive | - | negative | negative | negative |
| 2024BVD16208 | Carnivore | Red fox | *Vulpes vulpes* | Vorpommern-Rügen | 81 | negative | - | - | - | negative | not taken | negative |

| | | | | | | | | | | | | |
|---|---|---|---|---|---|---|---|---|---|---|---|---|
| 2024BVD16209 | Carnivore | Raccoon | *Procyon lotor* | Vorpommern-Rügen | 111 | negative | - | - | - | negative | negative | negative |
| 2024BVD16210 | Carnivore | Raccoon dog | *Nyctereutes procyonoides* | Vorpommern-Rügen | 98 | negative | - | - | - | negative | negative | negative |
| 2024BVD16211 | Carnivore | Raccoon dog | *Nyctereutes procyonoides* | Vorpommern-Rügen | 93 | negative | - | - | - | negative | negative | negative |
| 2024BVD16212 | Carnivore | Red fox | *Vulpes vulpes* | Vorpommern-Rügen | 88 | negative | - | - | - | negative | negative | negative |
| 2024BVD16213 | Carnivore | Raccoon dog | *Nyctereutes procyonoides* | Vorpommern-Rügen | 96 | negative | - | - | - | negative | negative | negative |
| 2024BVD16214 | Carnivore | Raccoon dog | *Nyctereutes procyonoides* | Vorpommern-Rügen | 58 | negative | - | - | - | negative | negative | negative |
| 2024BVD16215 | Carnivore | Raccoon dog | *Nyctereutes procyonoides* | Vorpommern-Rügen | 81 | negative | - | - | - | negative | negative | negative |
| 2024BVD16216 | Carnivore | Raccoon dog | *Nyctereutes procyonoides* | Vorpommern-Rügen | 10 | positive | 12 | positive | - | negative | negative | not taken |
| 2024BVD16217 | Carnivore | Raccoon dog | *Nyctereutes procyonoides* | Vorpommern-Rügen | 102 | negative | - | - | - | negative | negative | negative |
| 2024BVD16218 | Carnivore | Red fox | *Vulpes vulpes* | Vorpommern-Rügen | 84 | negative | - | - | - | negative | negative | negative |
| 2024BVD16219 | Carnivore | Red fox | *Vulpes vulpes* | Vorpommern-Rügen | 97 | negative | - | - | - | negative | negative | negative |
| 2024BVD16220 | Carnivore | Raccoon | *Procyon lotor* | Vorpommern-Rügen | 4 | positive | 5 | positive | - | negative | negative | negative |
| 2024BVD16221 | Carnivore | Red fox | *Vulpes vulpes* | Vorpommern-Rügen | 98 | negative | - | - | - | negative | negative | negative |
| 2024BVD16222 | Carnivore | Raccoon dog | *Nyctereutes procyonoides* | Vorpommern-Rügen | 27 | positive | 76 | negative | - | negative | negative | negative |
| 2024BVD16376 | Carnivore | Red fox | *Vulpes vulpes* | Vorpommern-Rügen | 58 | negative | - | - | - | negative | negative | negative |
| 2024BVD16377 | Carnivore | Raccoon | *Procyon lotor* | Vorpommern-Rügen | 90 | negative | - | - | - | negative | negative | negative |
| 2024BVD16378 | Carnivore | Red fox | *Vulpes vulpes* | Vorpommern-Rügen | 60 | negative | - | - | - | negative | negative | negative |
| 2024BVD16379 | Carnivore | Red fox | *Vulpes vulpes* | Vorpommern-Rügen | 73 | negative | - | - | - | negative | negative | negative |
| 2024BVD16380 | Carnivore | Raccoon dog | *Nyctereutes procyonoides* | Vorpommern-Rügen | 101 | negative | - | - | - | negative | negative | negative |

| | | | | | | | | | | | | |
|---|---|---|---|---|---|---|---|---|---|---|---|---|
| 2024BVD16381 | Carnivore | Raccoon dog | *Nyctereutes procyonoides* | Vorpommern-Rügen | 91 | negative | - | - | - | negative | negative | negative |
| 2024BVD16382 | Carnivore | Red fox | *Vulpes vulpes* | Vorpommern-Rügen | 85 | negative | - | - | - | negative | negative | negative |
| 2024BVD16383 | Carnivore | Raccoon dog | *Nyctereutes procyonoides* | Vorpommern-Rügen | 93 | negative | - | - | - | negative | negative | not taken |
| 2024BVD16384 | Carnivore | Raccoon dog | *Nyctereutes procyonoides* | Vorpommern-Rügen | not taken | not taken | not taken | not taken | - | negative | negative | negative |
| 2024BVD16385 | Carnivore | Raccoon dog | *Nyctereutes procyonoides* | Vorpommern-Rügen | 107 | negative | - | - | - | negative | negative | negative |
| 2024BVD16386 | Carnivore | Red fox | *Vulpes vulpes* | Vorpommern-Rügen | 52 | negative | - | - | - | negative | negative | negative |
| 2024BVD16387 | Carnivore | Raccoon dog | *Nyctereutes procyonoides* | Vorpommern-Rügen | not taken | not taken | not taken | not taken | - | negative | negative | negative |
| 2024BVD16388 | Carnivore | Raccoon dog | *Nyctereutes procyonoides* | Vorpommern-Rügen | 6 | positive | 5 | positive | - | negative | negative | negative |
| 2024BVD16389 | Carnivore | Raccoon dog | *Nyctereutes procyonoides* | Vorpommern-Rügen | 101 | negative | - | - | - | negative | negative | negative |
| 2024BVD16390 | Carnivore | Raccoon dog | *Nyctereutes procyonoides* | Vorpommern-Rügen | 62 | negative | - | - | - | negative | negative | negative |
| 2024BVD16391 | Carnivore | Red fox | *Vulpes vulpes* | Vorpommern-Rügen | 13 | positive | 5 | positive | - | negative | negative | negative |
| 2024BVD16392 | Carnivore | Raccoon | *Procyon lotor* | Vorpommern-Rügen | 105 | negative | - | - | - | negative | negative | negative |
| 2024BVD16393 | Carnivore | Red fox | *Vulpes vulpes* | Vorpommern-Rügen | 69 | negative | - | - | - | negative | negative | negative |
| 2024BVD16394 | Carnivore | Raccoon dog | *Nyctereutes procyonoides* | Vorpommern-Rügen | 107 | negative | - | - | - | negative | negative | negative |
| 2024BVD16395 | Carnivore | Red fox | *Vulpes vulpes* | Vorpommern-Rügen | 86 | negative | - | - | - | negative | negative | negative |
| 2024BVD16396 | Carnivore | Raccoon dog | *Nyctereutes procyonoides* | Vorpommern-Rügen | 7 | positive | 6 | positive | - | negative | negative | negative |
| 2024BVD16397 | Carnivore | Raccoon | *Procyon lotor* | Vorpommern-Rügen | not taken | not taken | not taken | not taken | - | negative | negative | negative |
| 2024BVD16398 | Carnivore | Red fox | *Vulpes vulpes* | Vorpommern-Rügen | 82 | negative | - | - | - | negative | negative | negative |
| 2024BVD16399 | Carnivore | Raccoon dog | *Nyctereutes procyonoides* | Vorpommern-Rügen | 93 | negative | - | - | - | negative | negative | negative |

| | | | | | | | | | | | | |
|---|---|---|---|---|---|---|---|---|---|---|---|---|
| 2024BVD16400 | Carnivore | Red fox | *Vulpes vulpes* | Vorpommern-Rügen | 100 | negative | - | - | - | negative | negative | negative |
| 2024BVD16401 | Carnivore | Raccoon dog | *Nyctereutes procyonoides* | Vorpommern-Rügen | 99 | negative | - | - | - | negative | negative | negative |
| 2024BVD16402 | Carnivore | Red fox | *Vulpes vulpes* | Vorpommern-Rügen | 77 | negative | - | - | - | negative | negative | negative |
| 2024BVD16403 | Carnivore | Red fox | *Vulpes vulpes* | Vorpommern-Rügen | 79 | negative | - | - | - | negative | negative | negative |
| 2024BVD16404 | Carnivore | Raccoon dog | *Nyctereutes procyonoides* | Vorpommern-Rügen | 81 | negative | - | - | - | negative | negative | negative |
| 2024BVD16405 | Carnivore | Red fox | *Vulpes vulpes* | Vorpommern-Rügen | 95 | negative | - | - | - | negative | negative | negative |
| 2024BVD16406 | Carnivore | Raccoon dog | *Nyctereutes procyonoides* | Vorpommern-Rügen | 94 | negative | - | - | - | negative | negative | negative |
| 2024BVD16407 | Carnivore | Red fox | *Vulpes vulpes* | Vorpommern-Rügen | 105 | negative | - | - | - | negative | negative | negative |
| 2024BVD16408 | Carnivore | Raccoon dog | *Nyctereutes procyonoides* | Vorpommern-Rügen | 93 | negative | - | - | - | negative | negative | negative |
| 2024BVD16409 | Carnivore | Red fox | *Vulpes vulpes* | Vorpommern-Rügen | 104 | negative | - | - | - | negative | negative | negative |
| 2024BVD16410 | Carnivore | Red fox | *Vulpes vulpes* | Vorpommern-Rügen | 106 | negative | - | - | - | negative | negative | negative |
| 2024BVD16411 | Carnivore | Raccoon dog | *Nyctereutes procyonoides* | Vorpommern-Rügen | not taken | not taken | not taken | not taken | - | negative | negative | negative |
| 2024BVD16412 | Carnivore | Raccoon | *Procyon lotor* | Vorpommern-Rügen | 107 | negative | - | - | - | negative | negative | negative |
| 2024BVD16413 | Carnivore | Raccoon | *Procyon lotor* | Vorpommern-Rügen | 109 | negative | - | - | - | negative | negative | negative |
| 2024BVD16414 | Carnivore | Raccoon | *Procyon lotor* | Vorpommern-Rügen | 15 | positive | 14 | positive | - | negative | negative | negative |
| 2024BVD16415 | Carnivore | Red fox | *Vulpes vulpes* | Vorpommern-Rügen | 103 | negative | - | - | - | negative | negative | negative |
| 2024BVD16416 | Carnivore | Raccoon dog | *Nyctereutes procyonoides* | Vorpommern-Rügen | 42 | positive | 6 | positive | - | negative | negative | negative |
| 2024BVD16417 | Carnivore | Raccoon dog | *Nyctereutes procyonoides* | Vorpommern-Rügen | 84 | negative | - | - | - | negative | negative | negative |
| 2024BVD16418 | Carnivore | Raccoon dog | *Nyctereutes procyonoides* | Vorpommern-Rügen | 38 | positive | 6 | positive | - | negative | negative | negative |

| | | | | | | | | | | | | |
|---|---|---|---|---|---|---|---|---|---|---|---|---|
| 2024BVD16419 | Carnivore | Raccoon dog | *Nyctereutes procyonoides* | Vorpommern-Rügen | 100 | negative | - | - | - | negative | negative | negative |
| 2024BVD16420 | Carnivore | Red fox | *Vulpes vulpes* | Vorpommern-Rügen | not taken | not taken | not taken | not taken | - | negative | negative | negative |
| 2024BVD16421 | Carnivore | Raccoon dog | *Nyctereutes procyonoides* | Vorpommern-Rügen | 8 | positive | 5 | positive | - | negative | negative | negative |
| 2024BVD16422 | Carnivore | Raccoon dog | *Nyctereutes procyonoides* | Vorpommern-Rügen | 8 | positive | 5 | positive | - | negative | negative | negative |
| 2024BVD16423 | Carnivore | Raccoon dog | *Nyctereutes procyonoides* | Vorpommern-Rügen | 89 | negative | - | - | - | negative | negative | negative |
| 2024BVD16424 | Carnivore | Raccoon dog | *Nyctereutes procyonoides* | Vorpommern-Rügen | 100 | negative | - | - | - | negative | negative | negative |
| 2024BVD16425 | Carnivore | Red fox | *Vulpes vulpes* | Vorpommern-Rügen | 105 | negative | - | - | - | negative | negative | negative |
| 2024BVD16426 | Carnivore | Red fox | *Vulpes vulpes* | Vorpommern-Rügen | not taken | not taken | not taken | not taken | - | negative | negative | negative |
| 2024BVD16427 | Carnivore | Raccoon dog | *Nyctereutes procyonoides* | Vorpommern-Rügen | 16 | positive | 7 | positive | - | negative | negative | negative |
| 2024BVD16428 | Carnivore | Red fox | *Vulpes vulpes* | Vorpommern-Rügen | 102 | negative | - | - | - | negative | negative | negative |
| 2024BVD16429 | Carnivore | Red fox | *Vulpes vulpes* | Vorpommern-Rügen | 91 | negative | - | - | - | negative | negative | negative |
| 2024BVD16430 | Carnivore | Raccoon dog | *Nyctereutes procyonoides* | Vorpommern-Rügen | 97 | negative | - | - | - | negative | negative | negative |
| 2024BVD16431 | Carnivore | Raccoon dog | *Nyctereutes procyonoides* | Vorpommern-Rügen | 73 | negative | - | - | - | negative | negative | negative |
| 2024BVD16432 | Carnivore | Raccoon | *Procyon lotor* | Vorpommern-Rügen | 98 | negative | - | - | - | negative | negative | negative |
| 2024BVD16433 | Carnivore | Raccoon | *Procyon lotor* | Vorpommern-Rügen | 103 | negative | - | - | - | negative | negative | negative |
| 2024BVD16434 | Carnivore | Raccoon | *Procyon lotor* | Vorpommern-Rügen | 105 | negative | - | - | - | negative | negative | negative |
| 2024BVD16435 | Carnivore | Red fox | *Vulpes vulpes* | Vorpommern-Rügen | 23 | positive | 22 | positive | - | negative | negative | negative |
| 2024BVD16436 | Carnivore | Raccoon | *Procyon lotor* | Vorpommern-Rügen | 109 | negative | - | - | - | negative | negative | negative |
| 2024BVD16437 | Carnivore | Red fox | *Vulpes vulpes* | Vorpommern-Rügen | 104 | negative | - | - | - | negative | negative | negative |

| | | | | | | | | | | | | |
|---|---|---|---|---|---|---|---|---|---|---|---|---|
| 2024BVD16438 | Carnivore | Raccoon | *Procyon lotor* | Vorpommern-Rügen | 105 | negative | - | - | - | not taken | negative | not taken |
| 2024BVD16439 | Carnivore | Raccoon dog | *Nyctereutes procyonoides* | Vorpommern-Rügen | 102 | negative | - | - | - | negative | negative | negative |
| 2024BVD16440 | Carnivore | Raccoon | *Procyon lotor* | Vorpommern-Rügen | 4 | positive | 4 | positive | - | negative | negative | negative |
| 2024BVD16441 | Carnivore | Raccoon dog | *Nyctereutes procyonoides* | Vorpommern-Rügen | 96 | negative | - | - | - | negative | negative | negative |
| 2024BVD16981 | Carnivore | Raccoon | *Procyon lotor* | Vorpommern-Rügen | 103 | negative | - | - | - | negative | negative | negative |
| 2024BVD16982 | Carnivore | Red fox | *Vulpes vulpes* | Vorpommern-Rügen | 79 | negative | - | - | - | negative | negative | negative |
| 2024BVD16983 | Carnivore | Red fox | *Vulpes vulpes* | Vorpommern-Rügen | 103 | negative | - | - | - | negative | negative | negative |
| 2024BVD16984 | Carnivore | Raccoon dog | *Nyctereutes procyonoides* | Vorpommern-Rügen | 106 | negative | - | - | - | negative | negative | negative |
| 2024BVD16985 | Carnivore | Red fox | *Vulpes vulpes* | Vorpommern-Rügen | 107 | negative | - | - | - | negative | negative | negative |
| 2024BVD16986 | Carnivore | Red fox | *Vulpes vulpes* | Vorpommern-Rügen | 96 | negative | - | - | - | negative | negative | negative |
| 2024BVD16987 | Carnivore | Red fox | *Vulpes vulpes* | Vorpommern-Rügen | 89 | negative | - | - | - | negative | negative | negative |
| 2024BVD16988 | Carnivore | Red fox | *Vulpes vulpes* | Vorpommern-Rügen | 98 | negative | - | - | - | negative | negative | negative |
| 2024BVD16989 | Carnivore | Red fox | *Vulpes vulpes* | Vorpommern-Rügen | 52 | negative | - | - | - | negative | negative | negative |
| 2024BVD16990 | Carnivore | Red fox | *Vulpes vulpes* | Vorpommern-Rügen | 68 | negative | - | - | - | negative | negative | negative |
| 2024BVD16991 | Carnivore | Raccoon dog | *Nyctereutes procyonoides* | Vorpommern-Rügen | 106 | negative | - | - | - | negative | negative | negative |
| 2024BVD16992 | Carnivore | Raccoon dog | *Nyctereutes procyonoides* | Vorpommern-Rügen | 110 | negative | - | - | - | negative | negative | negative |
| 2024BVD16993 | Carnivore | Red fox | *Vulpes vulpes* | Vorpommern-Rügen | 65 | negative | - | - | - | negative | negative | negative |
| 2024BVD16994 | Carnivore | Raccoon dog | *Nyctereutes procyonoides* | Vorpommern-Rügen | not taken | not taken | not taken | not taken | - | negative | negative | negative |
| 2024BVD16995 | Carnivore | Raccoon dog | *Nyctereutes procyonoides* | Vorpommern-Rügen | 96 | negative | - | - | - | negative | negative | negative |

| | | | | | | | | | | | | |
|---|---|---|---|---|---|---|---|---|---|---|---|---|
| 2024BVD16996 | Carnivore | Raccoon dog | *Nyctereutes procyonoides* | Vorpommern-Rügen | not taken | not taken | not taken | not taken | - | negative | negative | negative |
| 2024BVD16997 | Carnivore | Red fox | *Vulpes vulpes* | Vorpommern-Rügen | 104 | negative | - | - | - | negative | negative | negative |
| 2024BVD16998 | Carnivore | Raccoon | *Procyon lotor* | Vorpommern-Rügen | 109 | negative | - | - | - | negative | negative | negative |
| 2024BVD16999 | Carnivore | Raccoon | *Procyon lotor* | Vorpommern-Rügen | 105 | negative | - | - | - | negative | negative | negative |
| 2024BVD17000 | Carnivore | Raccoon | *Procyon lotor* | Vorpommern-Rügen | 105 | negative | - | - | - | negative | negative | not taken |
| 2024BVD17001 | Carnivore | Red fox | *Vulpes vulpes* | Vorpommern-Rügen | 101 | negative | - | - | - | negative | negative | negative |
| 2024BVD17002 | Carnivore | Raccoon | *Procyon lotor* | Vorpommern-Rügen | 105 | negative | - | - | - | negative | negative | negative |
| 2024BVD17003 | Carnivore | Raccoon | *Procyon lotor* | Vorpommern-Rügen | 105 | negative | - | - | - | negative | negative | negative |
| 2024BVD17004 | Carnivore | Red fox | *Vulpes vulpes* | Vorpommern-Rügen | 106 | negative | - | - | - | negative | negative | negative |
| 2024BVD17005 | Carnivore | Red fox | *Vulpes vulpes* | Vorpommern-Rügen | 94 | negative | - | - | - | negative | negative | negative |
| 2024BVD17006 | Carnivore | Red fox | *Vulpes vulpes* | Vorpommern-Rügen | 109 | negative | - | - | - | negative | negative | negative |
| 2024BVD17007 | Carnivore | Raccoon dog | *Nyctereutes procyonoides* | Vorpommern-Rügen | 103 | negative | - | - | - | negative | negative | negative |
| 2024BVD17008 | Carnivore | Raccoon dog | *Nyctereutes procyonoides* | Vorpommern-Rügen | 67 | negative | - | - | - | negative | negative | negative |
| 2024BVD17009 | Carnivore | Red fox | *Vulpes vulpes* | Vorpommern-Rügen | 97 | negative | - | - | - | negative | negative | negative |
| 2024BVD17010 | Carnivore | Raccoon | *Procyon lotor* | Vorpommern-Rügen | 109 | negative | - | - | - | negative | negative | negative |
| 2024BVD17011 | Carnivore | Raccoon dog | *Nyctereutes procyonoides* | Vorpommern-Rügen | 92 | negative | - | - | - | negative | negative | negative |
| 2024BVD17012 | Carnivore | Red fox | *Vulpes vulpes* | Vorpommern-Rügen | 101 | negative | - | - | - | negative | negative | negative |
| 2024BVD17013 | Carnivore | Raccoon dog | *Nyctereutes procyonoides* | Vorpommern-Rügen | 95 | negative | - | - | - | negative | negative | negative |
| 2024BVD17014 | Carnivore | Raccoon | *Procyon lotor* | Vorpommern-Rügen | 94 | negative | - | - | - | negative | negative | negative |

| | | | | | | | | | | | | |
|---|---|---|---|---|---|---|---|---|---|---|---|---|
| 2024BVD17015 | Carnivore | Red fox | *Vulpes vulpes* | Vorpommern-Rügen | 99 | negative | - | - | - | negative | negative | negative |
| 2024BVD17294 | Carnivore | Red fox | *Vulpes vulpes* | Vorpommern-Rügen | 64 | negative | - | - | - | negative | negative | negative |
| 2024BVD17295 | Carnivore | Red fox | *Vulpes vulpes* | Vorpommern-Rügen | 99 | negative | - | - | - | negative | negative | negative |
| 2024BVD17296 | Carnivore | Red fox | *Vulpes vulpes* | Vorpommern-Rügen | 46 | undetermined | not available | not available | - | negative | not taken | negative |
| 2024BVD17297 | Carnivore | Raccoon | *Procyon lotor* | Vorpommern-Rügen | 98 | negative | - | - | - | negative | negative | negative |
| 2024BVD17298 | Carnivore | Red fox | *Vulpes vulpes* | Vorpommern-Rügen | 66 | negative | - | - | - | negative | negative | negative |
| 2024BVD17299 | Carnivore | Raccoon dog | *Nyctereutes procyonoides* | Vorpommern-Rügen | 90 | negative | - | - | - | negative | negative | negative |
| 2024BVD17300 | Carnivore | Raccoon dog | *Nyctereutes procyonoides* | Vorpommern-Rügen | 101 | negative | - | - | - | negative | negative | negative |
| 2024BVD17301 | Carnivore | Red fox | *Vulpes vulpes* | Vorpommern-Rügen | not taken | not taken | not taken | not taken | - | negative | negative | negative |
| 2024BVD17302 | Carnivore | Red fox | *Vulpes vulpes* | Vorpommern-Rügen | not taken | not taken | not taken | not taken | - | negative | negative | negative |
| 2024BVD17303 | Carnivore | Red fox | *Vulpes vulpes* | Vorpommern-Rügen | 74 | negative | - | - | - | negative | negative | negative |
| 2024BVD17304 | Carnivore | Raccoon dog | *Nyctereutes procyonoides* | Vorpommern-Rügen | 81 | negative | - | - | - | negative | not taken | negative |
| 2024BVD17305 | Carnivore | Red fox | *Vulpes vulpes* | Vorpommern-Rügen | 38 | positive | 8 | positive | - | negative | negative | negative |
| 2024BVD17306 | Carnivore | Red fox | *Vulpes vulpes* | Vorpommern-Rügen | 98 | negative | - | - | - | negative | negative | negative |
| 2024BVD17307 | Carnivore | Raccoon | *Procyon lotor* | Vorpommern-Rügen | 98 | negative | - | - | - | negative | negative | negative |
| 2024BVD17308 | Carnivore | Red fox | *Vulpes vulpes* | Vorpommern-Rügen | 99 | negative | - | - | - | negative | negative | negative |
| 2024BVD17309 | Carnivore | Raccoon dog | *Nyctereutes procyonoides* | Vorpommern-Rügen | 98 | negative | - | - | - | negative | negative | not taken |
| 2024BVD17310 | Carnivore | Raccoon dog | *Nyctereutes procyonoides* | Vorpommern-Rügen | not taken | not taken | not taken | not taken | - | negative | not taken | not taken |
| 2024BVD17311 | Carnivore | Red fox | *Vulpes vulpes* | Vorpommern-Rügen | 96 | negative | - | - | - | negative | negative | negative |

| | | | | | | | | | | | | |
|---|---|---|---|---|---|---|---|---|---|---|---|---|
| 2024BVD17312 | Carnivore | Red fox | *Vulpes vulpes* | Vorpommern-Rügen | 103 | negative | - | - | - | negative | negative | negative |
| 2024BVD17313 | Carnivore | Red fox | *Vulpes vulpes* | Vorpommern-Rügen | 84 | negative | - | - | - | negative | negative | negative |
| 2024BVD17314 | Carnivore | Red fox | *Vulpes vulpes* | Vorpommern-Rügen | 88 | negative | - | - | - | negative | negative | negative |
| 2024BVD17315 | Carnivore | Red fox | *Vulpes vulpes* | Vorpommern-Rügen | 95 | negative | - | - | - | negative | negative | negative |
| 2024BVD17316 | Carnivore | Raccoon dog | *Nyctereutes procyonoides* | Vorpommern-Rügen | 70 | negative | - | - | - | negative | negative | negative |
| 2024BVD17317 | Carnivore | Raccoon dog | *Nyctereutes procyonoides* | Vorpommern-Rügen | 88 | negative | - | - | - | negative | negative | negative |
| 2024BVD17318 | Carnivore | European badger | *Meles meles* | Vorpommern-Rügen | 55 | negative | - | - | - | negative | negative | not taken |
| 2024BVD17319 | Carnivore | Red fox | *Vulpes vulpes* | Vorpommern-Rügen | 35 | positive | 30 | positive | - | negative | negative | negative |
| 2024BVD17320 | Carnivore | Red fox | *Vulpes vulpes* | Vorpommern-Rügen | not taken | not taken | not taken | not taken | - | negative | negative | negative |
| 2024BVD17321 | Carnivore | Raccoon | *Procyon lotor* | Vorpommern-Rügen | 26 | positive | 7 | positive | - | negative | negative | negative |
| 2024BVD17322 | Carnivore | Red fox | *Vulpes vulpes* | Vorpommern-Rügen | 63 | negative | - | - | - | negative | negative | negative |
| 2024BVD17323 | Carnivore | Raccoon dog | *Nyctereutes procyonoides* | Vorpommern-Rügen | 50 | undetermined | 23 | positive | - | negative | negative | not taken |
| 2024BVD17324 | Carnivore | Red fox | *Vulpes vulpes* | Vorpommern-Rügen | 71 | negative | - | - | - | negative | negative | negative |
| 2024BVD17325 | Carnivore | Raccoon dog | *Nyctereutes procyonoides* | Vorpommern-Rügen | not taken | not taken | not taken | not taken | - | negative | negative | negative |
| 2024BVD17326 | Carnivore | Raccoon dog | *Nyctereutes procyonoides* | Vorpommern-Rügen | not taken | not taken | not taken | not taken | - | negative | negative | negative |
| 2024BVD17327 | Carnivore | Red fox | *Vulpes vulpes* | Vorpommern-Rügen | 20 | positive | 30 | positive | - | negative | negative | negative |
| 2024BVD17328 | Carnivore | Red fox | *Vulpes vulpes* | Vorpommern-Rügen | not taken | not taken | not taken | not taken | - | negative | negative | negative |
| 2024BVD17329 | Carnivore | Red fox | *Vulpes vulpes* | Vorpommern-Rügen | 69 | negative | - | - | - | negative | negative | negative |
| 2024BVD17330 | Carnivore | Raccoon | *Procyon lotor* | Vorpommern-Rügen | 16 | positive | 30 | positive | - | negative | negative | negative |

| | | | | | | | | | | | | |
|---|---|---|---|---|---|---|---|---|---|---|---|---|
| 2024BVD17331 | Carnivore | Raccoon dog | *Nyctereutes procyonoides* | Vorpommern-Rügen | not taken | not taken | not taken | not taken | - | negative | negative | negative |
| 2024BVD17332 | Carnivore | Raccoon dog | *Nyctereutes procyonoides* | Vorpommern-Rügen | not taken | not taken | not taken | not taken | - | negative | negative | negative |
| 2024BVD17333 | Carnivore | Raccoon dog | *Nyctereutes procyonoides* | Vorpommern-Rügen | 106 | negative | - | - | - | negative | negative | negative |
| 2024BVD17334 | Carnivore | Raccoon dog | *Nyctereutes procyonoides* | Vorpommern-Rügen | 79 | negative | - | - | - | negative | negative | negative |
| 2024BVD17807 | Carnivore | Red fox | *Vulpes vulpes* | Vorpommern-Rügen | 73 | negative | - | - | - | negative | negative | negative |
| 2024BVD17808 | Carnivore | Red fox | *Vulpes vulpes* | Vorpommern-Rügen | 28 | positive | 5 | positive | - | negative | negative | negative |
| 2024BVD17809 | Carnivore | Raccoon dog | *Nyctereutes procyonoides* | Vorpommern-Rügen | 62 | negative | - | - | - | negative | negative | negative |
| 2024BVD17810 | Carnivore | Raccoon dog | *Nyctereutes procyonoides* | Vorpommern-Rügen | 65 | negative | - | - | - | negative | negative | negative |
| 2024BVD17811 | Carnivore | Red fox | *Vulpes vulpes* | Vorpommern-Rügen | 66 | negative | - | - | - | negative | negative | negative |
| 2024BVD17812 | Carnivore | Red fox | *Vulpes vulpes* | Vorpommern-Rügen | 42 | positive | 70 | negative | - | negative | negative | negative |
| 2024BVD17813 | Carnivore | Red fox | *Vulpes vulpes* | Vorpommern-Rügen | 89 | negative | - | - | - | negative | negative | negative |
| 2024BVD17814 | Carnivore | Red fox | *Vulpes vulpes* | Vorpommern-Rügen | not taken | not taken | not taken | not taken | - | negative | negative | negative |
| 2024BVD17815 | Carnivore | Raccoon | *Procyon lotor* | Vorpommern-Rügen | 50 | undetermined | 43 | positive | - | negative | negative | negative |
| 2024BVD17816 | Carnivore | Red fox | *Vulpes vulpes* | Vorpommern-Rügen | 46 | undetermined | 50 | positive | - | negative | negative | negative |
| 2024BVD17817 | Carnivore | Raccoon | *Procyon lotor* | Vorpommern-Rügen | 70 | negative | - | - | - | negative | negative | negative |
| 2024BVD17818 | Carnivore | Red fox | *Vulpes vulpes* | Vorpommern-Rügen | 60 | negative | - | - | - | negative | negative | negative |
| 2024BVD17819 | Carnivore | Red fox | *Vulpes vulpes* | Vorpommern-Rügen | 69 | negative | - | - | - | negative | negative | negative |
| 2024BVD17820 | Carnivore | Raccoon | *Procyon lotor* | Vorpommern-Rügen | 73 | negative | - | - | - | negative | negative | negative |
| 2024BVD17821 | Carnivore | Red fox | *Vulpes vulpes* | Vorpommern-Rügen | 16 | positive | 16 | positive | - | negative | negative | negative |

| | | | | | | | | | | | | |
|---|---|---|---|---|---|---|---|---|---|---|---|---|
| 2024BVD17822 | Carnivore | Fox | *Vulpes sp.* | Vorpommern-Rügen | 17 | positive | 18 | positive | - | negative | negative | negative |
| 2024BVD17823 | Carnivore | Fox | *Vulpes sp.* | Vorpommern-Rügen | 12 | positive | 8 | positive | - | negative | negative | negative |
| 2024BVD17824 | Carnivore | Fox | *Vulpes sp.* | Vorpommern-Rügen | 66 | negative | - | - | - | negative | negative | negative |
| 2025BVD00300 | Carnivore | Fox | *Vulpes sp.* | Vorpommern-Rügen | 45 | undetermined | 25 | positive | - | negative | negative | negative |
| 2025BVD00301 | Carnivore | Raccoon dog | *Nyctereutes procyonoides* | Vorpommern-Rügen | 12 | positive | 17 | positive | - | negative | negative | negative |
| 2025BVD00302 | Carnivore | Fox | *Vulpes sp.* | Vorpommern-Rügen | 23 | positive | 61 | negative | <1:20 | negative | negative | negative |
| 2025BVD00303 | Carnivore | Fox | *Vulpes sp.* | Vorpommern-Rügen | 57 | negative | - | - | - | negative | negative | negative |
| 2025BVD00304 | Carnivore | Raccoon | *Procyon lotor* | Vorpommern-Rügen | 71 | negative | - | - | - | negative | negative | negative |
| 2025BVD00305 | Carnivore | Raccoon dog | *Nyctereutes procyonoides* | Vorpommern-Rügen | 72 | negative | - | - | - | negative | negative | negative |
| 2025BVD00306 | Carnivore | Fox | *Vulpes sp.* | Vorpommern-Rügen | 57 | negative | - | - | - | negative | negative | negative |
| 2025BVD00307 | Carnivore | Fox | *Vulpes sp.* | Vorpommern-Rügen | 63 | negative | - | - | - | negative | negative | not taken |
| 2025BVD00308 | Carnivore | Fox | *Vulpes sp.* | Vorpommern-Rügen | 32 | positive | 52 | undetermined | - | negative | negative | negative |
| 2025BVD00309 | Carnivore | Raccoon | *Procyon lotor* | Vorpommern-Rügen | 10 | positive | 7 | positive | - | negative | negative | negative |
| 2025BVD00310 | Carnivore | Fox | *Vulpes sp.* | Vorpommern-Rügen | 13 | positive | 5 | positive | - | negative | negative | not taken |
| 2025BVD00311 | Carnivore | Raccoon dog | *Nyctereutes procyonoides* | Vorpommern-Rügen | 12 | positive | 8 | positive | - | negative | negative | not taken |
| 2025BVD00312 | Carnivore | Fox | *Vulpes sp.* | Vorpommern-Rügen | 75 | negative | - | - | - | negative | negative | negative |
| 2025BVD00313 | Carnivore | Raccoon dog | *Nyctereutes procyonoides* | Vorpommern-Rügen | 81 | negative | - | - | - | negative | negative | not taken |
| 2025BVD00314 | Carnivore | Fox | *Vulpes sp.* | Vorpommern-Rügen | 76 | negative | - | - | - | negative | negative | negative |
| 2025BVD00315 | Carnivore | Raccoon dog | *Nyctereutes procyonoides* | Vorpommern-Rügen | 70 | negative | - | - | - | negative | negative | negative |

| | | | | | | | | | | | | |
|---|---|---|---|---|---|---|---|---|---|---|---|---|
| 2025BVD00316 | Carnivore | Raccoon | *Procyon lotor* | Vorpommern-Rügen | 77 | negative | - | - | - | negative | negative | not taken |
| 2025BVD00317 | Carnivore | Raccoon | *Procyon lotor* | Vorpommern-Rügen | 60 | negative | - | - | - | negative | negative | negative |
| 2025BVD00318 | Carnivore | Raccoon dog | *Nyctereutes procyonoides* | Vorpommern-Rügen | 44 | positive | 75 | negative | - | negative | negative | negative |
| 2025BVD00319 | Carnivore | Fox | *Vulpes sp.* | Vorpommern-Rügen | 85 | negative | - | - | - | negative | negative | negative |
| 2025BVD00320 | Carnivore | Fox | *Vulpes sp.* | Vorpommern-Rügen | 74 | negative | - | - | - | negative | negative | negative |
| 2025BVD00321 | Carnivore | Raccoon dog | *Nyctereutes procyonoides* | Vorpommern-Rügen | 20 | positive | 7 | positive | - | negative | negative | negative |
| 2025BVD00322 | Carnivore | Fox | *Vulpes sp.* | Vorpommern-Rügen | 70 | negative | - | - | - | negative | negative | negative |
| 2025BVD00323 | Carnivore | Raccoon dog | *Nyctereutes procyonoides* | Vorpommern-Rügen | 20 | positive | 6 | positive | 1:60 | negative | negative | negative |
| 2025BVD00324 | Carnivore | Fox | *Vulpes sp.* | Vorpommern-Rügen | 53 | negative | - | - | - | negative | negative | negative |
| 2025BVD00325 | Carnivore | Fox | *Vulpes sp.* | Vorpommern-Rügen | 63 | negative | - | - | - | negative | negative | not taken |
| 2025BVD00326 | Carnivore | Fox | *Vulpes sp.* | Vorpommern-Rügen | not taken | not taken | not taken | not taken | - | negative | negative | negative |
| 2025BVD00327 | Carnivore | Fox | *Vulpes sp.* | Vorpommern-Rügen | 24 | positive | 28 | positive | - | negative | negative | negative |
| 2025BVD00328 | Carnivore | Fox | *Vulpes sp.* | Vorpommern-Rügen | 63 | negative | - | - | - | negative | negative | negative |
| 2025BVD00329 | Carnivore | Fox | *Vulpes sp.* | Vorpommern-Rügen | 58 | negative | - | - | - | negative | negative | negative |
| 2025BVD00330 | Carnivore | Fox | *Vulpes sp.* | Vorpommern-Rügen | 67 | negative | - | - | - | negative | negative | negative |
| 2025BVD00847 | Carnivore | Raccoon dog | *Nyctereutes procyonoides* | Vorpommern-Rügen | 73 | negative | - | - | - | negative | negative | negative |
| 2025BVD00848 | Carnivore | Fox | *Vulpes sp.* | Vorpommern-Rügen | 96 | negative | - | - | - | negative | negative | negative |
| 2025BVD00849 | Carnivore | Fox | *Vulpes sp.* | Vorpommern-Rügen | not taken | not taken | not taken | not taken | - | negative | negative | negative |
| 2025BVD00850 | Carnivore | Fox | *Vulpes sp.* | Vorpommern-Rügen | 43 | positive | 75 | negative | - | negative | negative | negative |

| | | | | | | | | | | | | |
|---|---|---|---|---|---|---|---|---|---|---|---|---|
| 2025BVD00851 | Carnivore | Fox | *Vulpes sp.* | Vorpommern-Rügen | 52 | negative | - | - | - | negative | negative | negative |
| 2025BVD00852 | Carnivore | Raccoon | *Procyon lotor* | Vorpommern-Rügen | 33 | positive | 70 | negative | - | negative | negative | negative |
| 2025BVD00853 | Carnivore | Raccoon dog | *Nyctereutes procyonoides* | Vorpommern-Rügen | 83 | negative | - | - | - | negative | negative | negative |
| 2025BVD00854 | Carnivore | Fox | *Vulpes sp.* | Vorpommern-Rügen | 56 | negative | - | - | - | negative | negative | negative |
| 2025BVD00855 | Carnivore | Fox | *Vulpes sp.* | Vorpommern-Rügen | 59 | negative | - | - | - | negative | negative | negative |
| 2025BVD00856 | Carnivore | Fox | *Vulpes sp.* | Vorpommern-Rügen | 53 | negative | - | - | - | negative | negative | negative |
| 2025BVD00857 | Carnivore | Raccoon dog | *Nyctereutes procyonoides* | Vorpommern-Rügen | 38 | positive | 68 | negative | - | negative | negative | negative |
| 2025BVD00858 | Carnivore | Raccoon | *Procyon lotor* | Vorpommern-Rügen | 63 | negative | - | - | - | negative | negative | negative |
| 2025BVD00859 | Carnivore | Fox | *Vulpes sp.* | Vorpommern-Rügen | not taken | not taken | not taken | not taken | - | negative | negative | negative |
| 2025BVD00860 | Carnivore | Fox | *Vulpes sp.* | Vorpommern-Rügen | 29 | positive | 6 | positive | - | negative | negative | negative |
| 2025BVD00861 | Carnivore | Fox | *Vulpes sp.* | Vorpommern-Rügen | 72 | negative | - | - | - | negative | negative | negative |
| 2025BVD00862 | Carnivore | Fox | *Vulpes sp.* | Vorpommern-Rügen | 34 | positive | 8 | positive | - | negative | negative | negative |
| 2025BVD00863 | Carnivore | Fox | *Vulpes sp.* | Vorpommern-Rügen | 52 | negative | - | - | - | negative | negative | negative |
| 2025BVD00864 | Carnivore | Fox | *Vulpes sp.* | Vorpommern-Rügen | 91 | negative | - | - | - | negative | negative | negative |
| 2025BVD00865 | Carnivore | Fox | *Vulpes sp.* | Vorpommern-Rügen | 65 | negative | - | - | - | negative | negative | negative |
| 2025BVD00866 | Carnivore | Raccoon dog | *Nyctereutes procyonoides* | Vorpommern-Rügen | 57 | negative | - | - | - | negative | negative | negative |
| 2025BVD00867 | Carnivore | Raccoon dog | *Nyctereutes procyonoides* | Vorpommern-Rügen | 35 | positive | 23 | positive | - | negative | negative | negative |
| 2025BVD00868 | Carnivore | Raccoon dog | *Nyctereutes procyonoides* | Vorpommern-Rügen | 76 | negative | - | - | - | negative | not taken | negative |
| 2025BVD00869 | Carnivore | Fox | *Vulpes sp.* | Vorpommern-Rügen | 8 | positive | 6 | positive | - | negative | negative | negative |

| | | | | | | | | | | | | |
|---|---|---|---|---|---|---|---|---|---|---|---|---|
| 2025BVD00870 | Carnivore | Fox | *Vulpes sp.* | Vorpommern-Rügen | 66 | negative | - | - | - | negative | negative | negative |
| 2025BVD00871 | Carnivore | Raccoon | *Procyon lotor* | Vorpommern-Rügen | 63 | negative | - | - | - | negative | negative | negative |
| 2025BVD00872 | Carnivore | Fox | *Vulpes sp.* | Vorpommern-Rügen | 65 | negative | - | - | - | negative | negative | negative |
| 2025BVD00873 | Carnivore | Fox | *Vulpes sp.* | Vorpommern-Rügen | 61 | negative | - | - | - | negative | negative | negative |
| 2025BVD00874 | Carnivore | Fox | *Vulpes sp.* | Vorpommern-Rügen | 24 | positive | 24 | positive | - | negative | negative | negative |
| 2025BVD00875 | Carnivore | Fox | *Vulpes sp.* | Vorpommern-Rügen | 75 | negative | - | - | - | negative | negative | negative |
| 2025BVD00876 | Carnivore | Fox | *Vulpes sp.* | Vorpommern-Rügen | 38 | positive | 17 | positive | - | negative | negative | negative |
| 2025BVD00877 | Carnivore | Fox | *Vulpes sp.* | Vorpommern-Rügen | 66 | negative | - | - | - | negative | negative | negative |
| 2025BVD00878 | Carnivore | Raccoon dog | *Nyctereutes procyonoides* | Vorpommern-Rügen | 70 | negative | - | - | - | negative | negative | negative |

***Supplementary Table 2*** *Environmental variables tested either as proportion (variables ending with "prop") in a 2.5 km buffer zone or as a distance in 100 m steps (variables ending with "dist100") to the point geographic location where the animal carcass had been found.*

| Variable (abbreviation used in modelling) | Short description | Detailed description, including codes of the „Biotoptypen- und Nutzungstypenkartierung" |
|---|---|---|
| Forest (Forest_prop, Forest_dist100) | forest (< 4 ha) | B10 Forest (> 4 ha) B11 Deciduous forest B12 Mixed deciduous forest (< 10 % conifers) B13 Mixed deciduous forest (ratio of deciduous to coniferous trees 90/10 - 70/30) B14 Mixed forest (ratio of deciduous to coniferous trees 50/50) B15 Mixed coniferous forest (ratio of coniferous to deciduous trees 90/10 - 70/30) B16 Coniferous forest B17 Forest edge B18 Clear-cut B19 Glade/clearing |
| Shrubland (Shrubland_prop, Shrubland_dist100) | group of trees, hedge, bushes | B20 Group of trees, hedge, bushes B21 Field copse (0.5 - 4 ha) B22 Group of trees (< 0.5 ha) B23 Row of trees B24 Avenue B25 Dominant single tree B26 Hedge B27 Bushes, group of shrubs |
| Farmland (Farmland_prop, Farmland_dist100) | arable land, commercial horticulture | L20 Arable land, commercial horticulture L21 Arable land L22 Commercial horticulture L23 Tree nursery L24 Fruit growing |
| Grassland (Grassland_prop, Grassland_dist100) | Grassland | L11 Wet grassland L12 Fresh grassland bntk_f.doc Page 3 27.02.2013 L13 Seasonally wet grassland L14 Dry grassland L15 Salt grassland |
| Open areas (Openareas_prop, Openareas_dist100) | open space | S40 Open space S41 Park S42 Wildlife enclosure, zoo S43 Leisure park S44 Sports facility S45 Golf course S46 Campsite S47 Allotment gardens, holiday homes S48 Village green S49 Cemetery |
| Transportation (Trafficarea_prop, Trafficarea_dist100) | traffic area | S50 Traffic area S51 Path S52 Unpaved farm track S53 Paved farm track S54 Road S55 Motorway S56 Railway/Railway system S57 Airfield S58 Port facility S59 Parking lot |
| Watercourses (Watercourses_prop, Watercourses_dist100) | Rivers | W10 Flowing water W11 Source area W12 Stream < 3m W13 Ditch < 3m W14 River > 3m W15 Canal > 3m |

| | | |
|---|---|---|
| Excavation (Excavation_prop, Excavation_dist100) | excavation and embankment | R00 Excavation and backfill R10 Raw material extraction R11 Quarry/chalk quarry R12 Sand/gravel pit R13 Clay pit R14 Undifferentiated site excavation R20 Backfill R21 Landfill R22 Uncontrolled landfill R23 Scour field R24 Agricultural storage area (dung heap, pile, silo) |
| Heathland (Heathland_prop, Heathland_dist100) | heath | T10 Heide T11 Zwergstrauchheide T12 Ginsterheide T13 Krähenbeerheide |
| Dryland (Dryland_prop, Dryland_dist100) | dry grassland | T20 dry grassland T21 silicate dry grassland T22 calcareous dry grassland |
| | rocky outcrop | T30 Rocky outcrop T31 Inset outcrop rock T32 Stone pile and wall T33 Single boulder |
| Infrastructure (Infrastruct_prop, Infrastruct_dist100) | residential area | S10 Residential area S11 Closed development S12 Individual development S13 New development area, undifferentiated |
| | mixed area | S20 Mixed-use area S21 Urban mixed-use area S22 Rural mixed-use area S23 Single farmstead |
| | production facility | S30 Production facility S31 Commercial and industrial area S32 Animal production facility S33 Military facility |
| | hydraulic structure | S60 Hydraulic structure S61 Dike, dam S62 Groyne S63 Stone wall S64 Pumping station |
| | supply and disposal facilities | S70 Supply and disposal facilities S71 District heating pipeline S72 Sewage treatment plant S73 Slurry tank |
| Other waterbodies (Otherwater_prop, Otherwater_dist100) | small standing water body < 1 ha | W20 Small standing water body < 1 ha W21 Temporary small water body, pond or pool W22 Permanent small water body |
| | standing water > 1 ha | W30 Standing water > 1 ha W31 Shallow lake < 5m W32 Lake |

| | | |
|---|---|---|
| | moor and swamp | W40 Moor and swamp W41 Fens W42 Raised and transitional moor W43 Swamp |
| | coastal biotopes | W70 Coastal biotopes W71 Sand hook W72 Beach W73 Beach lake W74 Beach wall W75 White dune W76 Cliff W77 Cliff edge dune |
| Bodden coast (Baycoast_prop, Baycoast_dist100) | bodden (bay) | W60 Bodden (bay) W61 Open water [Bodden] W62 Marine boulder and stone bottom [Bodden] W63 Sandbank [Bodden] |
| Urban (Urban_prop, Urban_dist) | DLM data | mixed use, cemetery, industry and commerce, sports / leisure / recreation, residential area, square, road traffic, traffic structures, open-cast mine / pit / quarry, rail traffic, air traffic, spoil heap |
| Baltic sea (Balticsea_prop, Balticsea_dist) | this study | all remaining areas surrounding the island |

***Supplementary Table** 3 Results of the univariable logistic regression analysis using data on landcover and individual animals (for land cover data on buffer proportions [prop] or distance in 100 m [dist100] were used; data on age was stratified for adult and juvenile foxes). Results displayed were restricted to models with P values < 0.2.*

| **Model (AIC[a])** | **Variable** | **Esti-mate** | **Standard Error** | **z-value** | ***P*-value** | **Odds Ratio** | **2.5% CI[b]** | **97.5% CI[b]** | **Category *P*-value (< 0.1)** |
|---|---|---|---|---|---|---|---|---|---|
| 1 (269.12) | (Intercept) | -1.58 | 0.20 | -7.81 | 5.76E-15 | 0.21 | 0.14 | 0.30 | *** |
| | Baycoast_prop | 1.22 | 0.88 | 1.39 | 0.166 | 3.37 | 0.57 | 19.10 | - |
| 2 (268.55) | (Intercept) | -1.16 | 0.21 | -5.42 | 5.91E-08 | 0.31 | 0.20 | 0.47 | *** |
| | Otherwater_prop | -1.57 | 1.06 | -1.48 | 0.139 | 0.21 | 0.02 | 1.46 | - |
| 3 (265.17) | (Intercept) | -0.81 | 0.29 | -2.80 | 0.0052 | 0.45 | 0.25 | 0.78 | ** |
| | Shrubland_prop | -2.89 | 1.28 | -2.25 | 0.0245 | 0.06 | 0.00 | 0.60 | * |
| 4 (267.96) | (Intercept) | -1.17 | 0.20 | -5.85 | 4.93E-09 | 0.31 | 0.21 | 0.46 | *** |
| | Forest_prop | -0.28 | 0.17 | -1.61 | 0.107 | 0.76 | 0.52 | 1.03 | - |
| 5 (267.97) | (Intercept) | -1.61 | 0.20 | -8.07 | 6.98E-16 | 1.99E-01 | 0.13 | 0.29 | *** |
| | Openareas_prop | 19.54 | 10.97 | 1.78 | 0.075 | 3.07E+08 | 0.07 | 5.16E+17 | . |
| 6 (268.4) | (Intercept) | -1.27 | 0.17 | -7.29 | 3.20E-13 | 0.28 | 0.20 | 0.39 | *** |
| | Excavation_prop | -2.89 | 2.03 | -1.43 | 0.154 | 0.06 | 0.00 | 1.74 | - |
| 7 (268.49) | (Intercept) | -1.31 | 0.16 | -7.99 | 1.37E-15 | 0.27 | 1.95E-01 | 0.37 | *** |
| | Dryland_prop | -4.45 | 3.17 | -1.40 | 0.161 | 0.01 | 7.29E-06 | 2.55 | - |
| 8 (267.66) | (Intercept) | -1.10 | 0.22 | -5.01 | 5.36E-07 | 3.32E-01 | 2.13E-01 | 0.51 | *** |
| | Trafficarea_prop | -10.69 | 6.01 | -1.78 | 0.0755 | 2.29E-05 | 1.15E-10 | 2.18 | . |
| 9 (265.7) | (Intercept) | -0.87 | 0.28 | -3.05 | 0.00231 | 0.42 | 0.24 | 0.74 | ** |
| | Urban_prop | -1.16 | 0.56 | -2.06 | 0.03945 | 0.31 | 0.09 | 0.86 | * |
| 10 (264.86) | (Intercept) | -1.66 | 0.19 | -8.72 | <2e-16 | 1.90E-01 | 0.13 | 0.27 | *** |
| | Watercourses_prop | 31.65 | 12.57 | 2.52 | 0.0118 | 5.54E+13 | 917.99 | 6.49E+24 | * |
| 11 (269.26) | (Intercept) | -1.52 | 0.18 | -8.48 | <2e-16 | 0.22 | 0.15 | 0.31 | *** |
| | Greenland_dist100 | 0.06 | 0.05 | 1.31 | 0.19 | 1.06 | 0.97 | 1.18 | - |
| 12 (269.27) | (Intercept) | -1.51 | 0.17 | -8.66 | <2e-16 | 0.22 | 0.16 | 0.31 | *** |
| | Srubland_dist100 | 0.04 | 0.03 | 1.30 | 0.195 | 1.04 | 0.98 | 1.11 | - |

| | | | | | | | | | |
|---|---|---|---|---|---|---|---|---|---|
| 13 (269.24) | (Intercept) | -1.62 | 0.23 | -7.17 | 7.40E-13 | 0.20 | 0.13 | 0.31 | *** |
| | Urban_dist100 | 0.06 | 0.04 | 1.36 | 0.175 | 1.06 | 0.97 | 1.15 | - |
| 14 (**263.91**) | (Intercept) | -0.99 | 0.21 | -4.65 | 3.34E-06 | 0.37 | 0.24 | 0.56 | *** |
| | Baycoast_dist100 | -0.02 | 0.01 | -2.49 | 0.0128 | 0.98 | 0.96 | 1.00 | * |
| 15 (**262.91**) | (Intercept) | -0.93 | 0.22 | -4.23 | 2.37E-05 | 0.39 | 0.25 | 0.60 | *** |
| | Balticsea_dist100 | -0.03 | 0.01 | -2.60 | 0.00935 | 0.97 | 0.94 | 0.99 | ** |
| 16 (**264.26**) | (Intercept) | -0.84 | 0.26 | -3.23 | 0.00126 | 0.43 | 0.26 | 0.71 | ** |
| | Watercourses_dist100 | -0.05 | 0.02 | -2.43 | 0.01527 | 0.96 | 0.92 | 0.99 | * |
| 17 (265.37) | (Intercept) | -1.18 | 0.17 | -6.77 | 1.28E-11 | 0.31 | 0.22 | 0.43 | *** |
| | Age-Adult (Ref.) | - | - | - | - | - | - | - | - |
| | Age-Juvenil | -0.85 | 0.38 | -2.24 | 0.0253 | 0.43 | 0.19 | 0.87 | * |

[a] Akaike Information Index, [b]2.5% and 97.5% Confidence interval of Odds ratio

***Supplementary Table 4*** *Final multivariable logistic regression model using landcover variables (buffer proportions (prop), distance in 100 m (dist100) or age (Adult vs juvenile).*

| **Model (AIC[a], Pseudo R2)** | **Variable** | **Esti-mate** | **Standard Error** | **z-value** | ***P*-value** | **Odds Ratio** | **2.5% CI[b]** | **97.5% CI[b]** | **Category *P*-value (< 0.1)** |
|---|---|---|---|---|---|---|---|---|---|
| 18 (248.75) | (Intercept) | 0.37 | 0.45 | 0.81 | 0.42 | 1.44 | 0.60 | 3.60 | - |
| | Shrubland_prop | -4.11 | 1.54 | -2.66 | 0.01 | 0.02 | 0.00 | 0.29 | ** |
| | Shrubland_dist100 | 0.11 | 0.04 | 2.67 | 0.01 | 1.11 | 1.02 | 1.20 | ** |
| | Age-Adult (Ref.) | - | - | - | - | - | - | - | - |
| | Age-Juvenil | -0.55 | 0.40 | -1.38 | 0.17 | 0.58 | 0.25 | 1.23 | |
| | Watercourses_dist100 | -0.09 | 0.02 | -3.89 | 0.00 | 0.92 | 0.88 | 0.96 | *** |

[a] Akaike Information Index, [b]2.5% and 97.5% Confidence interval of Odds ratio
